# Temporal tuning of actin nucleation by the repurposed DNA repair complex RAD51/fignl1 encodes distinct axon guidance decisions

**DOI:** 10.64898/2026.10.17.759799

**Authors:** Samya Zerkoune, Yvrick Zagar, Julie Delaroche, Julia Danicka, Juliette Vougny, Sinthuya Uthayasuthan, Julien Aureille, Fiona Roche, Rhanna Haanjtes, Claudia Gomez-Bravo, Sarah Baudet, Eric Denarier, Akbar Zainu, Philippe Bun, Xavier Guillonneau, Anne Debant, Frédéric Baudat, Xavier Nicol, Nicolas Pietrancosta, Jérôme Boudeau, Annie Andrieux, Isabelle Arnal, Adrien Antkowiak, Fassier Coralie

## Abstract

Axon guidance is essential for neural circuit assembly and function. While much of the repertoire of guidance cues and receptors is known, its limited diversity cannot fully explain the remarkable complexity of the brain connectome. How temporally ordered, context-dependent intrinsic programs shape the diversity of axonal responses to guidance signals remains poorly understood. We reveal how non-canonical Wnt5a signalling repurposes the DNA repair complex RAD51/fignl1 as a downstream integrator to control retinal ganglion cell axon decision-making at the midline. We demonstrate that differential RAD51/fignl1 expression and interaction dynamics between ipsilateral and contralateral axons tunes the magnitude and temporal dynamics of F-actin nucleation, thereby encoding their trajectory choice. Our findings establish temporal control of cytoskeletal dynamics as a key determinant of axon decision-making and thereby a fundamental layer of circuit wiring. They provide a potential mechanistic framework linking mutations in the DNA repair gene RAD51 to neurodevelopmental disorders with midline guidance defects.

## Introduction

The nervous system governs vital functions and complex behaviours, enabling animals and humans to interact with their environment through sensory and motor adaptations. It is an intricate network in which precise connectivity between neurons is crucial for proper function. Establishing such precise neuronal connections during development requires axons to navigate complex environments, integrating multiple guidance cues at key guidance choice points to reach their specific targets. Midline navigation serves as a critical decision point where axons choose either contralateral or ipsilateral trajectories. This binary choice is essential for establishing commissural connectivity and wiring bilateral circuits.^1–3^ Defects in axon guidance at the midline contribute to neurodevelopmental disorders^4–8^ and limit optic nerve regeneration.^9–11^

Mutations in the DNA recombinase RAD51 cause a neurodevelopmental disorder associated with axon misrouting at the midline, highlighting a paradoxical role for RAD51 in wiring neuronal circuits.^12, 8^ RAD51 is classically recognized for its nuclear function as a pivotal player in DNA double-strand break repair through homologous recombination. RAD51 forms nucleoprotein filaments on single-stranded DNA, thereby mediating homology search and strand invasion.^13–16^ Its activity is tightly regulated by the AAA+ ATPase fidgetin-like 1 (fignl1), which disassembles RAD51 filaments via a scaffolding mechanism, thereby preventing aberrant recombination and preserving genome stability.^16–23^ Together, these canonical nuclear functions establish the RAD51/fignl1 complex as a central safeguard of genome integrity. Yet, whether this molecular machinery operates outside the nucleus to influence other cellular processes, such as axon guidance at the midline choice point, remains unexplored. The implication of fignl1 in microtubule-dependent motor axon steering during zebrafish development^24,25^ raises the intriguing possibility of an unrecognized role for the RAD51/fignl1 complex in cytoskeletal control of axon guidance.

Axon guidance at the midline is governed by a finely balanced and temporally controlled interplay of attractive and repulsive cues, including classical guidance molecules (netrins, ephrins, Slits)^26–31^ and morphogens (Wnt5a, Shh).^28, 32^ Yet, our understanding of the intrinsic mechanisms that tune axon responsiveness to these signals remains incomplete. Known intrinsic modulators include alternative splicing/translation,^33–36^ proteolytic processing^37,38^ and membrane trafficking of guidance receptors,^39–41^ as well as local mRNA translation.^42,43^ Cytoskeletal remodelling has also emerged as a key driving force of growth cone steering^44–48^. Yet, how guidance signals are integrated and translated into coordinated cytoskeletal remodelling events that drive growth cone mechanical behaviours remains largely unknown. Notably, whether cytoskeletal dynamics merely execute decisions imposed by guidance signals or can themselves act as instructive determinants of axon trajectory choice remains unknown. Here, by integrating experimental approaches spanning molecular to circuit scales with advanced imaging techniques, we identify the DNA repair complex RAD51/fignl1 as an unexpected cytoplasmic integrator of non-canonical Wnt5a signalling controlling axon decision-making at the midline. We reveal PKC-mediated phosphorylation of RAD51-scaffolded fignl1 at Ser329 as a molecular switch that sensitizes axons to Wnt5a by suppressing fignl1’s microtubule-depolymerizing activity while potentiating its previously unrecognised basal inhibitory function on actin nucleation. Notably, we demonstrate that differential RAD51 and fignl1 expression levels and interaction dynamics between ipsilateral and contralateral axons tune the magnitude and duration of actin nucleation inhibition, thereby encoding their distinct trajectory choices. Our findings establish temporal control of cytoskeletal dynamics as a key determinant of axon decision-making and a fundamental layer of neuronal circuit wiring. They further provide a potential mechanistic framework for understanding the long-standing paradox linking mutations in a major DNA repair gene to neurodevelopmental disorders associated with midline axon guidance defects.

## Results

### Fignl1 is required for retinal axon midline crossing

Building on our previous work identifying fignl1 as a microtubule-depolymerizing ATPase essential for axon steering at guidance choice points in zebrafish,^24–25^ we asked whether this cytoskeletal regulator plays a role in axon guidance at the midline in the mammalian brain. We focused on the visual system, where retinal ganglion cell (RGC) axons make a binary decision to cross or avoid the midline, providing a tractable and experimentally accessible model to examine axon decision-making at this critical choice point. We found that mouse fignl1 is expressed in developing RGCs during the peak period of axon guidance at the optic chiasm (E16; Figures 1A and 1B and Figure S1A-S1D) and exhibits an ATPase-dependant microtubule-depolymerizing activity in mammalian cells (*e.g.* COS-7) – similar to its zebrafish counterpart.^24^ This activity relies on its RAD51 binding domain (RBD) and microtubule-binding domain (MTBD; Figures S1E-S1H). To test whether fignl1 regulates RGC axon guidance at the chiasm midline, *in utero* electroporation was performed on E14.5 embryos to express *fignl1* or control shRNA together with tdTomato in a single retina (electroporation of the central retina to target contralateral RGCs). Trajectories of electroporated axons were visualized at E18.5 by *in toto* immunolabelling combined with tissue clearing and 3D light-sheet imaging (Figure 1C and Video S1). Fignl1 knockdown impairs RGC axon pathfinding at the chiasm, resulting in an excess of ipsilateral projections (Figures 1D-1F and Video S1), suggesting a critical role for fignl1 in retinal axon midline crossing.

**Figure 1:**
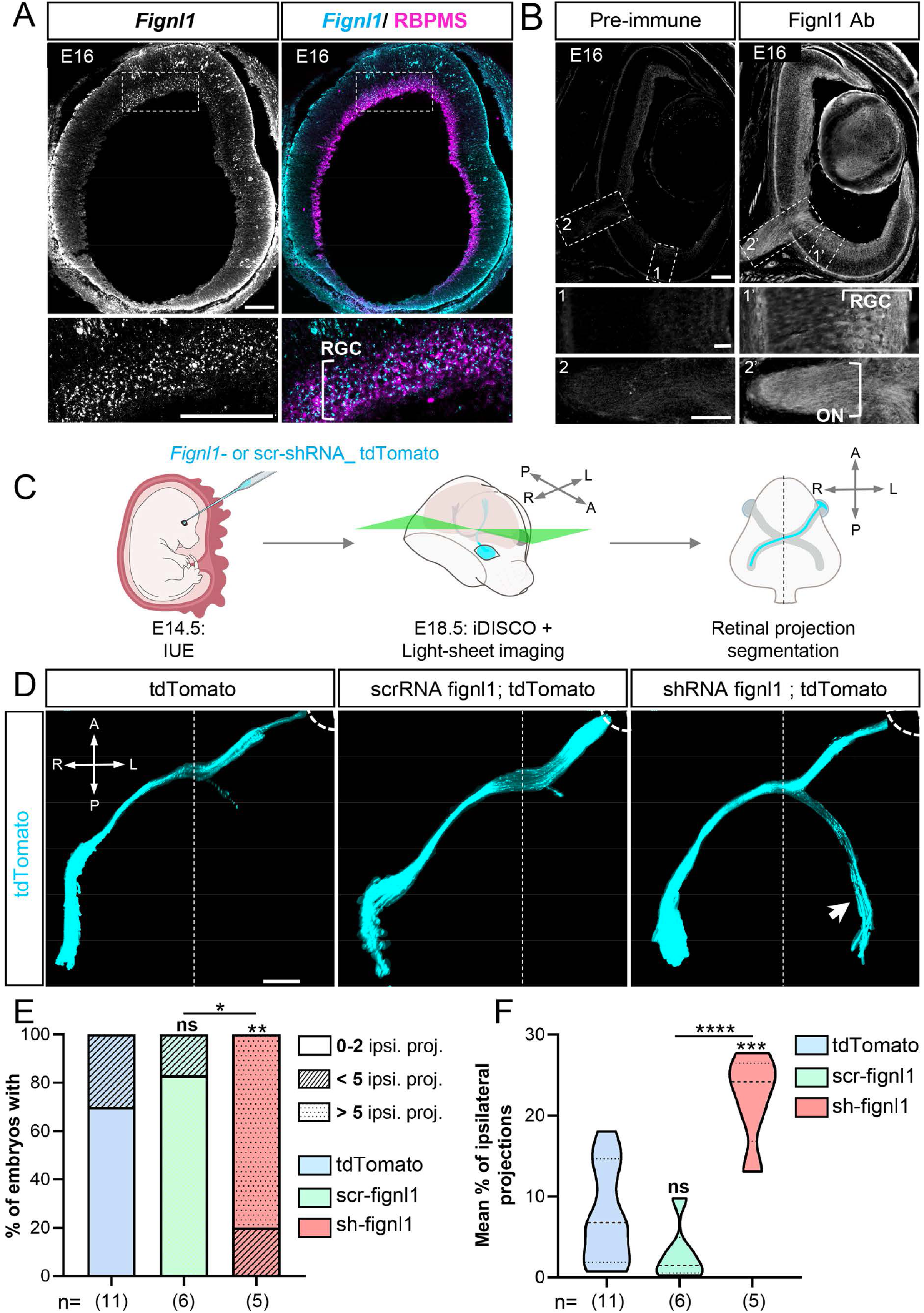
Retina-targeted fignl1 depletion leads to aberrant ipsilateral projections. (A) SABER-FISH using *Fignl1* probes (in cyan) on E16 retinal cryosection post-stained with anti-RBPMS antibody to label RGCs (in magenta). Bottom panels show higher magnifications of the boxed regions in the upper panels. Scale bar (zoom): 100 μm. (B) E16 retinal cryosection immunolabeled with either pre-immune serum or anti-fignl1 antibody. Bottom panels show higher magnifications of the boxed regions 1 (RGC layer) and 2 (optic nerve) in the upper panels. Scale bar: 100 μm. (A-B): RGC: Retinal Ganglion Cell; ON: Optic Nerve. (C) Schematic of the experimental procedure used to investigate the impact of fignl1 knock down on contralateral RGC axon guidance at the midline. IUE: *in utero* electroporation. (D) Z-projection of 3D light-sheet images from E18.5 cleared embryo heads electroporated in a single retina with *Fignl1* or scramble (scr) sh-RNA and tdTomato constructs and immunolabelled with anti-DsRed antibody. Arrow points at aberrant ipsilateral projections. Dotted lines indicate eye position. Scale bar: 500 μm. (E) Bar graph showing the percentage of electroporated embryos from D exhibiting ipsilateral axons. Chi2 test. (F) Violin plot showing the percentage of ipsilateral projections per embryo. Horizontal solid and dotted bars indicate the median ± the 1st and 3rd quartiles, respectively. One-Way ANOVA with Bonferroni’s post-hoc test. (E-F) *p≤0.05; **p≤0.01, *** p≤0.001; **** p≤0.001. ns: non-significant. The number (*n*) of embryos obtained from more than four independent *in utero* electroporation experiments is indicated under each bar or violin plot.

### Fignl1 regulates F-actin network homeostasis to control contralateral RGC axon outgrowth

To further investigate the role of this microtubule-depolymerizing enzyme in contralateral retinal axon guidance, we examined its subcellular distribution in central retina explants, composed exclusively of contralateral RGCs, using both overexpression and CRISPR/Cas9 Knock-in approaches combined with live TIRF imaging (Figure 2A). Intriguingly, while fignl1 localises to microtubules in zebrafish motor neuron axons,^24^ both endogenous and overexpressed fignl1-GFP co-localise with F-actin in the growth cone peripheral domain (P-domain) of mouse RGC axons (Figures 2B and 2C). Specifically, fignl1 is enriched along F-actin filopodia (arrows, Figure 2Cand Video S2), where it moves with the actin retrograde flow (Figures 2D-2E and Video S2), suggesting a physical link between this microtubule-depolymerizing enzyme and the actin cytoskeleton. This interaction was then further dissected in *in vitro* reconstituted systems using a recombinant fignl1ΔN variant lacking amino-acids 1 to 125, a deletion that markedly improves its solubility without impacting its microtubule-depolymerizing activity (Figure S1E-G). F-actin co-pelleting assays showed that fignl1ΔN directly binds F-actin with an apparent stoichiometry of 1μM fignl1ΔN per 1.5μM of actin (ratio = 0.67 ± 0.20; pellet fraction; Figure 2F). TIRF imaging of F-actin polymerisation assays performed in the presence of recombinant wild-type or catalytically dead – defective for ATP hydrolysis (fignl1ΔN_E510A) – fignl1ΔN further demonstrated that fignl1ΔN inhibits both F-actin nucleation and polymerisation in a dose- and ATPase-dependent manner (Figures 2G-2J). These results identify fignl1 as a novel inhibitor of F-actin dynamics. To investigate the mechanisms by which this dual regulator of microtubule and actin dynamics controls axon midline crossing, we generated a retina-specific *Fignl1* conditional mouse mutant. *Fignl1^Fl/Fl^*conditional mutant mice^21^ were crossed with *Dkk3*:Cre transgenic mice, which express the Cre recombinase in retinal progenitor cells^49^. Fignl1 expression was selectively depleted in the retina of *Dkk3*:Cre; *Fignl1^Fl/Fl^* mouse embryos (hereafter referred to as Cre+^Ret^; *Fignl1^Fl/Fl^*; Figures S2A-S2C), with no detectable fignl1 immunoreactivity in primary cultures of retinal explants from Cre+^Ret^; *Fignl1^Fl/Fl^* embryos (Figures S2D and S2E).

**Figure 2:**
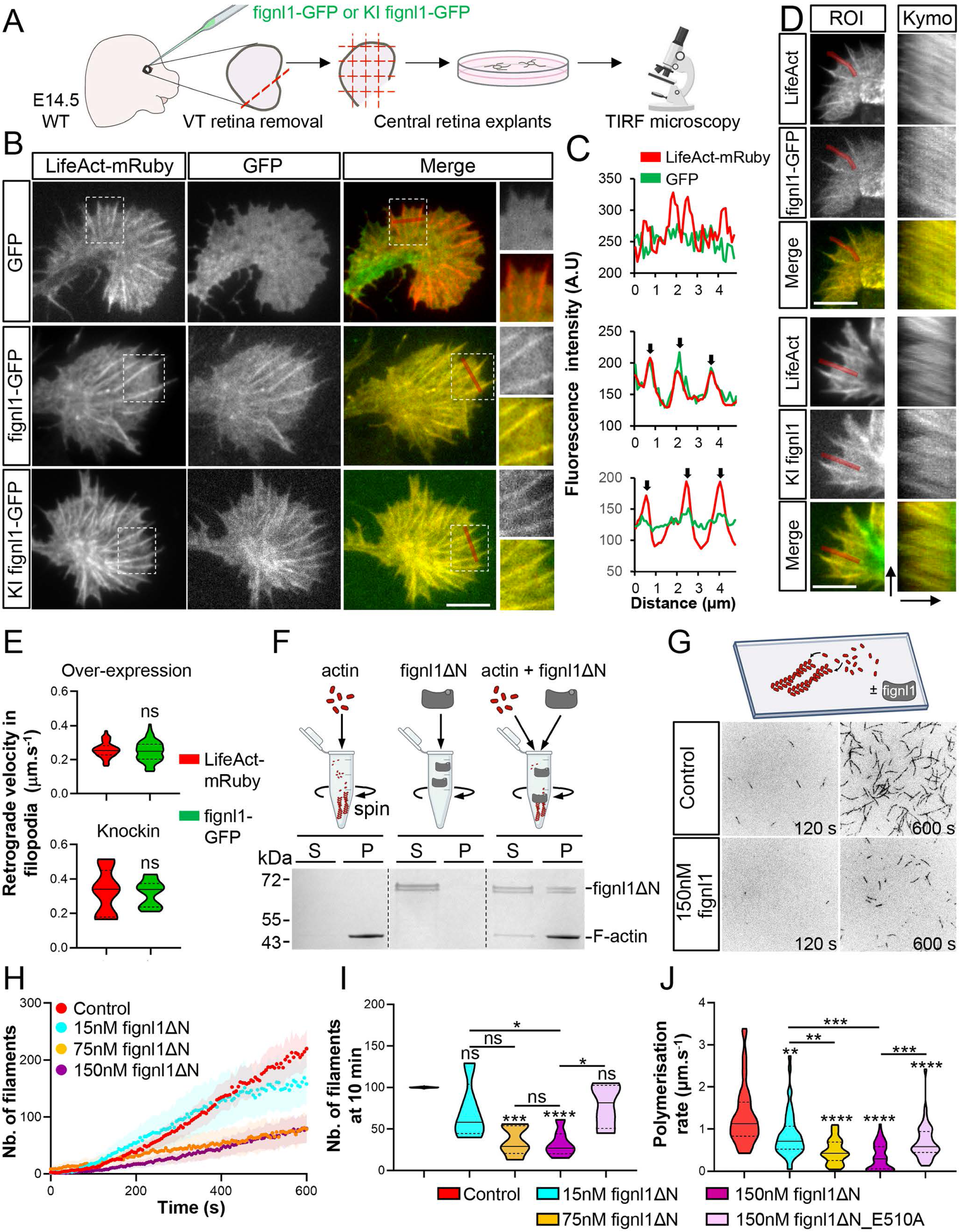
Fignl1 is an actin nucleation inhibitor enriched in RGC growth cone filopodia. (A) Schematic of the protocol used for live TIRF imaging of fignl1-GFP and LifeAct-mRuby in cultured contralateral RGC explants. VT: ventro-temporal; WT: wild-type. (B) Representative still TIRF images of RGC growth cones co-electroporated with fignl1-GFP, fignl1-GFP knockin (KI fignl1-GFP), or GFP control constructs together with a LifeAct-mRuby plasmid. Right panels show higher magnifications of the boxed regions. Red lines indicate ROIs used to generate the fluorescence intensity profiles shown in panel C. Scale bar: 5 μm. (C) Fluorescence intensity profiles for GFP and LifeAct-mRuby across the P-domain of RGC growth cones under each condition. A.U., arbitrary units. Arrows point at fignl1 and F-actin co-localisation in filipodia. (D) Left: Still TIRF images of RGC growth cones co-electroporated with fignl1-GFP or fignl1-GFP knockin (KI fignl1-GFP) constructs together with a LifeAct-mRuby plasmid. Red lines indicate ROIs used for kymograph analysis shown in right panels. Right: Corresponding kymographs showing F-actin and fignl1-GFP or knockin fignl1-GFP dynamics along filopodia. Image scale bar: 5 μm; Kymograph horizontal and vertical scale bars: 3,3 μm and 52 seconds, respectively. (E) Violin Plots showing the retrograde velocity of F-actin (LifeAct-mRuby) and overexpressed or endogenous (KI) fignl1-GFP measured along growth cone filopodia. ns: non-significant. The number of filopodia (*n*) analysed is indicated under each violin plot. (F) Top: Schematic of the F-actin co-sedimentation assay. Bottom: Representative Coomassie-stained SDS-PAGE image showing recombinant fignl1ΔN sedimented in the absence or presence of filamentous actin. S: supernatant; P: pellet. (G) Top: Schematic of the *in vitro* reconstructed F-actin polymerisation assay. Bottom: Representative TIRF time-lapse images of Alexa-Fluor-568-actin polymerization in the absence or presence of fignl1ΔN at the indicated time points. S: second. (H) Dose-response curves showing the effect of increasing concentrations of fignl1ΔN on the number of actin filaments nucleated over time (n = 5 experiments for each condition). (I) Violin plot showing the mean number of actin filaments nucleated, normalized to control, in the absence (n= 7 experiments) or presence of 15 nM (n=5), 75 nM (n= 5) or 150 nM (n= 7) fignl1ΔN, or 150 nM ATPase-dead fignl1ΔN (ΔN_E510A; n=4), measured 10 min after the start of the assay. (J) Violin plot showing the mean actin filament polymerization rate over a 10 min period in the absence (n=70 filaments) or presence of 15 nM (n=50), 75 nM (n=50), or 150 nM (n=70) fignl1ΔN, or 150 nM ATPase-dead fignl1ΔN (ΔN_E510A; n=40), corresponding to the conditions in I. (I-J): *p≤0.05; **p≤0.01, *** p≤0.001; **** p≤0.001. ns: non-significant. One-Way ANOVA with Bonferroni’s post-hoc test (I) and Kruskal–Wallis ANOVA with Dunn’s post hoc test (J). Horizontal solid and dotted bars denote the median ± the 1st and 3rd quartiles, respectively.

The impact of fignl1 depletion on contralateral RGC axon outgrowth was first examined under basal conditions – in the absence of guidance cues –, using primary cultures of central retina explants (Figure 3A). Twelve hours after plating, fignl1-deficient contralateral RGCs display a moderate increase in axon length (Figures 3B and 3C), accompanied by significant changes in growth cone morphology and cytoskeletal organisation (Figure 3D and 3E). STED microscopy analysis revealed that Fignl1 depletion significantly reduces growth cone size by decreasing the P-domain area, which is primarily composed of a branched F-actin network forming lamellipodia (Figures 3D and 3F). Conversely, it increases the number of linear F-actin-based filopodia, suggesting an unexpected role for fignl1 in controlling F-actin network homeostasis (Figures 3D and 3G). These structural changes of the actin cytoskeleton were associated with enhanced microtubule invasion into the P-domain (*i.e.* increased number and penetration distance; Figures 3H and 3I), including filopodial invasion (Figures 3J), a phenotype consistent with the axonal growth-promoting effect of fignl1 depletion.

**Figure 3:**
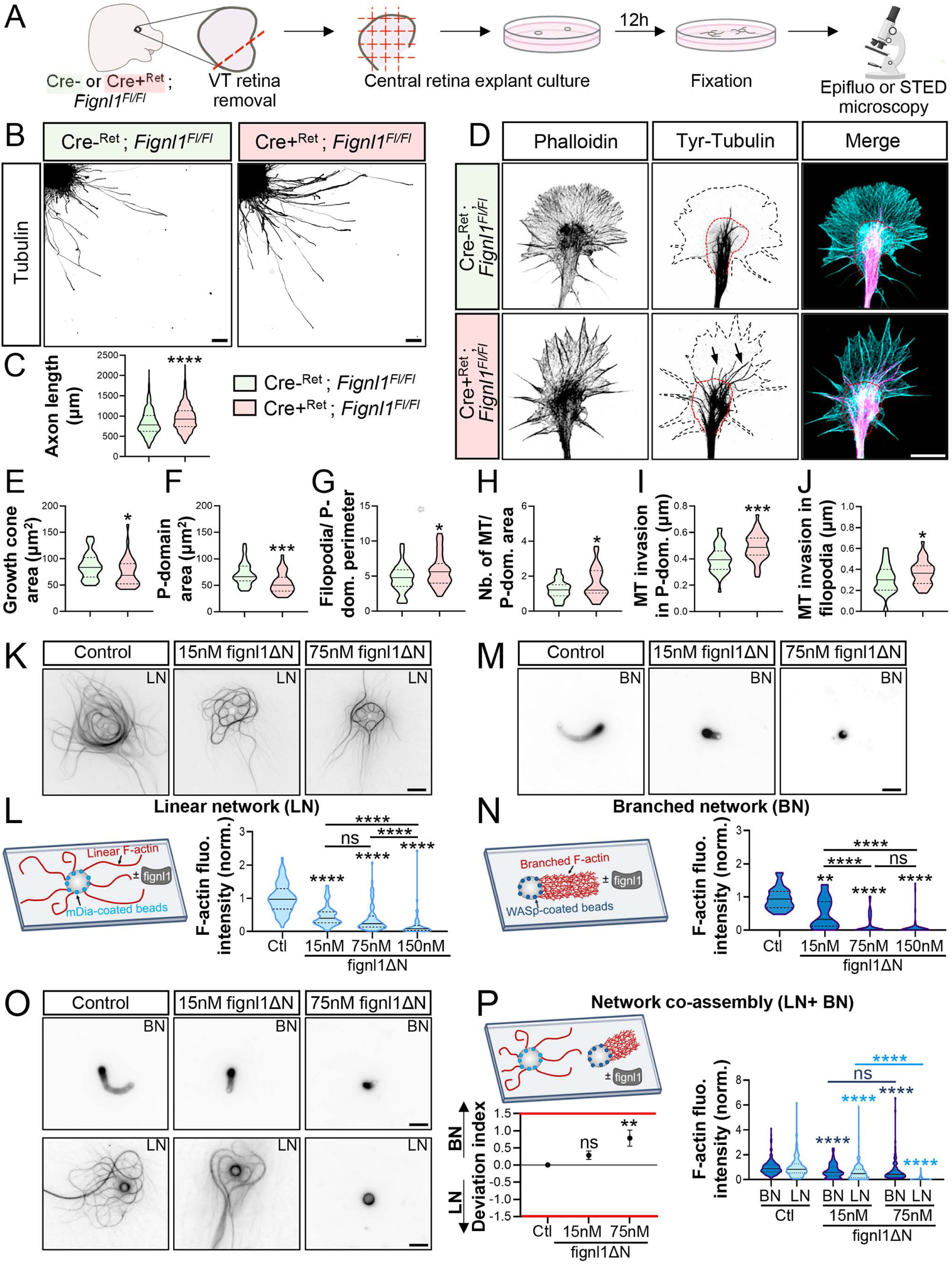
Balanced control of F-actin network dynamics by fignl1 controls contralateral RGC axon outgrowth. (A) Schematic of the retinal explant culture protocol used to assess the impact of fignl1 loss of function on contralateral RGC (cRGC) axon outgrowth and growth cone morphology. (B) Representative epifluorescence images of central retina explants from Cre-^Ret^; *Fignl1^Fl/Fl^* RGC and Cre+^Ret^; *Fignl1^Fl/Fl^* embryos immunolabelled with tubulin antibody. Scale bar: 100 μm. (C) Violin plot showing the mean axon length of control (n=738) and fignl1-deficient (n=620) contralateral RGC axons, measured on at least 31 explants pooled from three independent experiments. (D) Representative z-projections of deconvoluted 3D-STED images of growth cones from Cre-^Ret^; *Fignl1^Fl/Fl^* and Cre+^Ret^; *Fignl1^Fl/Fl^* RGCs stained for tyrosinated tubulin (magenta) and F-actin (phalloidin, cyan). Red and black dotted lines indicate the boundary between the central domain and the F-actin-rich peripheral domain (P-domain), and the P-domain perimeter, respectively. Arrows indicate the increased microtubule invasion of in the P-domain. Scale bar: 5 μm. Tyr-tubulin: tyrosinated tubulin. (E-J) Violin plots showing the mean growth cone area (E), P-domain area (F), number of filopodia per P-domain perimeter (G), number of MT per P domain area (H) and MT invasion distance in the P-domain (I) or filopodia (J), quantified from STED-imaged growth cones of Cre-^Ret^; *Fignl1^Fl/Fl^* (n= 38) and Cre+^Ret^; *Fignl1^Fl/Fl^* (n=39) RGCs, pooled from three independent experiments. MT: microtubules. (K, M, O) Epifluorescence images of linear (LN; filament-like) or branched (BN; comet-like) F-actin networks nucleated separately (K, M) or co-assembled in the same chamber (O) in the presence or absence of 15nM or 75nM fignl1ΔN. Images were acquired 30 min after nucleation initiation. Scale bar: 5 μm. (L, N, P) Left: Schematics of the *in vitro* F-actin nucleation assays used to assess the effects of fignl1ΔN on linear (L) or branched (N) F-actin network assembled around mDia- or WASp-coated microbeads, respectively, or on their co-assembly (P). Right (L, N): Violin plots showing the mean fluorescence intensity of linear (LN) or branched (BN) F-actin network assembled separately in the presence of 15nM (LN: n=74 beads; BN: n=46), 75 nM (LN: n=74; BN: n=53) or 150 mM fignl1ΔN (LN: n=70; BN: n=57), normalized to values obtained in the absence of fignl1ΔN (LN: n=72; BN: n=47), set to 1. Right (P): Violin plots showing the mean fluorescence intensity of linear or branched F-actin co-assembled in the presence of 15 nM (LN: n=142 beads; BN: n=177) or 75 nM fignl1ΔN (LN: n=82; BN: n=90), normalized to values obtained in the absence of fignl1ΔN (LN: n=150; BN: n=167), set to 1. (L, N, P) Data are pooled from 4 independent experiments. Bottom left (P): Deviation index quantifying shifts in actin assembly rates around WASp versus mDia-coated beads relative to a balanced state (in which both networks assemble optimally, Deviation Index = 0) in the presence or absence of 15 or 75 nM fignl1ΔN. Ctl: Control condition. (C, E-J, L, N, P): *p≤0.05; **p≤0.01, ***p≤0.001; ****p≤0.001. ns: non-significant. Mann Whitney test (C, E), unpaired t-test (F, G, H, I, J) and Kruskal–Wallis ANOVA with Dunn’s post hoc test (L, N, P). Horizontal solid and dotted bars in violin plots indicate the median ± the 1st and 3rd quartiles, respectively.

Given the impact of fignl1 on branched and linear F-actin network homeostasis in RGC growth cones, its specific and direct effects on each network were examined in cell-free reconstituted systems. Defined nucleation assays were used to selectively trigger linear or branched F-actin network formation around mDia- or WASp-coated beads, respectively, as previously described^50^ (Figure 3K-3N). When the two networks are nucleated in separate glass chambers, fignl1ΔN exhibits similar dose-dependent inhibitory effects on both linear and branched F-actin network assembly (Figures 3K-3N). However, when both networks co-polymerize in the same chamber, fignl1ΔN preferentially inhibits linear network assembly at 75nM (Figures 3O and 3P), thereby attenuating its effect on the branched network at this concentration. Together with STED analysis of RGC growth cones, these results indicate that fignl1 regulates F-actin network homeostasis during RGC axon outgrowth by biasing nucleation toward branched over linear networks (Figure 3P).

### Fignl1 mediates Wnt5a-induced contralateral RGC axon repulsion via F-actin nucleation inhibition

The identification of fignl1 as a novel F-actin inhibitor raised the possibility that it links actin regulation to axon guidance pathways. Consistent with this idea, loss of fignl1 function led to an excess of ipsilateral projections (Figures 1D-F), a phenotype reminiscent of that observed in embryos with defective non-canonical Wnt5a/β-catenin signalling,^32^ suggesting that fignl1 may act downstream of this guidance pathway. To test this hypothesis, we performed *in vitro* guidance assays combined with live differential interference contrast (DIC) imaging to assess how wild-type and fignl1-deficient contralateral RGC axons respond to the Wnt5a cue (Figure 4A and Video S3). Central retina explants were treated 24 h post-plating with recombinant Wnt5a (300 ng.mL^−1^; bath application) or PBS as vehicle control. Wnt5a acts as a potent repulsive cue, rapidly inducing growth cone collapse – characterized by disassembly of F-actin structures in the P-domain – followed by axon retraction in both control and fignl1-deficient contralateral RGCs, in contrast to PBS treatment (Figures 4B-4C and Video S3). However, loss of fignl1 function impairs the responsiveness of contralateral RGC axons to Wnt5a, leading to their premature desensitization to this repellent cue (arrows, Figures 4C-4M and Video S3). This is reflected by a higher proportion of fignl1-deficient axons resuming extension before the end of the 1-hour imaging period compared to controls (Figures 4E and 4F). Although control and fignl1-deficient axons exhibits comparable growth dynamics before Wnt5a exposure (Figures 4G and 4H), and similar growth cone collapse and axon retraction kinetics upon Wnt5a treatment (Figures 4I and 4K), the overall retraction distance of fignl1-deficient contralateral axons is significantly reduced relative to controls (Figure 4J). This reduction reflects the inability of fignl1-deficient growth cones to sustain Wnt5a-induced inhibition of their protrusive activity – driven by F-actin nucleation and polymerization –, resulting in the premature rebuilding of growth-competent growth cones (Figure 4C and 4L) and earlier axon regrowth (Figure 4C and 4M). Consistent with this, TIRF imaging of fixed central retina explants, reveals that 10 min after Wnt5a exposure, approximately 40% of control growth cones are collapsed, displaying a pronounced loss of F-actin structures within the P-domain (Figures 4N-4P). In contrast, the vast majority of fignl1-deficient contralateral growth cones (≈80%) are not collapsed and exhibit an F-actin-rich P-domain, with an area similar to that of PBS-treated growth cones (Figures 4N-4P). These findings align with the inhibitory activity of fignl1 on F-actin nucleation and polymerization observed in *in vitro* reconstitution systems (Figure 2G-J). Together, these data demonstrate that fignl1 drives Wnt5a-evoked repulsion of contralateral RGC axons by inhibiting F-actin nucleation.

**Figure 4:**
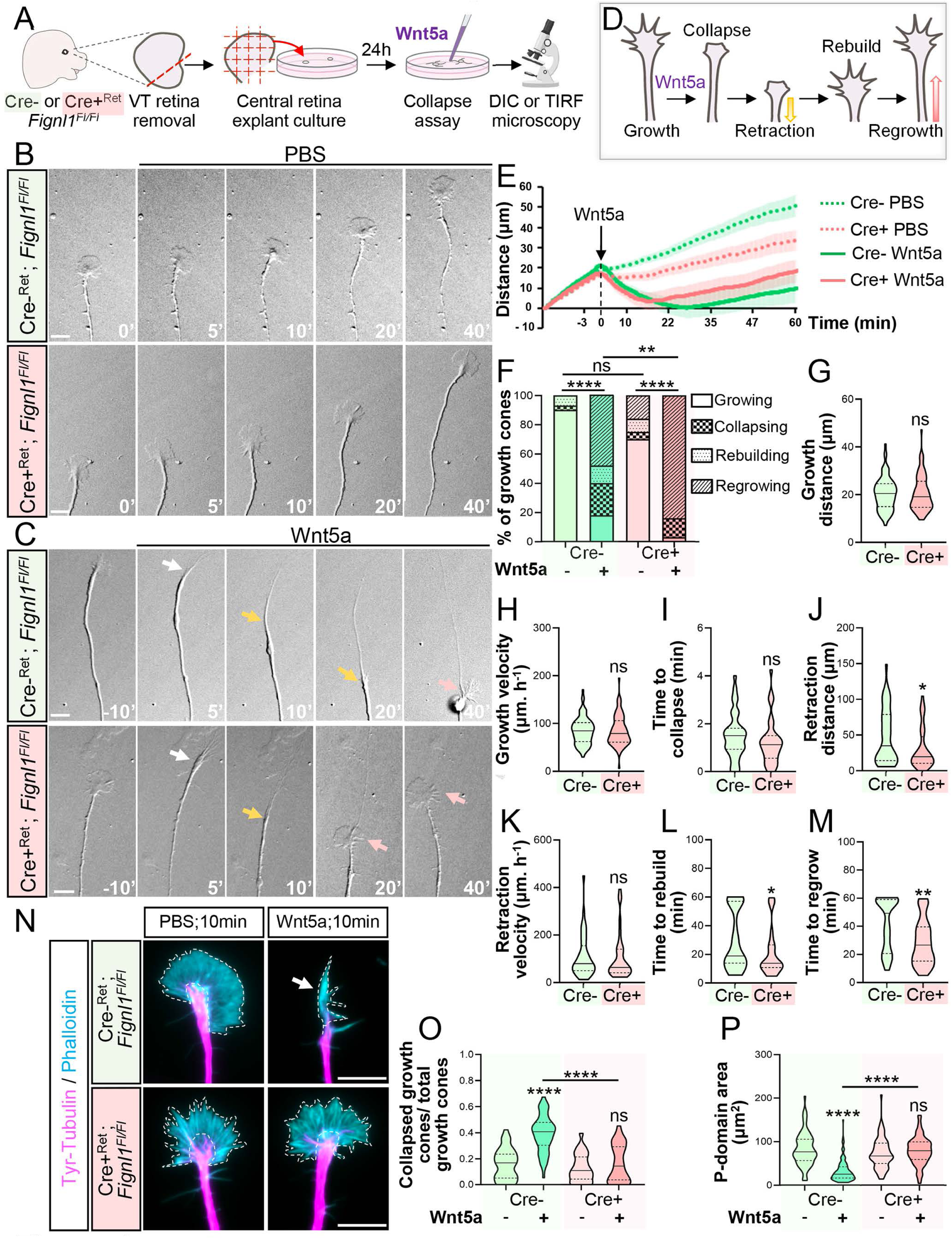
Fignl1 maintains Wnt5a-induced inhibition of actin nucleation to sustain contralateral axon repulsion. (A) Schematics of the culture and imaging protocol used for Wnt5a collapse assays. Axons of control and fignl1-depleted contralateral RGCs (cRGC) were imaged for 75 minutes (1 frame/15 s). PBS or Wnt5a (300 ng/mL) was added 15 minutes after the start of imaging. (B-C) Representative still DIC images of control (Cre-^Ret^; *Fignl1^Fl/Fl^*) and fignl1-depleted (Cre+^Ret^; *Fignl1^Fl/Fl^*) cRGC axons acquired before (0 min) and 5, 10, 20 and 40 minutes after exposure to PBS (Sham; B) or Wnt5a (300ng.mL^−1^; C). White arrows indicate collapsed growth cones. Orange and Pink arrows point at retracting and prematurely regrowing axons, respectively. Scale bar: 10 μm. (D) Schematic illustrating the sequence of RGC growth cone behaviours triggered by Wnt5a treatment. (E) Traces showing the average directed growth length of axons analysed in live DIC imaging: control Cre-PBS (n= 29), Cre+ PBS (n= 44), Cre-Wnt5a (n= 51) and Cre+ Wnt5a (n= 38). (F) Bar graph showing the distribution of axon behaviours over the 75-min imaging period from panel (E). Growth cones were classified as Growing (no collapse), Collapsing (collapse only), Rebuilding (collapse followed by growth cone rebuilding), or Regrowing (collapse, growth cone reformation, and resumption of axon extension).**p≤0.01, **** p≤0.001, ns: non-significant. Chi2 test. (G-M) Violin plots showing the axon growing distance (G) and axon growth velocity (H) of cRGC axons before Wnt5a stimulation, the elapsed time between Wnt5a addition and growth cone collapse (I), the axon retraction distance (J) and velocity (K), and the elapsed time between growth cone collapse and growth cone rebuilding (L) or axon regrowth (M). Only axons collapsing in response to Wnt5a are included in (G-M) quantifications: Cre-Wnt5a (n=42) and Cre+ Wnt5a (n=36) axons pooled from at least 3 independent experiments. (N) Representative TIRF images of control (Cre-^Ret^; *Fignl1^Fl/Fl^*) and fignl1-deficient (Cre+^Ret^; *Fignl1^Fl/Fl^*) cRGC growth cones treated for 10 min with PBS or Wnt5a and labelled for tyrosinated tubulin (magenta) and F-actin (phalloidin, cyan). Dotted lines delineate the P-domain perimeter. Arrow points at a collapsed growth cone. Scale bar: 10μm. (O) Violin plot showing the proportion of collapsed growth cones in control and fignl1-deficient central retina explants treated with Wnt5a (Cre-Wnt5a: n=58; Cre+ Wnt5a: n=49) or PBS (Cre-PBS: n=35; Cre+ PBS: n=58) obtained from three independent experiments. (P) Violin plot showing the P-domain area of control and fignl1-deficient cRGC growth cones treated with Wnt5a (Cre-Wnt5a: n= 100; Cre+ Wnt5a: n= 70) or PBS (Cre-PBS: n=82; Cre+ PBS: n=75) obtained from 3 independent experiments. (G-M, O, P) *p≤0.05; **p≤0.01, **** p≤0.001. ns: non-significant. Mann Whitney test (G-M) and Kruskal–Wallis ANOVA with Dunn’s post-hoc test (O, P). Horizontal solid and dotted lines in violin plots indicate the median ± the 1st and 3rd quartiles, respectively.

### PKC-mediated phosphorylation reprograms fignl1 cytoskeletal activity to sustain Wnt5a-induced axon repulsion

We next investigated how Wnt5a controls the cytoskeleton-related activities of fignl1 to promote midline crossing of contralateral retinal axons. Using i*n silico* analysis, we identified seven putative phosphorylation sites for the non-canonical Wnt5a signalling downstream protein kinase, PKC, five within the RAD51-binding domain and two within the microtubule-binding domain of fignl1. PKC phospho-resistant (fignl1-PKC-7A) and phospho-mimetic (fignl1-PKC-7D) variants targeting these residues were generated (Figure 5A). Using both *in cellulo* and *in vitro* PKCα kinase assays, we demonstrated that fignl1 is indeed phosphorylated by PKCα on a subset of these residues (Figures 5B and 5C). Although PKCα has been implicated in RGC axon routing at the optic chiasm,^51^ its function downstream of Wnt5a in midline crossing of RGC axons has never been examined. We thus performed *in vitro* growth cone collapse assays on central retina explants expressing a dominant-negative PKC variant fused to GFP (PKC-DN) or GFP alone. Genetic inhibition of PKCα activity impairs contralateral RGC axon responsiveness to Wnt5a-induced repulsion (reflected by reduced percentage of collapsed growth cones, Figures 5D and 5E) similar to fignl1 loss of function (Figure 4P). To determine how PKC-mediated phosphorylation of fignl1 modulates its cytoskeletal activities, we employed both genetic (overexpression of phospho-resistant and phospho-mimetic fignl1ΔN variants; Figures 5F-H) and pharmacological (PKC inhibition with Gö6983 or activation with PMA; Figures S3A-C) approaches in COS-7 cells. PKC-mediated phosphorylation of fignl1 inhibits its microtubule-depolymerizing activity (Figures 5F and 5G, Figures S3A and S3B) while significantly enhancing its co-localization with F-actin at the cortex (arrows, Figures 5F and 5H, Figures S3A and S3C). To test the functional significance of these effects for Wnt5a-evoked contralateral RGC axon repulsion, we next conducted rescue experiments in primary cultures of central retina explants from retina-specific fignl1conditional mutant embryos (Cre+^Ret^; *Fignl1^Fl/Fl^*mice embryos) using wild-type and PKC phospho-resistant (fignl1ΔN-PKC-7A) variants of fignl1ΔN (Figure 5I). Expression of wild-type fignl1ΔN fully rescues Wnt5a-induced responses in fignl1-deficient contralateral RGC axons, restoring control levels of growth cone collapse (+/- 50% of collapsed growth cones; Figure 5J and 5K versus Figure 4N and 4O). In contrast, the phospho-resistant fignl1ΔN-PKC-7A variant fails to restore Wnt5a responsiveness (Figures 5J and 5K). Together, these results demonstrate that PKC-mediated phosphorylation of fignl1 is required for cytoskeletal remodelling driving Wnt5a-evoked contralateral axon repulsion *in vitro*.

**Figure 5:**
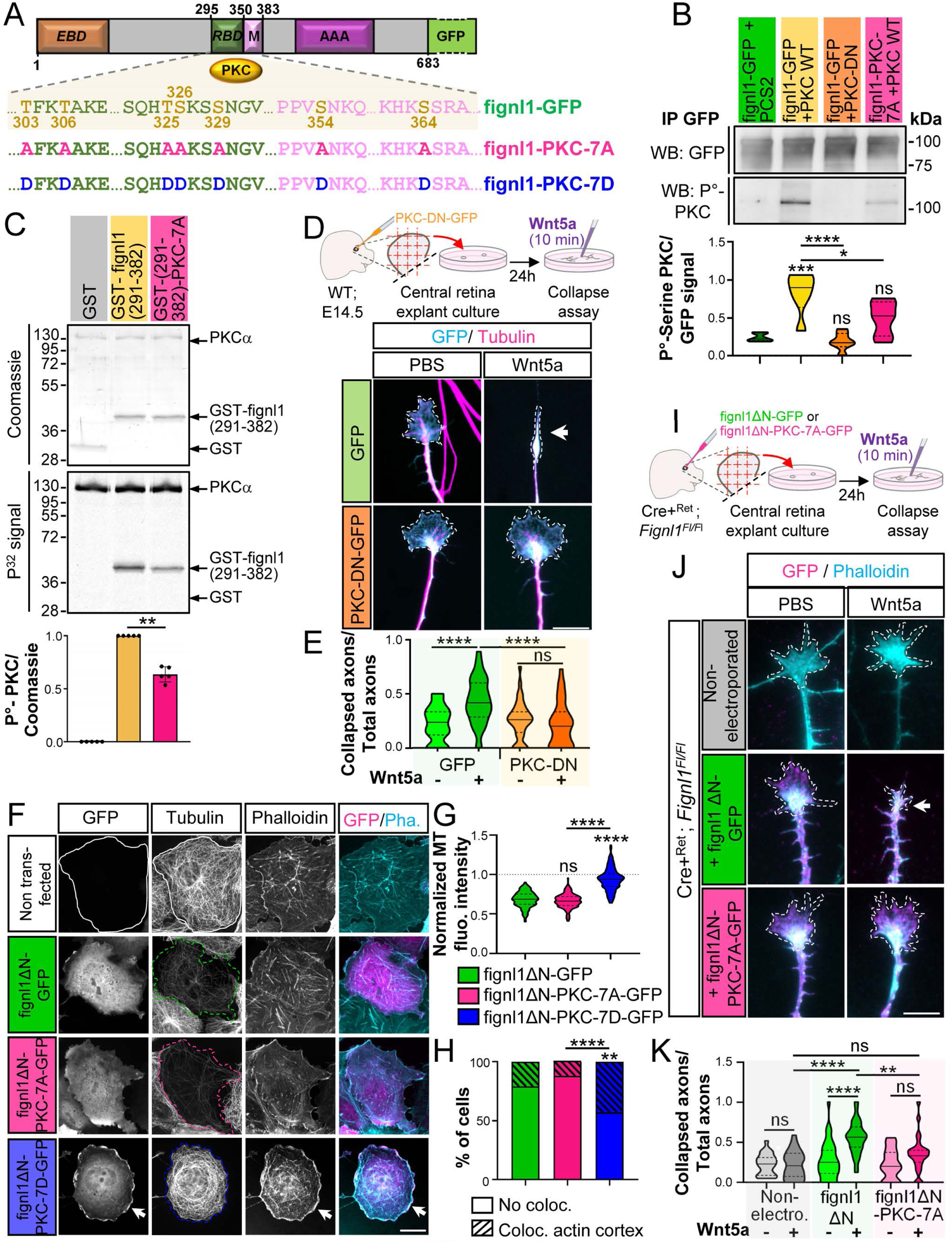
PKC phosphorylation of fignl1 maintains contralateral axon sensitivity to Wnt5a by inhibiting F-actin nucleation. (A) Schematic representation of the putative PKC phosphorylation sites identified in the fignl1 RAD51-binding (RBD) and microtubule-binding (M) domains, and of the phospho-resistant (fignl1-PKC-7A) and -mimetic (fignl1-PKC-7D) variant used. (B) *In cellulo* PKC phosphorylation assays. top: Representative Western blot analysis of immunoprecipitation assays performed with a GFP antibody on protein extracts from COS-7 cells co-transfected with fignl1-GFP or fignl1-PKC-7A-GFP and either wild-type (PKC-WT) or catalytically dead (PKC-DN) PKC variants or an empty pCS2 control plasmid. Blots were probed with Phospho-(Ser) PKC substrate and GFP antibodies. Lower panel: Violin plots showing the Phospho-(Ser) PKC signal normalized to GFP signal obtained from n=6 experiments. (C) Top: *In vitro* PKCα kinase assays using 0.2 μM GST-fignl1(291-382), GST-fignl1(291-382)-PKC-7A, or GST as substrates. Bottom: Quantification of phospho-fignl1 signal normalized to Coomassie staining and expressed relative to wild-type fignl1(291-782) (set to 1), from n=5 independent experiments. (D) Top: Schematic of *in vitro* Wnt5a growth cone collapse assays conducted with cultured contralateral retinal explants electroporated with GFP or PKC-DN-GFP. Bottom: Representative epifluorescence images of cRGC growth cones electroporated with GFP or PKC-DN-GFP and immunostained with GFP and tyrosinated tubulin antibodies 10 min after PBS or Wnt5a treatment. Dotted lines outline the P-domain perimeter. Arrow points at a collapsed growth cone. Scale bar: 10μm. (E) Violin plot showing the proportion of collapsed growth cones in central retina explants overexpressing GFP or PKC-DN-GFP exposed to PBS (GFP: n=35; PKC-DN: n=39) or Wnt5a (GFP: n=34; PKC-DN: n=34) from D, and pooled from three independent experiments. (F) Representative epifluorescence images of COS-7 cells overexpressing GFP-tagged fignl1ΔN, fignl1ΔN-PKC-7A, or fignl1ΔN-PKC-7D, and stained with anti-tyrosinated tubulin antibody and phalloidin. Solid and dotted lines indicate non-transfected and transfected cells, respectively. Arrows indicate fignl1ΔN-PKC-7D colocalization with the cortical actin network. Scale bar: 20μm. (G) Violin plot showing microtubule fluorescence intensity in COS-7 cells overexpressing fignl1ΔN (n=186), fignl1ΔN -PKC-7A (n=219), or fignl1ΔN-PKC-7D (n=221) from F, normalized to non-transfected cells. (H) Bar graph showing the percentage of cells overexpressing fignl1ΔN (n=186), fignl1ΔN-PKC-7A (n=219), or fignl1ΔN-PKC-7D (n=221) from F that exhibit significant colocalization of fignl1 with cortical F-actin. (I) Schematic of rescue experiments performed in fignl1-deficient (Cre+^Ret^; *Fignl1^Fl/Fl^*) central retina explants electroporated with fignl1ΔN-GFP or fignl1ΔN-PKC-7A-GFP constructs. (J) Representative epifluorescence images of growth cones of fignl1-deficient cRGCs electroporated or not with fignl1ΔN-GFP or fignl1ΔN-KC-7A-GFP, treated for 10 min with PBS or Wnt5a, and stained with anti-GFP antibody and phalloidin. Dotted lines outline the P-domain. The arrow points at a collapsed growth cone. Scale bar: 10μm. (K) Violin plots showing the proportion of collapsed axons in fignl1-deficient retinal explants electroporated or not with fignl1ΔN-GFP or fignl1ΔN-PKC-7A-GFP and treated with PBS (non-electroporated: n=24, fignl1ΔN: n=39, fignl1ΔN-PKC-7A: n=27) or Wnt5a (non-electroporated: n=26, fignl1ΔN: n=42, fignl1ΔN-PKC-7A: n=27), pooled from three independent experiments. (B, E, G-H, K) **p≤0.01, *** p≤0.001 **** p≤0.0001. ns: non-significant. One-Way ANOVA with Bonferroni’s post-hoc test (B, E, G), Kruskal–Wallis with Dunn’s post-hoc test (K), and Chi2 test (H). Horizontal solid and dotted lines in violin plots indicate the median ± the 1st and 3rd quartiles, respectively.

### The RAD51/fignl1 complex functions as a key Wnt5a/PKC signalling integrator required for retinal axon repulsion

I*n silico* analyses were used to assess the evolutionary conservation of PKC phospho-sites previously identified in the fignl1 sequence (Figure 5A). Residues within the fignl1 RBD display low to high conservation across binocular species – (T325< T303< T326< T306< S329; Figure S4A), whereas MTBD residues show little conservation. In non-binocular species only the T306 residue within the RBD remains conserved (Figure S4B). While the RAD51/fignl1 complex is known for its key nuclear role in maintaining genome integrity,^17, 20–23^ RAD51 mutations also cause Congenital Mirror Movements, a neurodevelopmental disorder associated with excessive ipsilateral corticospinal projections.^8,12^ These observations prompted us to investigate potential non-canonical roles for RAD51 in Fignl1-mediated axon midline crossing. SABER-FISH analysis confirms that *Rad51*, like *Fignl1*, is expressed in RGCs at E16, a critical developmental stage for RGC axon navigation at the chiasm (Figure S5A). Using CRISPR/Cas9-mediated GFP-tagging of RAD51 in central retina explants combined with TIRF imaging (Figure 6A), we observed robust localization of RAD51 along F-actin filopodia of RGC growth cones (Figure 6B), mirroring fignl1 distribution (Figures 2B and 2C). To probe fignl1/RAD51 interactions in growth cones of living contralateral RGC axons, we employed the splitFAST fluorescence complementation system^52^ and TIRF imaging (Figures 6C-D). Consistent with their respective subcellular distributions, a specific splitFAST signal (lime fluorescence) – reporting binding between fignl1-CFAST and RAD51-NFAST – was detected along growth cone filopodia in all growth cones examined. This signal is abolished when using a fignl1 variant deficient for RAD51 binding^17^ (F304E/A307E-CFAST) or a non-interacting NFAST-FRB control protein (0% of growth cone with lime signal for both conditions; Figures 6E). These results demonstrate that fignl1 and RAD51 form a complex along F-actin filopodia in RGC growth cones. To test whether RAD51 locally acts as an anchoring protein for fignl1, we examined the subcellular distribution of two fignl1 variants defective in RAD51 binding (Fignl1-ΔRBD or F304E/A307E)^17^ in RGC growth cones using live TIRF imaging. Both variants show dramatically reduced filopodia targeting compared to wild-type fignl1 (Figures S5B and 5C), demonstrating that RAD51 recruits fignl1 to the actin-rich P-domain of RGC growth cones. Given that PKC phosphorylation sites of fignl1 conserved in binocular species lie within its RBD, we next assessed whether PKC-mediated phosphorylation of fignl1 influences its binding to RAD51 using splitFAST assays in COS-7 cells. PKC-mediated phosphorylation of fignl1 is not required for fignl1/RAD51 complex formation but instead promotes its disassembly (Figures S5D-5F). Conversely, *in vitro* kinase assays revealed that RAD51 enhances PKC-mediated phosphorylation of fignl1, an effect lost with the RBD phospho-resistant variant of fignl1 (Fignl1-PKC-5A; Figure 6F). Structural modelling suggested a mechanistic basis: RAD51 binding to the fignl1 RBD modulates accessibility of this domain to PKC, redirecting PKC docking and phosphorylation site selection. In the absence of RAD51, PKC docks to the fignl1 RBD and targets residues T303/T306 (Figure S5G). Conversely, RAD51 binding to fignl1 repositions PKC onto the fignl1 RBD, which induces a conformational rearrangement that facilitates phosphorylation of the highly conserved S329 residue (Figure S5G). Strengthening this model, mutation of serine 329 to alanine (S329A) abolished RAD51-mediated stimulation of fignl1 phosphorylation by PKC in *in vitro* kinase assays (Figure 6F). Together, these findings suggest that RAD51 acts as a scaffolding protein for fignl1, both recruiting it to filopodia and enabling its phosphorylation at S329 by the Wnt5a downstream kinase PKC. This interaction may represent a critical signalling event driving the actin remodelling required for Wnt5a-evoked midline crossing of retinal axons. These data thus support a role for the RAD51/fignl1 complex as a potential Wnt5a/PKC signalling integrator.

**Figure 6:**
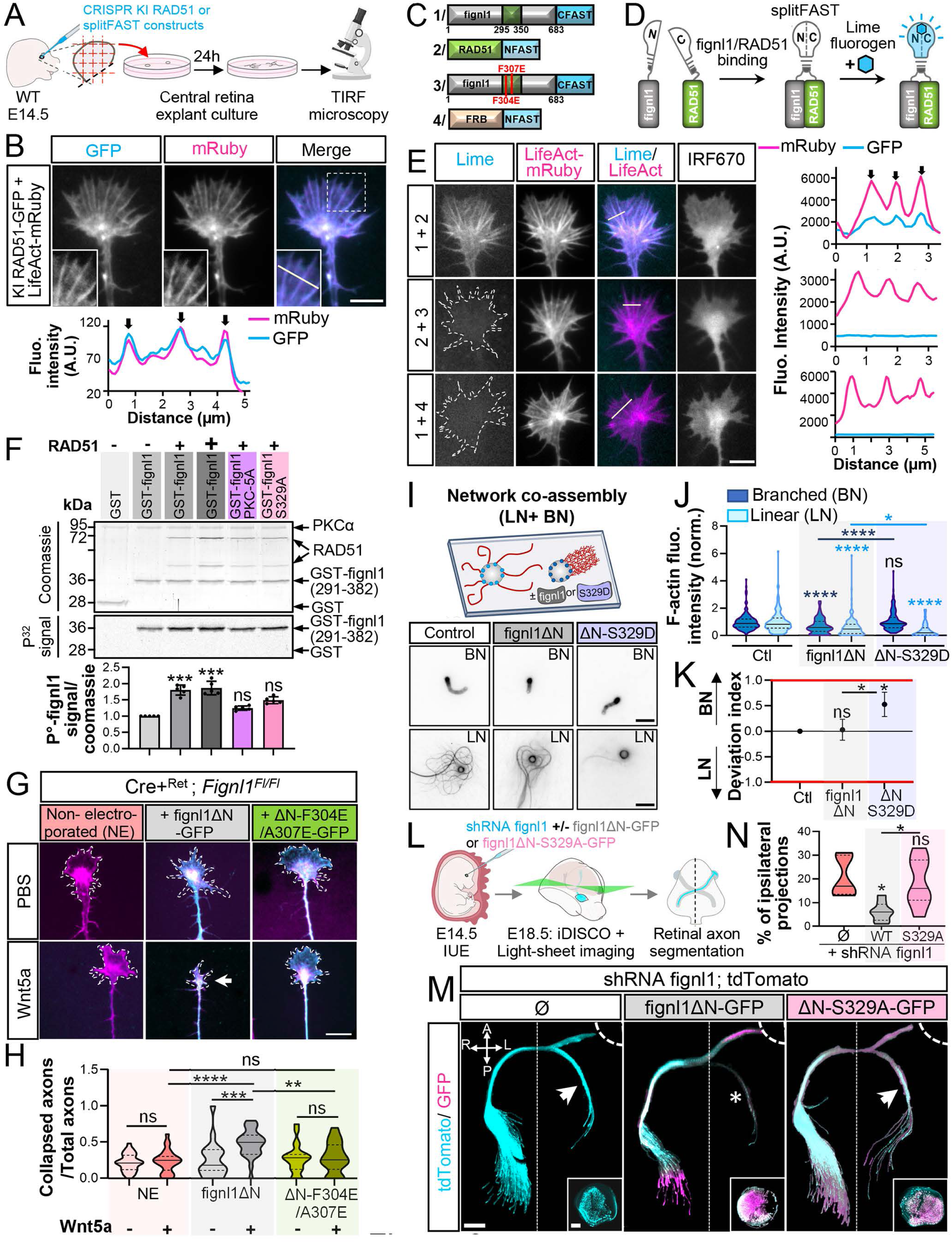
RAD51/fignl1 complex integrates Wnt5a/PKC signalling to mediate retinal axon repulsion. (A) Schematic of the protocol used to monitor RAD51-GFP subcellular distribution (KI RAD51-GFP) and fignl1-RAD51 interaction (splitFAST) in living cRGC growth cones. (B) Top: Representative TIRF images of RGC growth cones co-electroporated with KI RAD51-GFP and LifeAct-mRuby plasmids. Insets show enlarged views of filopodia from the boxed regions. White line indicates the ROI used to analyse the Lime and LifeAct-mRuby fluorescence intensity profiles in the growth cone P-domain (bottom). Arrows indicate Lime signal along filopodia. Scale bar: 5 μm. (C) Schematic of the constructs used in splitFAST assays. FRB: FKBP12-rapamycin binding domain (of mTOR). (D) Schematic of the splitFAST principle and design used to detect fignl1-RAD51 interaction. (E) Left: Representative TIRF images of RGC growth cones co-electroporated with LifeAct-mRuby and the indicated NFAST and CFAST constructs and incubated with 10μM lime fluorogen. Lime fluorescence reports fignl1-RAD51 interaction along filopodia. Dotted lines outline the growth cone perimeter. White lines indicate ROIs used to analyse the Lime and LifeAct-mRuby fluorescence intensity profiles in the growth cone P-domain (right). Arrows indicate filopodia. Plot profile analysis were performed on n= 29, 26 and 9 growth cones for conditions (1+2), (2+3) and (1+4), respectively. Scale bar: 5 μm. (F) Top: *In vitro* PKCα kinase assays using 0.2 μM wild-type, PKC-5A (RBD mutant), or S329A fignl1(291-382) fragments as substrates in the presence or absence of 0.2 or 0.4 μM RAD51. Bottom: Quantification of phospho-fignl1 signal normalized to Coomassie staining and expressed relative to wild-type fignl1(291-382) assayed in the absence of RAD51 (set to 1). Dots represent individual biological replicates (n=5). (G) Representative epifluorescence images of rescue experiments performed in fignl1-deficient central retina explants (Cre+^Ret^; *Fignl1^Fl/Fl^*) electroporated or not (NE) with fignl1ΔN-GFP or fignl1ΔN_F304E/A307E-GFP mutant (RAD51 binding-deficient) constructs, treated for 10 min with PBS or Wnt5a, and stained with anti-GFP antibody (cyan) and phalloidin (magenta). Dotted lines outline the P-domain perimeter. Arrow indicates a collapsed growth cone. Scale bar: 10 μm. (H) Violin plots showing the proportion of collapsed axons in fignl1-deficient retinal explants electroporated or not (NE) with fignl1ΔN-GFP or fignl1ΔN_F304E/A307E-GFP constructs and treated for 10 min with PBS (NE: n=31, fignl1ΔN: n=19, ΔN_F304E/A307E: n=21) or Wnt5a (NE: n=27, fignl1ΔN: n=26, ΔN_F304E/A307E: n=18). Explants were pooled from three independent experiments. (I) Top: Schematic of the *in vitro* reconstitution assays used to test the effect of PKC-mediated phosphorylation of fignl1ΔN on linear and branched F-actin network co-assembly. Bottom: Representative epifluorescence images of linear (LN) and branched (BN) F-actin networks co-assembled in the absence or presence of 15 nM fignl1ΔN or fignl1ΔN-S329D. Images were captured 30 min after initiation of the assay. Scale bar: 5 μm. (J) Violin plot showing mean fluorescence intensity of linear and branched F-actin networks co-assembled around mDia-(LN) or WASp-coated beads (BN), respectively, in the presence of 15nM of fignl1ΔN (LN: n=142 beads; BN: n=177) or fignl1ΔN-S329D (LN: n=82; BN: n=130). Values were normalized to the mean F-actin fluorescence intensity measured in the absence of fignl1ΔN and set to 1 (LN: n=150; BN: n=167). Data were pooled from 5 independent experiments. Control (Ctl) and fignl1ΔN datasets from Figure 2 were used as controls for comparison with fignl1ΔN-S329D, since all experiments were performed in the same batch. Fignl1ΔN-S329D data are shown separately for clarity and to preserve the logical flow of the study. (K) Deviation index showing shifts in actin assembly rates around WASp-versus mDia-coated beads related to a balanced state (Deviation Index = 0) in the presence or absence (Ctl) of fignl1ΔN or fignl1ΔN-S329D. (L) Schematic of the experimental procedure employed for *in vivo* rescue experiments in retina-specific fignl1 conditional mutants using fignl1ΔN-GFP or fignl1ΔN-S329A-GFP. IUE: *In Utero* electroporation. (M) Z-projections of 3D light-sheet images from cleared E18.5 embryo heads electroporated in one retina with *Fignl1* sh-RNA;tdTomato and fignl1ΔN-GFP or fignl1ΔN-S329A-GFP constructs and immunolabelled with anti-DsRed antibody (detection of tdTomato). Arrows indicate aberrant ipsilateral projections and asterisk highlights a rescue effect. Dashed and dotted lines indicate the eye and midline positions, respectively. Insets show the electroporation patterns of the retinas. Scale bar: 500μm. (N) Violin Plot showing percentages of ipsilateral projections in embryos electroporated with *Fignl1* sh-RNA alone (n=7) or co-electroporated with *Fignl1* sh-RNA and fignl1ΔN-GFP (n=7) or fignl1ΔN-S329A-GFP (n=6), pooled from at least three independent experiments. (F, H, J, K, N): *p≤0.05; **p≤0.01, *** p≤0.001; **** p≤0.001. ns: non-significant. One-Way ANOVA with Bonferroni’s post-test (F, N), Kruskal–Wallis ANOVA with Dunn’s post hoc test (H, J-K). Horizontal solid and dotted lines in violin plots indicate the median ± the 1st and 3rd quartiles, respectively.

To test this hypothesis, we investigated whether disrupting fignl1/RAD51 binding alters Wnt5a-induced contralateral RGC axon repulsion *in vitro*. To do so, we performed rescue experiments in fignl1-deficient contralateral retinal explants with fignl1ΔN or the RAD51-binding deficient F304E/A307E variant.^17^ Fignl1ΔN fully restores the responsiveness of fignl1-depleted contralateral RGC axons to Wnt5a by inhibiting F-actin protrusions, bringing collapse rates back to control levels (50% of axons; Figures 6G and 6H versus Figures 4N and 4O). In contrast, the RAD51-binding-deficient F304E/A307E mutant fails to rescue this phenotype, with only 20% of growth cones collapsing, comparable to the basal level observed in PBS-treated cultures (Figures 6G and 6H). This phenotype mirrors the ones obtained with PKC dominant negative (PKC-DN) and fignl1ΔN PKC phospho-resistant (fignl1ΔN-PKC-7A) variants (Figures 5D and 5E, Figure 5J and 5K). Collectively, these results pinpoint the fignl1/RAD51 complex as a pivotal Wnt5a/PKC signalling integrator required for retinal axon guidance.

### RAD51-dependent PKC phosphorylation of fignl1 at S329 drives Wnt5a-evoked RGC axon midline crossing by inhibiting linear F-actin nucleation

To further validate this model, we next investigated whether phosphorylation of fignl1 at residue S329 by PKC – which is dependent on fignl1 scaffolding by RAD51 – is (i) sufficient to fine-tune fignl1’s cytoskeletal activities *in vitro* and (ii) drives RGC axon midline crossing. We found that PKC-mediated phosphorylation of fignl1 at S329 dampens fignl1 microtubule-depolymerizing activity while enhancing its localisation to the actin cytoskeleton *in cellulo* (Figures S5H-5J) as the fignl1ΔN-PKC-7D variant does (Figures 5F-5H). Consistent with this, *in vitro* reconstitution systems in which linear and branched actin networks were co-polymerised, revealed that phosphorylation of fignl1 at Ser329 is sufficient to selectively potentiate its inhibitory activity on linear F-actin nucleation (Figures 6I-6K). Finally, *in vivo* rescue experiments using *in utero* electroporation, confirmed the relevance of this phosphorylation event for retinal axon midline crossing (Figure 6L and Video S4). While the expression of fignl1ΔN significantly reduces the percentage of aberrant ipsilateral projections induced by retina-specific fignl1 knockdown, overexpression of the phospho-resistant fignl1ΔN-S329A variant fails to rescue this midline guidance phenotype (Figures 6M and 6N and Video S4). Altogether, these results demonstrate that the fignl1/RAD51 complex functions as a key Wnt5a/PKC signalling integrator, with RAD51 acting as scaffold that enables PKC-mediated phosphorylation of fignl1 at S329, thereby driving retinal axon midline crossing via inhibition of linear F-actin nucleation.

### Differential Rad51 and Fignl1 gene expression in ipsilateral vs. contralateral RGCs drives divergent axonal responses to Wnt5a

Ipsilateral RGC (iRGC) axons are also repelled by Wnt5a^32^ but appear more sensitive than contralateral (cRGC) axons, responding at lower concentrations (200 ng.mL^−1^ for iRGC^32^ *vs* 300 ng.mL^−1^ for cRGC; Figure S6B) and for longer durations (>1h for iRGC^32^ *vs* <30 min for cRGC and Figures S6A). To explore whether the Wnt5a integrator complex RAD51/fignl1 contributes to these differences, we analysed bulk RNA-seq data from E16 mouse cRGCs and iRGCs.^53^ Both *Fignl1* and *Rad51* transcripts are significantly enriched in iRGCs compared to cRGCs (Log_2_ fold-changes of 2 and 2.5, respectively), similar to established iRGC markers (*e.g*., Zic2, Epha2; Figures 7A and 7B). These gene expression differences may partly account for the heightened sensitivity of iRGCs to Wnt5a. To test this hypothesis, we assessed whether reducing fignl1 levels in iRGC – using *Fignl1* heterozygous mutants– would shift iRGC axonal responsiveness to Wnt5a toward a “cRGC-like” behaviour *in vitro*. Collapse assays performed on cultured ventro-temporal (iRGC-enriched) *versus* central (cRGC-only) retina explants revealed that *Fignl1* haploinsufficiency attenuates iRGC sensitivity to Wnt5a. *Fignl1* heterozygous mutant iRGC axons (Cre+^Ret^; *Fignl1^Fl/+^)* fail to respond to low Wnt5a concentrations (200 ng.mL^−1^), thereby phenocopying cRGC behaviour, whereas their temporal responsiveness at higher doses remains unchanged (iRGCs are still responsive after 1 h at 300 ng.mL^−1^, unlike cRGCs) (Figures 7C-7E). Together, these findings indicate that differential expression of *Fignl1* may contribute to the divergent sensitivity of iRGC and cRGC axons to midline-derived Wnt5a gradient and may limit the ability of iRGC axons to reach the midline.

**Figure 7:**
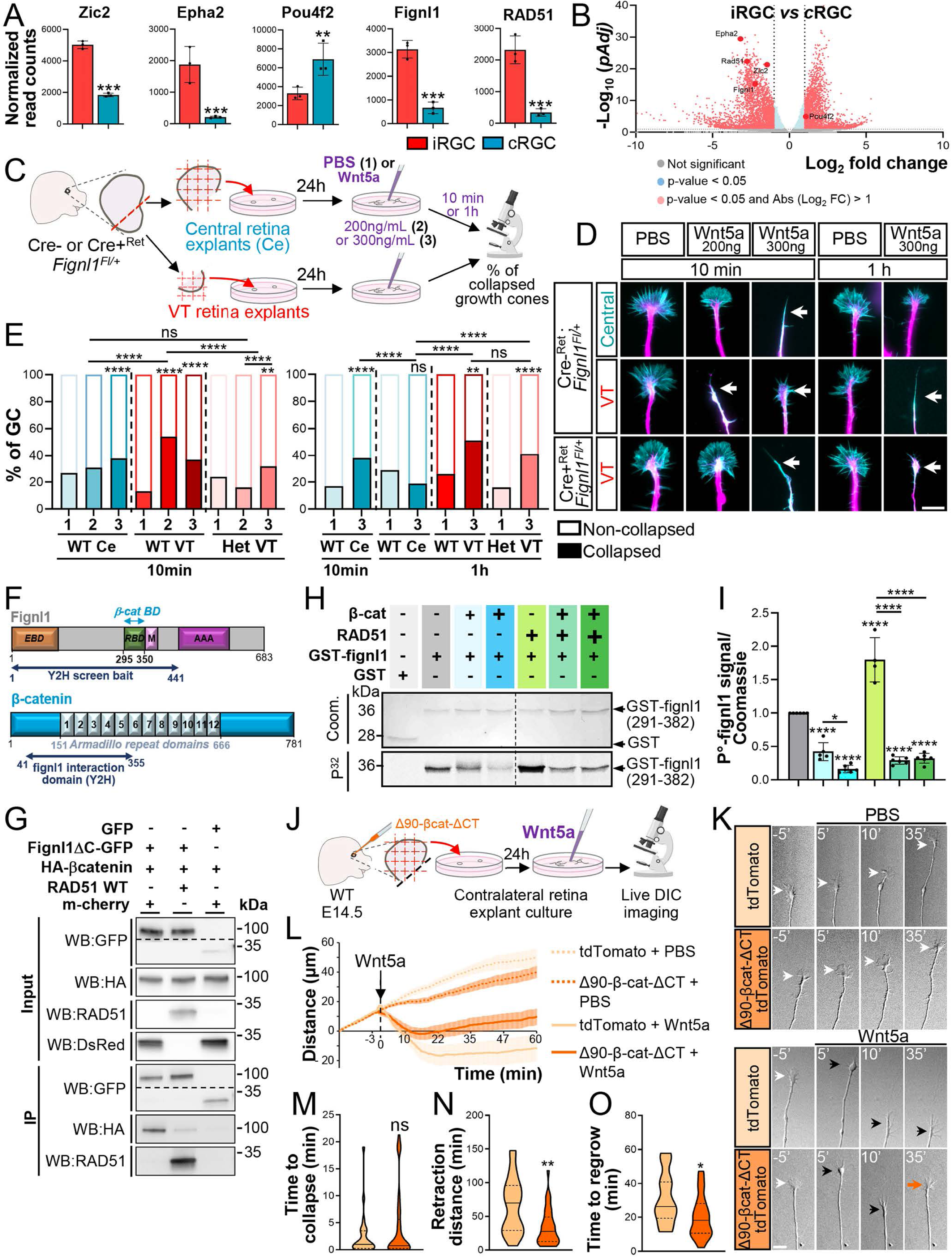
Fignl1/RAD51 expression levels and competition with β-catenin dictate the distinct responsiveness of ipsilateral and contralateral axons to Wnt5a. (A) Transcriptomic analysis of bulk-RNAseq data from Fernández-Nogales et al. (2022). Graphs show mean normalized read counts ± SEM for *Fignl1* and *Rad51* transcripts (Log_2_ fold changes of 2 and 2.5, respectively), and transcripts of iRGC (*Zic2*, *epha2*) or cRGC (*Pou4f2*) markers. Dots and squares represent biological replicates. Statistical significance is based on DESeq2 Wald test with adjusted p-values (pAdj). (B) Volcano plot showing the enrichment of *Fignl1* and *Rad51* transcripts in mouse iRGC vs cRGC at E.16. (C) Schematic of the experimental workflow used to assess the effect of fignl1 expression level on iRGC and cRGC axon responsiveness to Wnt5a. (D) Representative TIRF images of RGC growth cones from control (Cre-^Ret^; *Fignl1^Fl/+^*) or fignl1 heterozygous mutant (Cre+^Ret^; *Fignl1^Fl/+^*) ventro-temporal (iRGC enriched) or central (cRGC only) retina explants treated for 10 min or 1h with PBS or 200 or 300 ng.mL^−1^ Wnt5a and stained with tyrosinated tubulin antibody (magenta) and phalloidin (cyan). Arrows indicate collapsed growth cones. Scale bar: 10μm. (E) Bar graphs showing the percentage of collapsed growth cone quantified from at least 50 axons per VT condition and 275 axons per central retina explant (non-VT) condition. WT: wild-type, VT: ventro-temporal, Ce: central; **1**: PBS; **2**: Wnt5a 200 ng.mL^−1^; **3**: Wnt5a 300 ng.mL^−1^. (F) Schematic of mouse fignl1 and β-catenin domain organization. Dark blue arrows summarize the results of the yeast two-hybrid assay (Y2H). EBD: EB-binding domain; M: MTBD; RBD: RAD51 binding domain; AAA: ATPase domain; β-cat BD: β-catenin binding domain (identified Figure. S7C). (G) Western blot analysis of co-immunoprecipitation assays performed with a GFP antibody from protein extracts of COS-7 cells expressing HA-β-catenin, fignl1-GFP, or GFP alone, and RAD51 or m-Cherry (control). WB: Western blot; IP: immunoprecipitation. (H) *In vitro* PKCα kinase assays using 0.2 μM GST-fignl1(291–382) fragment as substrate in the presence or absence of 0.2 (**+**) or 0.4 μM β-catenin (+), and 0.4 (+) or 0.8 μM (+) of RAD51. (I) Quantification of phospho-fignl1 signal normalized to fignl1 Coomassie staining and expressed relative to fignl1 assayed in the absence of RAD51 and β-catenin (set to 1). Data are from at least four independent experiments. Dots indicate biological replicates. (J) Schematic of Wnt5a collapse assays performed in living cRGC axons electroporated with a non-degradable cytoplasmic β-catenin variant (Δ90-β-cat-ΔCT_T2A_tdTomato) or tdTomato. Axons were imaged for 75 minutes (1 frame/15 s). PBS or Wnt5a (300 ng.mL^−1^) was added 15 minutes after the start of imaging. (K) Representative DIC images acquired 5 min before and 5, 10, 35 minutes after cRGC axon exposure to PBS (Sham; top) or Wnt5a (300ng.mL^−1^; bottom). White, black and orange arrows indicate growing, collapsed, and prematurely regrowing axons, respectively. Scale bar: 10 μm (L) Traces showing the average directed growth length (μm) of axons analysed in (K): tdTomato + PBS (n=22), Δ90-β-cat-ΔCT PBS (n=17), tdTomato + Wnt5a (n=31) and Δ90-β-cat-ΔCT + Wnt5a (n=39), pooled from at least three independent experiments. (M-O) Violin plots showing the elapsed time between Wnt5a stimulation and growth cone collapse (M), the retraction distance (N), and the interval between growth cone collapse and regrowth (O), measured for Wnt5a-stimulated cRGC axons overexpressing tdTomato (n=25 for panels M-N and n=16 for panel O) or Δ90-β-cat-ΔCT -tdTomato (n=33 for panel M-N and n=25 for panel O). (E, I, M-O) *p≤0.05; **p≤0.01, **** p≤0.001. ns: non-significant. Chi2 test (E), One-Way ANOVA with Bonferroni’s post-test (I), Mann-Whitney test (M-N), and Unpaired t-test (O). Horizontal solid and dotted lines in violin plots indicate the median ± the 1st and 3rd quartiles, respectively.

### Competitive interaction between β-catenin and RAD51 for fignl1 binding promotes retinal axon midline crossing by temporally tuning their sensitivity to Wnt5a

To uncover the molecular mechanisms that regulate the divergent temporal sensitivity of cRGC and iRGC axons to Wnt5a, we used a global yeast-two hybrid proteomic screen to define the fignl1 interactome in the developing mouse brain (E16) (Figure 7F). This analysis identified the Wnt5a effector β-catenin^32^ as a direct fignl1-binding partner and potential regulator of RAD51/fignl1 functions, and mapped the fignl1-interaction domain of β-catenin to amino acids 41–355 (including armadillo repeats 1–6) based on four independent clones (Figure 7F). Co-immunoprecipitation assays validated this interaction while domain-mapping using a series of fignl1 deletion constructs showed that only the removal of the fignl1 RBD domain significantly impairs fignl1 interaction with β-catenin (Figure S6C). Consistent with this, we found that β-catenin, which accumulates in cRGC growth cones upon Wnt5a stimulation,^32^ preferentially binds the non-phosphorylated form of fignl1 (Figure S6D, 6E), competes with RAD51 for fignl1 binding in co-immunoprecipitation assays (Figure 7G) and still efficiently binds the RAD51-binding deficient F304E/A307E variant^17^ (Figure S6F, 6G). Using *in vitro* kinase assays, we further demonstrated that β-catenin inhibits PKC-mediated phosphorylation of fignl1 in a dose-dependent manner and impedes RAD51-mediated enhancement of this process (Figure 7H). Structural modelling supports this mechanism, showing that β-catenin (residues 150 to 360) binds fignl1 RBD, occluding both RAD51 and PKC docking and thereby preventing S329 phosphorylation (Figure S6H). Together, these findings suggest that β-catenin establishes a negative feedback loop that desensitizes cRGCs to Wnt5a. By competing with RAD51 and limiting PKC access to fignl1, β-catenin relieves fignl1-dependent inhibition of F-actin asseembly – enabling growth cones to exit the Wnt5a-induced paused state at the midline and resume contralateral growth.

To test this model, we overexpressed a cytoplasmic non-degradable β-catenin variant (Δ90-β-cat-ΔCter_T2A_tdTomato) in central retina explants and monitored their responsiveness to Wnt5a using live DIC imaging (Figure 7J). Forced β-catenin accumulation in contralateral RGC axons (but not that of tdTomato alone) induces their premature desensitization to Wnt5a – resulting in reduced axon retraction distance and shortened growth cone collapse-to-regrowth intervals (Figures 7K-7O). These behavioural defects closely phenocopied those of fignl1-deficient cRGC axons (Figure 4B and Figure 4E-4M). Strengthening these results, overexpression of the same β-catenin variant via *in utero* electroporation in the central retina impaired midline crossing of RGC axons,^32^ similar to fignl1 deficiency (Figure 1D-1F). Altogether, these findings support a model in which competitive binding of RAD51 and β-catenin to fignl1 promotes retinal axon midline crossing by temporally restricting Wnt5a responsiveness through control of Fignl1-dependent F-actin nucleation dynamics.

## Discussion

Our study reveals a previously unrecognized cytoplasmic role for the DNA repair complex fignl1/RAD51 in axon guidance in the mammalian brain. It uncovers how this complex translates axon guidance signals into temporal regulation of actin cytoskeletal remodelling to control axon decision-making at the midline.

Although several guidance pathways control ipsilateral or contralateral axon trajectories at the optic chiasm,^54–55^ the molecular mechanisms by which these signals are translated into cytoskeletal remodelling programmes governing the binary choice to cross or avoid the midline remains poorly understood. Among these pathways, non-canonical Wnt5a/β-catenin signalling has emerged as an atypical regulator influencing both ipsilateral and contralateral axon behaviours.^32^ Yet, how this guidance pathway generates distinct axon steering outcomes remains an unresolved question. Our findings indicate that both contralateral and ipsilateral RGC axons interpret Wnt5a as a repulsive cue rather than responding with opposite outcomes – growth promotion versus repulsion – as suggested by Morenilla-Palao and colleagues.^32^ Notably, we uncovered that their divergent behaviour emerges from intrinsic differences in the activation threshold and duration of a shared Wnt5a/PKC/RAD51/fignl1 repulsive programme, where PKC-mediated phosphorylation of RAD51-scaffolded fignl1 suppresses F-actin nucleation thereby maintaining growth cone sensitivity to Wnt5a (Figure S7). We further demonstrated that these divergent sensitivity and temporal responsiveness of cRGC and iRGC axons to Wnt5a are underpinned by differential expression levels *of Fignl1*/*Rad51* genes and differential kinetics of β-catenin-mediated feedback inhibition of RAD51/fignl1 binding and thereby F-actin nucleation inhibition. These findings provide a molecular framework explaining how distinct neuronal subtypes translate a common guidance signal into tailored cytoskeletal dynamic changes that direct specific axonal behaviours. These findings demonstrate that cytoskeletal dynamics can serve not only as an executor of guidance responses but also as an instructive determinant of axon trajectory choice.

This model reconciles our findings with those of Morenilla-Palao et al. by providing the molecular basis underlying the β-catenin-related guidance phenotypes they described.^32^ First, it accounts for the midline stalling of β-catenin-depleted cRGCs by proposing that loss of β-catenin-mediated inhibition of fignl1 activity prevents axons from exiting the paused state. Second, it explains the randomized ipsilateral versus contralateral trajectories induced by forced cytoplasmic accumulation of β-catenin in cRGCs^32^ as a consequence of premature fignl1 inhibition and Wnt5a desensitization (Figure. S7). Importantly, β-catenin-mediated temporal control of RAD51/fignl1 interaction may provide an integration hub for coordinating additional non-redundant key midline guidance pathways such as ephrinB2/EphB1 or Shh/Boc pathways.^56–57^ Notably, EphB1-mediated phosphorylation of β-catenin in iRGCs has been proposed to promote its dissociation from the F-actin/cadherin complex and its subcellular relocalization.^32^ This raises the possibility that ephrinB2/EphB1 signalling enhances iRGC sensitivity to Wnt5a by transiently relieving β-catenin-mediated feedback inhibition of RAD51/fignl1 control of actin dynamics, thereby promoting midline avoidance. Yet, this model, which pinpoints the RAD51/fignl1 complex as a key midline signalling hub integrating multiple guidance cues, remains to be directly tested. More intriguingly, the elevated expression of *Fignl1* and *Rad51* in iRGCs (Figure 6 and Fernández-Nogales et al., 2022^53^) appears independent of the ipsilateral fate determinant Zic2, unlike EphB1 and several Wnt receptor genes.^32^ This suggests that ipsilateral identity emerges from parallel Zic2-dependent and Zic2-independent transcriptional trajectories that ultimately converge to specify a unified guidance behaviour.

Beyond actin regulation, Wnt5a signalling has also been implicated in microtubule remodelling through a Ca²⁺/CaMKII/Tau-dependent pathway during contralateral growth of commissural axons.^58–60^ While inhibition of fignl1 microtubule-depolymerizing activity by Wnt5a/PKC signalling might contribute to growth cone pausing at the midline, Wnt5a/β-catenin-mediated relief of both fignl1-driven inhibition of F-actin nucleation and microtubule-depolymerization may operate in parallel with Tau-dependent microtubule reorientation and bundling.^59–60^ Together, these pathways may coordinate the large-scale remodelling of the growth cone cytoskeleton induced by Wnt5a, enabling exit from the pause state and resumption of growth in the contralateral tract.

Together with our previous work in zebrafish,^24–25^ this study reveals an evolutionary conserved role for the Meiotic Clade AAA+ ATPase fignl1 as a central player in neuronal circuit wiring and broadens its mechanistic landscape. In RGC axons, RAD51 binding to fignl1 directs fignl1 toward an F-actin remodelling activity required for axon decision-making at the midline. In contrast, in motor neurons, fignl1 interactions with end-binding proteins (EBs) or motor/adaptor complexes such as KIF1bβ/Bicd1, direct its activity towards the regulation of microtubule plus-end dynamics and dynein-dependent transport, respectively, two processes critical for motor axon guidance at choice points.^24,25^ These findings demonstrates that cytoskeletal regulators do not function as dedicated effectors of a specific guidance pathway but rather as versatile signalling hubs whose activities are reshaped by both the cellular context and upstream signalling inputs. Through differential interaction networks and post-translational modifications, factors such as fignl1 can be redeployed to control fundamentally different cytoskeletal processes. Such functional plasticity may constitute a key principle by which a limited repertoire of guidance cues generates the remarkable diversity of axonal behaviours underlying the assembly of complex neural circuits.

Yet, beyond its role in growth cone responsiveness, our findings reveal a compartment-dependent molecular versatility of the RAD51/fignl1 complex. In neuronal growth cones, RAD51 functions as an anchoring and scaffolding platform for fignl1, potentiating its F-actin inhibitory activity and thereby sensitizing growth cones to the Wnt5a repellent cue (present study). In contrast, in the nucleus, RAD51 nucleoprotein filaments serves as a substrate for fignl1 hexamers, which power RAD51 dissociation through ATPase-dependent conformational remodelling, thereby limiting RAD51 accumulation and preserving genome integrity.^20,22^ This striking functional shift suggests that RAD51/fignl1 activity is not intrinsically determined, but instead emerges from the local molecular environment, which regulates RAD51 partitioning between a nucleoprotein filament state and a monomeric scaffold state engaging fignl1.

To our knowledge, this study provides the first evidence that a Meiotic-Clade AAA ATPase directly regulates F-actin dynamics.^61^ Although members of this family, such as spastin, katanin and fidgetin, have been primarily characterized as microtubule-remodelling enzymes,^62–65^ we demonstrate that fignl1 inhibits both linear and branched F-actin assembly in vitro. These actin networks coexist in many cellular compartments, including neuronal growth cones, where their balance is critical for shaping cell morphology and behaviour.^66–68^ Accordingly, the ability of fignl1 to modulate their dynamics has important functional consequences. Consistent this idea, we show that fignl1 controls F-actin network homeostasis in both reconstituted systems and growth cones, preferentially suppressing linear over branched actin assembly. Notably, this activity is enhanced by PKC-dependent phosphorylation of fignl1 at Ser329 and is required for Wnt5a-induced axon repulsion. Although our data establish fignl1 as a key inhibitor of actin dynamics, the molecular basis of this activity remains to be fully elucidated. The differential requirement for ATP hydrolysis in fignl1-mediated inhibition of actin nucleation versus filament elongation (Figure 2), together with the presence of a predicted WH2 G-actin-binding domain^69^ in its sequence and its ability to bind F-actin, supports a dual mechanism. G-actin may serve as a substrate for ATPase-dependent remodelling by fignl1 hexamers as shown for tubulin^24^ and RAD51.^20, 22^ Such remodelling could generate a pool of assembly-incompetent actin monomers, thereby limiting filament formation. In parallel, fignl1 may directly associate with F-actin and restrict barbed-end elongation independently of ATP hydrolysis, similarly to capping proteins.^70,71^ This dual mode of action could explain the preferential inhibition of linear actin networks. In growth cones, enrichment of fignl1 within filopodia may simultaneously restrict the elongation of existing linear F-actin filaments while locally reducing the pool of actin monomers available for de novo filopodia formation. Interestingly, similar dual functionality has recently been reported for spastin, which combines ATP-dependent microtubule severing^62, 72^ and ATP-independent microtubule nucleation activities.^73,74^ This suggests that such dual regulatory mechanisms may represent a conserved feature within meiotic-clade AAA ATPases.

The discovery of an actin-remodelling activity for fignl1, together with the emerging role of nuclear F-actin in homologous recombination repair,^75–77^ raise the intriguing possibility that fignl1 may also contribute to the temporal regulation of nuclear F-actin during DNA repair.

Finally, our findings provide a potential mechanistic basis for the pathophysiology of RAD51-related Congenital Mirror Movements (CMM), a long-standing paradox linking mutations in a major DNA repair gene to defects in axon midline crossing.^8,12^ In this framework, our results suggest that CMM-causing RAD51 variants perturb the cytoskeletal functions of fignl1, thereby altering axon decision-making at the midline. Together with recent evidences implicating defective actin remodelling as a potential pathogenic mechanism in other genetic subtypes of CMM,^78,79^ our findings support a unifying model of CMM pathophysiology centred on cytoskeletal dysfunction.

More broadly, our study identifies it identifies temporal control of cytoskeletal dynamics by non-canonical Wnt5a signalling as an instructive mechanism that shapes cell behaviours. Notably, this principle may extend beyond axon guidance to other biological processes including polarized cell migration, with broad implications for embryonic morphogenesis, neuronal circuit assembly, and cancer metastasis. ^80–83^

## Material and Methods

### DNA constructs

The strategies used to generate all the DNA constructs used in this study are listed in Table S1 and are available upon request from the corresponding author.

DNA plasmids were prepared with the QIAquick Gel extraction kit (Qiagen, 28704) and the nucleospin plasmid kit (Qiagen, 12123). After preparation, plasmids were amplified with an endotoxin-free Maxi-prep kit (NucleoBond Xtra Maxi EF, Macherey-Nagel), sequenced (Eurofins) and stored at -20°C.

### Primary antibodies

The following primary antibodies were used in this study: mouse TUJ1 (MMS-435P, Biolegend), mouse Tyrosinated tubulin (clone TUB-1A2, T9028, Sigma), rabbit RAD51 (PC130, Millipore), mouse DsRed (632392, Takara), rabbit DsRed (632496, Takara), rabbit Fignl1 (17604-1-AP, Proteintech), rabbit Phospho-(Ser) PKC Substrate (#2261, cell signalling), rabbit RBPMS (PA5 31231, Invitrogen, 1:500), HA (clone 3F10, 11867423001, Roche), rabbit GFP (A-11122, Invitrogen), chicken GFP (GFP-1020, AVES Labs), mouse GAPDH (SC-32233, Santa Cruz Biotechnology Inc.).

### Fignl1 antibody production

A rabbit polyclonal fignl1 antibody (SF127) was generated against a mix of two synthetic peptides located in the mouse fignl1 protein (residues 230-244: GNTEKDTQSFPKTST and residues 286-300: APSKTEDRGRREDNS) by using the anti-peptide 28-day Speedy protocol from Eurogentec (Belgium). Final bleed was affinity purified.

### Animals and Ethical statement for mouse experimentation

All experiments were performed on mouse embryos at E.13.5, E14.5 or E16 collected from WT or *Fignl1^Fl/Fl^* pregnant females. Pregnant C57BL6/J WT mice used for *in utero* electroporation were purchased from Janvier Labs. All animal procedures were performed in accordance with the European legislation Directive 2010/63/UE and the French law (n° 2013-118 dated February 1, 2013) “on the protection of animals used for scientific purposes”. The Institute de la Vision (IdV) animal facility is approved to carry out animal experiments (arrêté préfectoral: DDPP A75-12-02). Mouse experimental procedures have been validated by the IdV local ethic committees (C2EA-05: Comité d’éthique en expérimentation animale Charles Darwin) and obtained the agreement of the Ministry for Higher Education and Research (IdV: APAFIS#22331-2019100814127972v4) and Charles Darwin Committee, agreement number 02241-02). Animals were delivered at the IdV facility at least 1 week before the surgery and were housed on a 12 h light/12 h dark cycle in temperature- and humidity-controlled environment (19–23 °C, 45–60%). Embryos from dated mattings (developmental stage stated in each section describing individual experiments) were not sexed during the experiments and the female over male ratio is expected to be close to 1.

### SABER- FISH

E16 embryo heads were fixed with 4% PAF and post-fixed for maximum 24 h, cryoprotected in 30% sucrose in PBS, embedded in 7.5% gelatin/10% sucrose in 0.12 M phosphate buffer, and frozen in isopentane at -50°C followed by cryosections of 20 µm.

#### Fluorescent in situ hybridization probe synthesis

Probe sets targeting mouse *Fignl1* and *Rad51* transcripts were designed using PaintSHOP35^84^ and ordered in oligonucleotide plate format from Integrated DNA Technologies (Leuven, Belgium). The sequences of the oligonucleotide probes, hairpins, HRP oligos used are listed in Table S2.

#### Primer exchange reaction

To extend FISH probes into PER concatemers, 100-μl reaction mix were prepared containing 1× PBS, 10 mM MgSO₄, 600 μM dNTP mix (including only dATP, dCTP, and dTTP; Invitrogen), 100 nM Clean. G hairpin, 600 U mL⁻¹ Bst LF polymerase (NEB, M0275M), 1.6 μM hairpin, and distilled water to a final volume of 90 μl. Reactions were incubated at 37 °C for 15 minutes, followed by the addition of 10 μl of a 10 μM primer solution. The mixtures were then incubated at 37 °C for 4 hours and subsequently heated at 80 °C for 20 minutes to inactivate the polymerase. To assess the efficacy of the extensions and lengths of the concatemers, samples were analysed on 1.5% TAE agarose gels for 60 minutes with Sybr Gold dye (Invitrogen, 10358492). Reactions were then purified and concentrated in 25 μL of distilled water using a MinElute kit (Qiagen, n28004).

#### Fluorescent RNA pSABER on Fixed Retina cryosections followed by immunostainings

Hybridizations were performed in hybridization chambers (Grace Bio-labs SecureSeal, GBL622514). Prior to hybridization, fixed retina sections were permeabilized with 1× PBS containing 0.1% (vol/vol) Triton X-100 for 10 minutes at room temperature to remove OCT. Samples were rinsed three times with 1× PBS (5 minutes each), incubated in 0.5% (vol/vol) H₂O₂ in 1× PBS for 10 minutes, rinsed twice with 1× PBS, and washed once with 1× PBS containing 0.1% (vol/vol) Tween (PBST). Sections were then incubated at 43 °C in a hybridization oven using a formamide hybridization wash buffer (40% formamide, 2× SSC, pH 7, 1% Tween-20) for 30 minutes. Probes (1.6 μg) were subsequently added to 160 μL of hybridization solution (40% formamide, 2× SSC, pH 7, 1% Tween-20, and 10% dextran sulfate (Sigma, D8906)). Hybridization with primary probes was carried out overnight, followed by washes in formamide hybridization wash buffer (2 × 30 minutes) and in 2× SSCT (0.1% Tween-20) (2 × 10 minutes). For secondary hybridization (fluorescent detection), samples were rinsed with 1× PBS and incubated in blocking solution (1% (wt/vol) BSA and 0.1% (vol/vol) Tween-20 in 1× PBS) for 30 minutes, followed by washing with 1× PBS (3 × 5 minutes). Secondary hybridization was performed by adding 100 nM HRP–oligonucleotide in 160 μL of a solution containing 2× SSCT, 0.1% (vol/vol) Tween-20, and 10% (wt/vol) dextran sulfate. Samples were incubated at 37 °C for 1.5 hours in the dark, washed with PBST (3 × 5 minutes) at 37 °C, and rinsed with 1× PBS (5 minutes). To develop the signal, Cy3 Tyramide (Tocris, 6457) was diluted 1:7500 from an 8 mM stock solution in a Tyramide reaction buffer (1× PBS, 0.0015% (vol/vol) H₂O₂) and applied to the samples for 10 minutes at room temperature. To quench enzyme activity, the samples were washed three times for 5 minutes each with a quenching solution containing 10 mM sodium azide and 10 mM sodium ascorbate in PBST, followed by two additional washes with PBST alone and subsequent DAPI staining.^85^

For subsequent immunostaining, sections were blocked with PBS-GT (0.2% gelatin, 0.25% Triton-X100 (sigma) in PBS) and incubated overnight at RT with anti-RBPMS antibody (PA5 31231, Invitrogen, 1:500) and Tag1 (AF4439, RD Systems,1:500). After several washes, sections were incubated 1 h at RT with an Alexa 647-conjugated donkey anti-rabbit antibody (ab150063, Abcam, 1/500). Sections were counterstained with DAPI and embedded in Mowiol.

### Immunohistochemistry on mouse brain sections

Series of 20µm-thick frontal mouse brain sections of E16 embryos (perfused with 4%PFA) were dried at room temperature for 15 minutes, dipped and heated in sodium citrate buffer (10mM sodium citrate, pH 6.0) for 5 minutes in the microwave (800W) for epitope retrieval. After several washes in PBS, sections were blocked in PBSGT (PBS 1X, 0.2% gelatin, 0.25% Triton X-100) for 2 hours at room temperature and incubated overnight at 4°C with the fignl1 antibody described above (purified SF127, 1:100), its corresponding pre-immune serum (PI127, 1:100) diluted in PBSGT. Sections were then washed in PBS and incubated for 2 hours at room temperature with the appropriate secondary antibody diluted in PBSGT. Finally, sections were washed in PBS and mounted in Mowiol (Calbiochem) and acquired with a fluorescence microscope equipped with an Apotome module (Axiovert 200M), the Axiocam MRm camera, and Axiovision software (ZEISS). Images were processed with ImageJ software.

### *In utero* electroporation

*In utero* electroporation was performed as previously described.^86^ Briefly, timed-pregnant mice (Janvier Labs) were delivered to the animal facility at least one week prior to the surgery to allow a minimum of 5 days adaptation. C57BL/6J pregnant mice were analgised pre-surgery with a subcutaneous injection of buprenorphine (Buprecare; 0,1mg.kg^−1^). After 20 minutes, mice were anesthetized deeply with 4% isofluorane for the induction and 2% for the maintenance. Midline laparotomy was performed, exposing uterine horns and allowing visualization of embryos. The left eye of E14.5 embryos was injected with a 2 µg.µL^−1^ DNA solution mix (containing the plasmids of interest diluted in PBS supplemented with FastGreen) using an elongated glass capillary (Harvard apparatus). The success of DNA injection was assessed using the green coloration of the eye. The eye was then electroporated with 5 pulses of 45 V during 50 ms every 950 ms (Nepagene electroporator). To target the central part of the retina (i.e. containing contralateral RGCs), the positive electrode (CUY650P5, Sonidel) was placed on the side of the injected eye.^87,88^ Following surgery, the incision site was sutured (4-0, Ethicon), and mice were injected with Meloxicam (5 mg.kg^−1^) for post-surgery analgesia. Pregnant mice were euthanized at E18.5 to collect the embryos.

### Whole mount immunostaining and tissue clearing

E18.5 embryos were harvested and the heads were fixed by immersion in a PBS/4% PFA solution overnight. The skulls were dissected out to expose the brain and samples were subsequently post-fixed for 2 h in PBS/4% PFA and prepared according to the iDISCO+protocol adapted from Belle et al., 2017 ^89^. Briefly, brains were dehydrated in successive baths of graded methanol concentration (40% MeOH, 60% MeOH, 80% MeOH, 100% MeOH) of 1.5 h each at RT. Samples were then transferred overnight at 4 °C in a methanol solution supplemented with 11% H2O2 (VWR, 216763) under light exposure to induce retina depigmentation. Samples were then rehydrated through successive PBS-methanol baths of decreasing methanol concentration for 1.5 hours each at room temperature (100% MeOH, 80% MeOH, 60% MeOH, 40% MeOH, PBS1X) and kept in PBS at 4 °C until proceeding to *in toto* immunostaining.

For *in toto* immunolabelling, E18.5 heads (*i.e.* brain and retinas) were permeabilized and blocked in PBS-GT solution (PBS1X supplemented with 0.5% Triton-X100, 0.2% gelatin and 0.1 g/L thimerosal) for 24 h at 37°C under agitation. Samples were subsequently incubated for 1 week at 37°C under agitation with rabbit anti-DsRed (for the detection of tdTomato; 1/1000, Takara Bio, lot #2103116) and/or chick anti-GFP primary antibodies diluted in PBS-GT solution. Samples were then washed several times in PBS-GT for 1 day and incubated for 2 consecutive days at 37 °C under agitation with the following secondary antibodies diluted in a PBS-GT and filtered through a 0.22 µm membrane filter: Alexa Fluor 594-conjugated Donkey anti-rabbit (1/500, Thermofisher, lot #2563838) and Alexa Fluor 647-conjugated Donkey anti-chicken (1/300, Jackson Immunoresearch, lot #148687). Heads were then washed over day in PBS-GT (at least 6 different baths), rinsed 2 times in 1X PBS and stored in the dark at 4 °C until clearing.

For clearing, samples were transferred overnight in a 20% MeOH solution before being gradually dehydrated through successive incubation in PBS-methanol baths of increasing methanol concentrations (40% MeOH, 60% MeOH, 80% MeOH and 2 X 100% MeOH) for 1 hour each at room temperature. Samples were then transferred overnight in a solution of 2/3 dichloromethane and 1/3 MeOH. The following day, they were incubated 30 min in a 100% dichloromethane bath prior being transferred in 100% benzyl ether (DBE) and stored until imaging.

### Light-sheet microscopy and data analysis

#### Image acquisition

Cleared heads were imaged in a DBE bath on a light sheet microscope (Ultramicroscope Blaze, Miltenyi Biotec, Germany) equipped with a 5.5MP sCMOS camera (2560×2160 pixels). The light-sheet, of a 4μm thickness, was generated by lasers at two different wavelengths (561nm, 639nm). 4X objective was used as magnification. To generate a 3D image set, the sample was scanned in LightSpeed mode with a 11% overlap. The images are then processed via Imspector Pro 7.5.3 acquisition software (Miltenyi Biotec).

#### 3D images stitching and Imaris conversion

3D tilled images were reconstructed using MACS® iQ View-3D Large Volume Analysis Software (Miltenyi Biotec GmbH, v.1.0.3). Stitched files were converted in Imaris files through ImarisFileConverter 10.2 (Bitplane) and then converted into a SyGlass (.syg) file. Optic nerve and tracts signal was segmented manually using SyGlass 2.2 software (and MetaQuest 2 VR headset) on Max Projection with the ROI tool with appropriate contrast and brightness settings. The optic nerve and eyes signal were segmented with separate masks on both RFP and GFP channels. Masked RFP and GFP signals were exported and converted back into an Imaris file using an in-house Python script. Images were captured using Imaris screenshot tool with fixed size as .tiff files.

#### Signal quantification

Quantification of the signal were done using the Imaris software. ‘Spots’ of a fixed diameter of 180 µm were placed on three vicinal points along each optic tract (s1,s2,s3), along with an adjacent Spot on the background signal (b), around 1200 µm from the optic chiasm. The Intensity Sum value of each Spot was extracted for the RFP signal. Each tract intensity was calculated as the average of the three measurements minus the corresponding background (((s1+s2+s3)/3)-b). The percentage of ipsilateral projections was calculated as follows: the background-corrected average ipsilateral signal divided by the sum of background-corrected average ipsi- and contralateral signals.

### Generation of the retina-specific fignl1 conditional mutant and genotyping

Fignl1 conditional mutants (*Fignl1^Fl/Fl^* ^21^) were generated and kindly provided by Frédéric Baudat (IGF, Montpellier) and crossed with *Dkk3*:Cre transgenic mice expressing the Cre recombinase in the retinal progenitor cells^49^ to generate *Fignl1* retina-specific conditional mutants (*Dkk3*-Cre+^Ret^; *Fignl1^Fl/Fl^*). Progeny of *Dkk3*:Cre; *Fignl1^Fl/+^*x *Fignl1^Fl/Fl^* were collected at E14.5 for primary culture of retinal explants or tissues sampling and genotyped as followed. For DNA extraction, tails, brain or retina of E14.5 embryos were lysed in NaOH (50mM; Sigma Aldrich, S8045-5009) for 15 minutes at 95°C, followed by centrifugation after addition of Trizma hydrochloride (1 M, pH 7.6) (Sigma Aldrich, T2788-1L). DNA (0.7 μL) was added to 24 μL of FIREPol® Master Mix Ready to Load (Solis Biodyne, 04-12-0125) and PCR amplified using different primer pairs:

- *Cre detection*: Cre Forward: 5’-TTCCCGCAGAACCTGAAGATGTTCG-3’ and Cre Reverse: 5’-GGGTGTTATAAGCAATCCCCAGAAATGC-3’;

- *Fignl1 WT and flox allele detection:* Fignl1 Forward: 5’-GGGATCAAACACTAGGGTTCA GGC-3’ and Fignl1 Reverse: 5’-GATACAGTCTTCAAGATTAAGGACAACC-3’;

*Fignl1 deleted allele detection:* Fignl1 Forward: 5’-GGGATCAAACACTAGGGTTCAGGC-3’ and Fignl1deleted Reverse: 5’-CGGGTTACGGTAGTTTACTCCC-3’.

### RNA extraction and RT-qPCR analysis

Total mRNA of E16 Cre-^Ret^; *Fignl1^Fl/Fl^* and Cre+^Ret^; *Fignl1^Fl/Fl^* retina or brain were extracted with Trizol according to the manufacturer’s instructions. *Fignl1* and *GAPDH* transcripts were reverse-transcribed and PCR-amplified using the SuperScript™ III One-Step RT-PCR System with Platinum® Taq (ThermoFisher Scientific, France) from 200 ng of RNA extracts using the following primers:

ms_Fignl1_RT_EX1-2_FOR: 5’-GGGAGCACAGCTCATCCTGTGC-3’

ms_Fignl1_RT_EX3_REV: 5’-GAGCTGGACGTCTCCATGTCAAAGG-3’

Ex4-5_GAPDH_FOR: 5’-GGAAGGGCTCATGACCACAGTCC-3’

Ex7-UTR_GAPDH_REV: 5’-GTCCAGGGTTTCTTACTCCTTGGAGG-3’

### Primary culture of retinal explants

Embryos were harvested at E14.5 and microinjected into both eyes with a DNA solution (containing plasmids of interest diluted in PBS1X supplemented with FastGreen) using tapered glass capillaries. Injected retinas were electroporated using two poring pulses (square wave, 175 V, 5 ms duration, with 50 ms interval) followed by four transfer pulses (40 V, 50 ms and 950 ms interpulse) with a Nepa21 Super Electroporator (NepaGene). Electroporated retinas were extracted from the eye and the ventro-temporal quadrant was removed and either discarded (for experiments included in Figure 2 to 6) or cultured separately (Figure 7). Contralateral or ventro-temporal retinal explants were cultured in suspension for 30 min at 37°C under 5% CO2 in a culture medium composed of DMEM/F12, (Gibco, 11039-047) 200 mM glutamine (Gibco, 25030081), 0.4 mg/mL BSA (Sigma, A1933-256), 25% glucose (Sigma/Merck, G8769-100ml) and 1% penicillin/streptomycin (Gibco, 15140-122). Retinas were then cut into 200μm-square explants using a tissue-chopper (McIlwan), plated onto 14 mm glass coverslips, 35 mm glass bottom dishes (Ibidi) or 4-well glass bottom slides (Ibidi), coated with 100 μg·mL^−1^ poly-lysine (Sigma/Merck, SLCF7563) and 20 μg·mL^−1^ Laminin (Sigma Aldrich, L2020-1MG). Retinal explants were cultured for 24 hours in the culture medium supplemented with 0.5% (w/v) methylcellulose (Sigma/Merck, M0512-1006) and B27 (1/50; Life Technologies) at 37°C under 5% CO_2_.

### Growth cone collapse assays combined live DIC Imaging or immunolabelling

Contralateral retinal explants (*i.e.* explants from retina depleted of the ventro-temporal part) were cultured as mentioned above and the responsiveness of contralateral RGC axons to Wnt5a was assessed 24 hours post-plating using live DIC imaging. At least, 30 minutes prior the imaging of retinal explants, 0.5 μL.mL^−1^ of HEPES was added to each Ibidi-dish. After selection of single axons with non-collapsed growth cones, imaging was performed using an inverted Eclipse Ti2 (Nikon) microscope, equipped with a 40× oil-immersion objective, a Prime95B camera (Photometrics) and the Metamorph software. Retinal explants were imaged at 37 °C in culture medium without phenol red and supplemented with 2 μM HEPES. Retinal explants were treated by manual pipetting with 300nM of Human/mouse Wnt5a (R&D Systems; 645-WN-010/CF) or the equivalent volume of PBS (control vehicle). Images were acquired in DIC every 15 seconds for a 15- and a 60-minute period (1/15 Hz) before and after the addition of the cue, respectively. Images were processed using the ImageJ software. Growth/retraction length and speed were measured using the Manual tracking plugin (ImageJ software). Only the axons exhibiting an active growth (≥10 μm; *i.e.* ≥40 μm·h^−1^) in the first 15 minutes before the addition of Wnt5a were included in the analysis. For collapse assay experiments conducted on lived contralateral explants electroporated with Δ90-β-cat-ΔCT_T2A_tdTomato constructs, a single fluorescence image was acquired in the red channel at the beginning of the DIC acquisition to identify the electroporated axons.

For fixed experiments, ventro-temporal (*i.e.* iRGC enriched) or non ventro-temporal explants (cRGC only) were treated for 10 min with 200 or 300 ng·mL-1 of recombinant mouse Wnt5a (R&D Systems) or PBS (control vehicle) diluted in prewarmed culture medium and fixed with a 4% PFA/4% Sucrose (w/v) solution for 20 min at 37°C.

### Immunolabelling on retinal explants

For immunostaining experiments involving retinal explants electroporated with GFP- or tdTomato-tagged proteins, explants were fixed in PFA4%/Sucrose4% for 15min at 37°C. They were subsequently permeabilized in PBS-Triton 0.1% (X-100, Sigma) for 7min, blocked in PBS-BSA3% -5% goat serum for 1h at RT and incubated overnight at 4°C with anti-tyrosined tubulin (1/2000, Sigma), GFP (1/1000, Invitrogen) or DsRed (1/1000, Takara) antibodies. After rinsing in PBS1X, cells were incubated for 1h at room temperature with the appropriate secondary antibodies and phalloidin probes (1/100; Invitrogen). Explants were rinsed and mounted on slides with Mowiol. Images were acquired using either a DM6000 epifluorescence microscope (Leica) with a x10 dry objective (N.A. 0.3) or x63 immersion objective (N.A. 1.4) and the Metamorph software (Molecular Devices) for axonal length and growth cone collapse quantifications. For P-domain surface analysis (Figure 4N-P) growth cones were acquired using an inverted Eclipse Ti2 (Nikon) coupled with a 100× oil-immersion objective, TIRF module (Gataca system) and Metamorph software (Molecular Devices). For the analysis of axonal growth, a mosaic of images covering the entire surface of the explant was reconstructed using the 2D Stitching plugin (Fiji). The 20 longest axons of each explant were measured using the ‘segmented line’ option of the same software. The mean obtained for each explant from three independent experiments were used for statistical tests. Quantifications of the growth cone P-domain area were performed using the Fiji software on at least 30 growth cones pooled from 3 independent experiments.

For fignl1 immunostaining experiments used to validate the fignl1 retina-specific conditional mutant, retinal explants were pre-permeabilised 24 hours post-plating in PHEM buffer (60 mM PIPES, 27 mM HEPES, 10 mM EGTA, 8 mM MgSO4; pH 6.9) containing 0.4% of glutaraldehyde and 0.2% Triton X-100 for 90 s at 37°C, then post-fixed with PHEM containing 3% glutaraldehyde for 15 min at 37°C. Samples were then rinsed in PBS, permeabilised with PBS-Triton 0.25% for 10 min and incubated in a solution of sodium borohydride (NaBH4 1mg/mL, Sigma) for 10 min. Explants were blocked in PBS-BSA 10% for 1h and incubated overnight at 4°C with fignl1 antibody (1/100; Proteintech). Explants were then washed in PBS and incubated for 1.5h at room temperature with an Alexa Fluor 488-conjugated goat anti-rabbit antibody (1/800, Fisher Scientific) and phalloidin-Atto647N (1/100, 65906, Sigma).

### STED super-resolution microscopy on retinal explants

Contralateral retina explants from *Dkk3*:Cre+^Ret^; *Fignl1^Fl/Fl^* and Cre-^Ret^; *Fignl1^Fl/Fl^*were pre-permeabilised 24 hours post-plating in PHEM buffer (60 mM PIPES, 27 mM HEPES, 10 mM EGTA, 8 mM MgSO4; pH 6.9) containing 0.4% of glutaraldehyde and 0.2% Triton X-100 for 90 seconds at 37°C, then post-fixed with PHEM containing 3% glutaraldehyde for 15 min at 37°C. Samples were then rinsed in PBS, permeabilised with PBS-Triton 0.25% for 10 min and incubated in a solution of sodium borohydride (NaBH4 1mg/mL, Sigma) for 10 min. Explants were blocked in PBS-BSA 10% for 1h and incubated overnight at 4°C with anti-tyrosinated-tubulin antibody (1/1000; Sigma). Explants were then washed in PBS and incubated for 1.5h at room temperature with a goat anti-mouse secondary antibody coupled to Alexa Fluor 594 (1/500, A11005, Fisher Scientific) and phalloidin-Atto647N (1/50, 65906, Sigma). After rinsing in PBS, explants were mounted on slides with ProLong Gold Antifade Mounting media (Fischer Scientific). Growth cones were imaged using on a LEICA SP8 STED3X microscope with a 93x glycerol objective (pixel size between 15 nm and 21 nm). A pulsed 775 nm depletion laser was used at 80% and 75% of its power to quench F-actin labelled with Phalloidin-Atto 647N and tyrosinated-tubulin labelled with Alexa Fluor 594, respectively. Images were acquired in sequential mode with a Z-step of 50 nm and then deconvoluted using Huygens Professional Deconvolution v14.10 (Scientific Volume Imaging). Quantifications of the growth cone surface, P-domain area, number of filopodia/growth cone perimeter, microtubule number and invasion length in the P-domain were performed on at least 30 growth cones per genotype pooled from 3 independent experiments using a home-made plugging (from Eric Denarier) in the Fiji software. The transition zone between the central and peripheral domain of the growth cone is defined by a line crossing the base (pointed end) of each actin filipodia.

### Live TIRF imaging on retinal explants

Contralateral retinal explants were co-electroporated with the following pairs of plasmids: pCX:LifeAct-mRuby and pORANGElock:msfignl1-PAM3-GFP-Cas9 (Fignl1 KI), pORANGElock:mRAD51-PAM2-GFP-Cas9 (RAD51 KI), fignl1WT-GFP, fignl1ΔRBD-GFP, fignl1 F304E/A307E-GFP or pCX:GFP diluted at 0.5 μg. μl^−1^ (pCX:LifeAct-mRuby) and 1.5 ug.μl^−1^ (fignl1 or RAD51 constructs) in PBS supplemented with FastGreen (Sigma; 60 ng μl^−1^). For live imaging, 2 mM HEPES at 37 °C was added to the retinal explant cultures at least 20 min before imaging. F-actin and Fignl1-GFP or RAD51-GFP dynamics were monitored in a thermostatic chamber set at 37 °C using an inverted Eclipse Ti2 (Nikon) coupled with a 100× oil-immersion objective (N.A. 1.49), TIRF module (Gataca system) and Metamorph software (Molecular Devices). The z-drift was minimised using the Nikon Perfect Focus correction system. The incidence angles of the lasers were calibrated to obtained a 190 nm deep excitation for each wavelength and images were acquired every 1 sec during a 5 min time period using a Prime95B camera (Photometrics). Fignl1, RAD51 and LifeAct-mRuby fluorescence intensity profiles in growth cones were estimated for each experimental condition along a 3-6 μm-long lane (ROI) traced in the growth cone P-domain and spanning at least two filopodia (Image Fiji software; NIH). For quantification of fignl1 WT and mutated/deleted variants colocalization with F-actin filopodia in growth cone, fignl1 and F-actin fluorescence were analysed along four filopodia and a percentage of filopodia showing significant fignl1 staining was estimated per growth cone for at least 10 growth cones per condition pooled from three independent experiments. Fignl1-GFP, RAD51-GFP and LifeAct-mRuby dynamics was measured along 4 filopodia per growth cones in 8 to 10 growth cones per condition from using the Kymograph pluggin the Fiji software software and a home-made macro designed by E. Denarier.

### COS-7 cell culture, transfection and treatment

COS-7 cells were cultured on glass coverslips or glass-bottom 35mmd-Ibidi dishes (for splitFAST experiments) in DMEM supplemented with 10% foetal calf serum (FCS) at 37°C under 5% CO2 and transfected with the different cDNA constructs by using Lipofectamine 2000 (Thermo Fisher Scientific) according to the manufacturer’s instructions with a DNA/Lipofectamine ratio of 1:1.5. COS-7 cells overexpressing HA_Msfignl1 were lysed in SDS lysis buffer (25 mM sodium sulfate, pH 7.2, 5 mM EDTA, and 1% SDS) supplemented with 1 mM PMSF and a cocktail of protease inhibitors (Roche). Protein extracts were analysed by western-blot.

For experiments investigated the impact of PKC activity on fignl1 microtubule-depolymerizing activity transfected cells were treated 4 hours post-transfection with 10μM of the pan-PKC inhibitor Gö6983 (Trocris Bioscience), 200nM of the Phorbol 12-myristate 13-acetate (PMA, Trocris Bioscience) or the equivalent volume of DMSO (control vehicle) and fixed 12 hours post-treatment. For WB analysis assessing the impact of PKC activity on fignl1 phosphorylation, cells were treated for 10 min or 1h with 10μM Gö6983 and 200nM PMA or an equivalent volume of DMSO before been harvested and lysed in RIPA buffer (50 mM Tris-HCl, pH 7.4, 150 mM NaCl, 20 mM EGTA, 1% NP-40, 0.5% sodium deoxycholate, and 0.1% SDS).

### Immunocytochemistry on COS-7 cells

Cells were fixed 24 h post-transfection in 4% PFA/4% sucrose for 10 min, permeabilized in PBST 0,1% (0,1%Triton X-100 in PBS) for 5 minutes, blocked in PBS/5%NGS and incubated overnight with the following antibodies: fignl1 (1:100, SF127), fignl1 pre-immune serum (1:100, PI127), HA (1 :100, 11867423001, Roche), GFP (1 :1000, A11122, Invitrogen), tyrosinated tubulin (1:3000, YL1/2, Sigma). After several washes in PBS, cells were incubated with appropriate Alexa Fluor-conjugated secondary antibodies (Invitrogen) and Alexa Fluor 488 or 647-conjugated phalloidin (1:150, Invitrogen) to stain F-actin. Images were acquired using a Leica DM6000 epifluorescence microscope and a 63× oil-immersion objective (N.A. 1.4). For fignl1 antibody validation, fignl1 and HA mean fluorescence intensity was measured in the nucleus of at least 30 transfected and non-transfected cells per conditions using the Image J software. Assessment of fignl1 wild-type or PKC phospho-variants microtubule depolymerizing activity was performed by measuring the mean tubulin fluorescence intensity in the cytoplasm of transfected cells normalised to the mean tubulin fluorescence intensity of non-transfected neighbouring cells. On the same cells, the mean fluorescence intensity profile of F-actin (phalloidin) and GFP-tagged fignl1 wild-type or PKC phospho-variants was evaluated along four different 5-µm-long lanes (ROI) positioned at location with significant actin cortex staining. Each lane was drawn from and oriented perpendicular to the cell membrane. Cells with at least 3 out of 4 fluorescence intensity plot profiles exhibiting significant overlap between phalloidin and GFP staining at the actin cortex were scored as “YES” for colocalization and the others as “NO”. Quantifications were performed on at least 50 cells per experimental conditions pooled from three independent experiments.

### SplitFAST experiments

*- In COS7 cells*: COS-7 cells were plated in 35mm Ibidi-dishes and transfected with the different splitFAST DNA constructs (Figure S6D-6F) using Lipofectamine 2000 (Thermo Fisher Scientific) according to the manufacturer’s instructions with a DNA/Lipofectamine ratio of 1:1.5. Ibidi-dishes were washed 24 hours post-transfection, with prewarmed PBS1X before adding 1.8 mL of serum-free DMEM/GlutaMAX supplemented with 2 μM HEPES and 10μM lime dye (Twinkle Factory, 480541). Cells were imaged at 37°C a Nikon Eclipse Ti2 TIRF microscope, equipped with a 100X immersion objective (N.A. 1.49) using the HiLo mode. Images were acquired with a Prime95B camera (Photometrics) controlled by Metamorph software. The z-drift was minimized using the Nikon Perfect Focus correction system. Binding between Fignl1 and its partners was assessed by measuring the mean fluorescence intensity of the splitFAST signal (green channel) in the cytoplasm using the Fiji software.

*- In Retinal explants*: The plasmids used for splitFAST experiments were diluted at 1.2, 0.7 and 0.5 μg.μL^−1^ respectively in PBS supplemented with FastGreen (Sigma; 60 ng.μL^−1^) and the same imaging parameters as for COS7 experiments were used. LifeAct-mRuby and Lime fluorescence intensity profiles in growth cones were estimated for each experimental condition along a 3-6 μm-long lane (ROI) traced in the growth cone P-domain and spanning at least two filopodia (Image Fiji software; NIH). A least 8 growth cones pooled from 3 independent experiments were analysed per conditions.

### Co-immunoprecipitation experiments

For co-IP assays, COS-7 cells were transfected with the appropriate constructs and lysed 24 hours-post-transfection in radioimmunoprecipitation assay (RIPA) buffer (50 mM Tris-HCl, pH 7.4, 150 mM NaCl, 20 mM EGTA, 1% NP-40, 0.5% sodium deoxycholate, and 0.1% SDS) supplemented with a cocktail of protease inhibitors (Roche). Co-IPs were carried out by incubating 500μg of pre-cleared protein extracts with 20μL of GFP-Trap® (Chromotek) agarose beads for 4 hours at 4°C. Beads were washed several times in RIPA buffer and resuspended in 2X Laemmli buffer. The proteins retained on the beads were analysed by Western blot.

All co-IP experiments involving the fignl1 protein were performed with fignl1ΔC constructs deleted of the C-terminal part (a.a. 442-683) to impede fignl1 oligomerization with the endogenous wild-type fignl1.

### Western blot analysis

COS-7 cells (5 μg) or Co-IP protein extracts were electrophoresed SDS-PAGE gel (4–15% precast polyacrylamide gel, Biorad) and transferred to nitrocellulose membranes (Trans-Blot Turbo Mini 0.2 µm Nitrocellulose Transfer Packs, Biorad). Immunoblotting was performed after overnight incubation at 4°C with the following primary antibodies: fignl1 (SF127, 1/2000), fignl1 pre-immune serum (PI127, 1/2000), HA (1:5000, Roche), GFP (1:8,000, Invitrogen), DsRed (1/10000, Takara), RAD51 (1/1000, Millipore), Phospho-(Ser) PKC Substrate (1/1000, Cell signalling), or GAPDH (1:20000, Santa Cruz Biotechnology Inc.). For some experiments GAPDH was used as the loading control. Immunostained proteins were visualised using appropriate peroxydase-labelled secondary antibodies (Jackson ImmunoResearch) and the ECL Prime chemiluminescent substrate (Cytiva). For *in cellulo* PKC kinase assays, the Phospho-(Ser) PKC signal in the IP fraction was estimated by quantifying band density normalized to GFP signal (ImageJ software) for each experimental condition in at least four independent experiments. For Co-IP assays assessing the binding between HA-β-catenin and GFP-tagged wild-type or fignl1 mutated variants, HA signal in the IP fraction was estimated by quantifying band density normalized to GFP signal (ImageJ software) for each experimental condition in at least three independent experiments.

### Recombinant protein production

#### Fignl1 and its variants

Recombinant fignl1ΔN (lacking amino-acids 1 to 125) and its mutant/deletion variants used for F-actin co-pelleting or nucleation/polymerisation assays were purified using the Strep(II)-Strep-Tactin affinity purification system as described previously.^90^ Briefly, HEK293T cells were transfected with Lipofectamine 2000 and harvested 24 or 48 hours post-transfection and lysed in cell lysis buffer (50 mM HEPES, 300 mM NaCl and 0.5% Triton X-100, pH 7.4) supplemented with protease inhibitor cocktail (Roche). Cell lysate was subjected to centrifugation at 14,800 rpm for 20 minutes at 4°C. The supernatant was incubated with Strep-Tactin Sepharose beads (GE Healthcare) for 45 minutes. Following incubation, beads were washed with lysis buffer without protease inhibitors 2 times and protein was eluted in 80 μL of elution buffer (50 mM HEPES, 150 mM NaCl, 1 mM MgCl2, 1 mM EGTA, 1 mM dithiothreitol (DTT), 2.5 mM d-Desthiobiotin and 0.05% Triton X-100, pH 7.4). Purified proteins were snap-frozen and stored at −80°C.

For *in vitro* kinase assays, pGEX-6P constructs encoding GST-tagged fragments of fignl1 (a.a 291-382) were transformed into *Escherichia coli* Rosetta (DE3) pLysS competent cells, and bacteria were grown at 37°C in Luria-Bertani medium supplemented with 100 g.mL^−1^ ampicillin until OD_600_ was ∼1.0. Expression was induced by adding 1 mM isopropyl 1-thio-β-galactopyranoside (IPTG) and incubating bacteria for 2 h at 37°C. Cells were harvested by centrifugation at 5 000 g for 20 min at 4°C and resuspended in ‘lysis buffer’ containing 50 mM Tris-HCl, pH 7.5, 300 mM NaCl, 0.5% [wt/vol] Triton X-100, 0.2 mg.ml^−1^ lysozyme, 1 mM dithiothreitol (DTT), 1 mM benzamidine, and 0.5 mM 4-(2-aminoethyl)benzenesulfonyl fluoride hydrochloride (AEBSF), and lysed on ice by ultrasonication. Lysates were clarified by centrifugation at 20 000 g for 20 min at 4°C, and supernatants were incubated for 1.5 h at 4°C with glutathione-Sepharose 4B beads (GE Healthcare) previously equilibrated in ‘lysis buffer’. Beads were washed 4 times in ‘high salt’ buffer containing 50 mM Tris-HCl (pH 7.5), 500 mM NaCl, 0.1% Triton X-100, 1 mM DTT, and twice in ‘low salt’ buffer containing 50 mM Tris-HCl (pH 7.5), 0.1 mM ethylene glycol-bis (2-aminoethyl ether)-N,N,N′,N′-tetraacetic acid (EGTA), and 1 mM DTT. Proteins retained on the beads were eluted by incubation in ‘low salt’ buffer supplemented with 50 mM reduced glutathione and 10% sucrose for 30 min at 4°C. Eluates were divided into aliquots and stored at -80°C.

#### Actin

Rabbit skeletal muscle actin was purified according to established protocols and labelled with Alexa Fluor 568 Succinimidyl Ester, following the procedure described by (Antkowiak et al., 2019).^50^

#### WASp, Arp2/3 complex and capping protein

S. cerevisiae WASp and capping protein were purified as described in Antkowiak et al. (2019).^50^ Briefly, the corresponding plasmids were overexpressed in E. coli Rosetta 2(DE3)pLysS cells, and proteins were purified via their His- and/or GST-tags using affinity chromatography. The Arp2/3 complex was purified according to the protocol described by Antkowiak et al. (2019),^50^ with the modification that bovine brain tissue was used as the source material instead of yeast.

#### mDia1 formin

The mDia1 construct was derived from the plasmid described in Jégou et al., 2013,^91^ with the addition of a TEV protease cleavage site.^50^ mDia1 was expressed in E. coli BL21 cells. Protein expression was induced with 0.7 mM IPTG at an OD of 0.6 and incubated overnight at 18°C. Cells were harvested by centrifugation, and the pellet was lysed, clarified by ultracentrifugation, and purified using a TALON cobalt affinity column. The protein was eluted with an imidazole gradient, concentrated using an Amicon filter, and dialyzed overnight against a dialysis buffer (20 mM HEPES pH 7.5, 200 mM KCl, 6% glycerol). The final protein preparation was stored at -80°C.

#### Profilin

Profilin (Pfy1) was expressed in E. coli BL21 cells transformed with an ampicillin-resistant pET-derived plasmid. Protein expression was induced with 0.5 mM IPTG at an OD of 0.6, and cells were harvested after 3 hours. The cells were lysed by sonication, and the protein was purified using HiTrap Q chromatography followed by Superdex-200 gel filtration. Pure fractions were concentrated and stored at -80°C.

### F-actin co-pelleting assays

G-actin (1 μM) and fignl1ΔN (0.4 µM) were incubated for 40 min at RT in an actin polymerization (AP) buffer containing 2 mM Tris pH 8.0, 0.2 mM ATP, 0.5 mM DTT, 0.1 mM CaCl2, 50 mM KCl, 1 mM MgCl2, 1 mM EGTA and 10 mM imidazole. Samples were centrifuged at 60,000 rpm in a TLA100.3 rotor for 1 h and analysed by SDS-PAGE. The amount of fignl1ΔN bound to F-actin recovered in the pellet was quantified from Coomassie-stained gels using ImageJ software. Ratios were calculated based on the average of at least three independent experiments.

### F-actin nucleation/polymerisation assays combined with TIRF microscopy

Perfusion chambers were prepared with glass slides and coverslips functionalized with polyethylene glycol (PEG)-silane as described previously.^92^ The chamber was perfused with (PLL)-g-PEG (2 kDa, 0.1 mg.mL^−1^ in 10 mM HEPES, pH 7.4) and washed with 1 % BSA in AP buffer. Actin was polymerized by adding 0.4 µM G-actin (containing 25 % of Alexa-568 labelled G-actin) with increasing concentration of fignl1ΔN in AP buffer containing 1 mg. mL^−1^ glucose, 70 µg.mL^−1^ catalase, 580 µg. mL^−1^ glucose oxidase and 0.3 % methylcellulose. Chambers were sealed and observed under an inverted microscope (Eclipse Ti, Nikon) equipped with ilas2 TIRF system (Roper Scientific), a CMOS camera (Prime 95B, photometrics) under Metamorph software (version 7.7, Molecular Devices). Samples were excited at 561-nm and observed using an Apochromat 100x oil-immersion objective (N.A., 1.49). Time-lapse imaging was performed during 30 minutes at RT, at 1 frame every 5 s with a 100 ms exposure time.

All image analyses were performed using the ImageJ software. Growth rates were calculated by measuring the length of actin filaments over time. The number of filaments was determined using automated segmentation at each time point (script available at https://github.com/EricDenarier/Fignl1).

### Linear and branched F-actin network assembly assays

Polystyrene microspheres (2 µm diameter, 2.5 % solids (w/v) aqueous suspension, Polysciences, Inc) were diluted 10 times in HK buffer (20 mM HEPES pH 7.5, 150 mM KCl) and incubated with 1 µM WASp, a nucleation-promoting factor that activates the Arp2/3 complex, for 30 min on ice. These microspheres are referred to as “WASp-coated beads” throughout the manuscript. Microspheres were then blocked with 1 % bovine serum albumin (BSA) for 15 min, washed and eventually stored on ice in HK buffer supplemented with 0.1 % BSA. The mDia1 formin, comprising only its FH1 and FH2 domains (1 µM), was coated on glutathione-coated microspheres (4.55 µm diameter, 0.5 % solids (w/v) aqueous suspension, Spherotech, Inc) with a similar protocol.

1% labeled actin was centrifuged at 60,000 rpm in a TLA100.3 rotor for 1 h and the actual protein concentration was checked with a Nanodrop spectrophotometer. To initiate actin network assembly, WASp- and mDia1-coated beads were incubated with G-actin and other proteins in a motility buffer (20 mM HEPES pH 7.5; 150 mM KCl; 6 mM MgCl2; 10 mM DTT; 2.5 mM ATP; 0.2 % methylcellulose 1500 cP and 2 % BSA)^50^. Standard optimal protein concentrations were: 8 µM G-actin, 8 µM profilin, 1 µM capping protein, 0.5 µM Arp2/3 complex and various concentrations of fignl1ΔN or mutated variants. Control conditions, when appropriate, contained only the fignl1ΔN purification buffer.

Images were acquired on a Zeiss AxioObserver microscope equipped with a 63x/1.4NA Oil Ph3 Plan-Apochromat objective and EMCCD (CoolsnapHQ2, Photometrics) camera. Images were acquired with Zen blue edition. All images were acquired using identical illumination intensity and exposure settings. Intensity values were measured as previously described.^50^

### *In vitro* kinase assays

The GST-tagged fragments of fignl1 (a.a 291-382) purified from *Escherichia coli* were incubated at a concentration of 200 nM in a 20-mL reaction mixture containing 50 mM Tris-HCl (pH 7.5), 10 mM magnesium chloride, 1 mM DTT, and 100 mM [g-^33^P]ATP (Hartmann Analytic; 10 mCi/reaction) in the presence of 25 nM recombinant active GST-PKCα (Division of Signal Transduction Therapy, University of Dundee, UK). After incubation for 40 min at 30°C, the reactions were terminated by the addition of 10 mL of SDS sample buffer. Proteins were resolved by SDS-PAGE, and Coomassie blue-stained gels were dried and subjected to autoradiography.

### Evolutionary conservation analysis

FIGNL1 orthologs from binocular vertebrates (M. musculus, H. sapiens, P. troglodytes, C. lupus familiaris, F. catus, M. mulatta, G. gallus) were aligned using Clustal Omega. Alignments were curated in Jalview and visualized with WebLogo. T303, T306, and S329 were strongly conserved, along with flanking basic residues, while S326 showed moderate conservation. This supports the functional relevance of these PKCα target sites.

### Homology modelling

The amino acid sequence of fidgetin-like 1 (fignl1) from Mus musculus (UniProt Q8BPY9) was retrieved from UniProt. Functional domains were annotated based on conserved AAA+ ATPase motifs and the C-terminal MT-binding domain (MTBD). The RAD51-binding region (residues 303-329), containing several putative phosphorylation sites, was mapped on the sequence.^93^ Structural templates were identified using HHpred and BLASTp against the Protein Data Bank, followed by manual alignment curation to preserve secondary structure and functional motifs. Multi-template models were built using Modeller (v9.x) following already published protocols.^94^ Models were scored with DOPE and QMEAN and validated using PROCHECK, MolProbity, and ProSA.^95^ For regions with low template coverage, structures predicted by AlphaFold (v3) were used to guide loop modelling and refinement.^96^ Structural comparison of homology and AlphaFold models was performed with TM-align and Discovery Studio focusing on the RAD51-binding region. Final hybrid models combined template-supported AAA+/MTBD domains with flexible loop conformations predicted by AlphaFold/modeler. Selected models were protonated, solvated (TIP3P, 0.15 M NaCl), minimized (10 000 steps), heated to 300 K, equilibrated (2 ns), and relaxed for 2 ns using NAMD (v2.14) and the CHARMM36m force field. Final structures were inspected and prepared for docking in Discovery Studio. In parallel, a homology model of β-catenin (residues 76–684) from Mus musculus was generated using Modeller (v9.x), based on structural templates identified by HHpred and the AlphaFold v3 prediction.^95,96^ Template alignments were manually curated to preserve the armadillo repeat architecture and inter-repeat interactions. The best model, as assessed by DOPE, QMEAN, and MolProbity scores, was refined through 2 ns molecular dynamics equilibration under the same NAMD/CHARMM36m protocol.^97^ The resulting β-catenin model was used for subsequent docking studies with fignl1 (residues 294-683).

### Protein–protein docking

Phosphorylation sites were predicted with NetPhos 3.1 and Scansite 4.0. Consensus PKC motifs [R/K]-X-S/T or [R/K]-X-X-S/T were identified at T303, T306, and S329, with intermediate scores at T325 and S326, and lower at S354 and S369. Structural mapping confirmed surface exposure of these residues, supporting their potential regulatory function. Docking between fignl1 and PKCα or RAD51 was performed using ZDOCK, focusing on residues 303-329 and MTBD. Top poses were clustered, ranked by score, and analysed with Discovery Studio refined poses protocol.^98^ PKCα docking identified T303, T306, and S329 as probable contact sites. RAD51 docking induced local rearrangements that increased solvent exposure of S326 and S329. Sequential docking generated ternary complexes (fignl1-PKCα-RAD51). Phosphorylation at T303, T306, T325, S326, S329, S354, and S369 was modelled using included Discovery Studio protocols. All docking solutions were visualized and analysed in Discovery Studio. All binary and ternary complexes were simulated with NAMD (v2.14), CHARMM36m, TIP3P water, and 0.15 M NaCl. After minimization a 2 ns equilibration was performed at 300 K. Additionally, protein–protein docking between fignl1 (residues 294-683) and β-catenin (residues 76-684) was carried out with ZDOCK using the previously generated homology models.^98^ The docking search was constrained around the RAD51-binding region of FIGNL1 and the armadillo repeat surface of β-catenin. Top-ranking poses were refined and rescored using Discovery Studio’s energy minimization and interface analysis modules. The resulting complexes were minimized and equilibrated (2 ns, 300 K) in explicit solvent using NAMD and CHARMM36m.^97^

### Yeast two-hybrid screen

The yeast two-hybrid screen was conducted by the Hybrigenics company. cDNA encoding the N-terminal part of the mouse fignl1 (a.a. 1 to 441) was used as bait to screen an E16 mouse embryonic brain cDNA library. A total of four independent clones containing β-catenin were isolated from this screen.

### Transcriptomic analysis

Raw FASTQ files were retrieved from the Gene Expression Omnibus (GEO) under accession number GSE184275.^53^ Reads were aligned to the mouse reference genome (GRCm39) using STAR (v2.7.9). Gene-level read counts were extracted using STAR’s built-in quantMode GeneCounts option and imported into our in-house R Shiny application EYEDVseq for downstream differential gene analysis (using DESeq2 (v1.40)) and graphical representations (IdV Cell and Tissue Phenotyping Platform). Genes with fewer than 10 total counts across all samples were filtered out in EYEDVseq. DESeq2 normalization and dispersion estimation were applied, and pairwise contrasts were computed using the Wald test with Benjamini–Hochberg correction. Significantly differentially expressed genes were defined as adjusted p value < 0.05 and absolute log2 fold change > 1. A volcano plot and bar graphs were generated from DESeq2 results using the EnhancedVolcano package (v1.18) and ggplot2 (v3.5), available within EYEDVseq.

## Statistical Analysis

All data were obtained from at least two to three independent experiments. Statistical significance of the data was evaluated using the non-parametric Mann-Whitney test when comparing two groups assuming non-Gaussian distributions. The Kruskal–Wallis ANOVA test with Dunn’s post-test and the One-Way ANOVA test with Bonferroni’s post-test were used when comparing more than two groups assuming non-Gaussian or Gaussian distribution, respectively. The Chi2 test was used to compare the distribution of different cell or axonal populations in different experimental conditions. Data distribution was tested for normality using the D’Agostino and Pearson normality test or Shapiro-Wilk normality test for Western-blot analysis for which the n number of values was smaller. For all tests, alpha was set at 0.05. All Statistical analyses were performed using GraphPad Prism 9.00 (GraphPad Software, San Diego, CA).

**Table 1.**
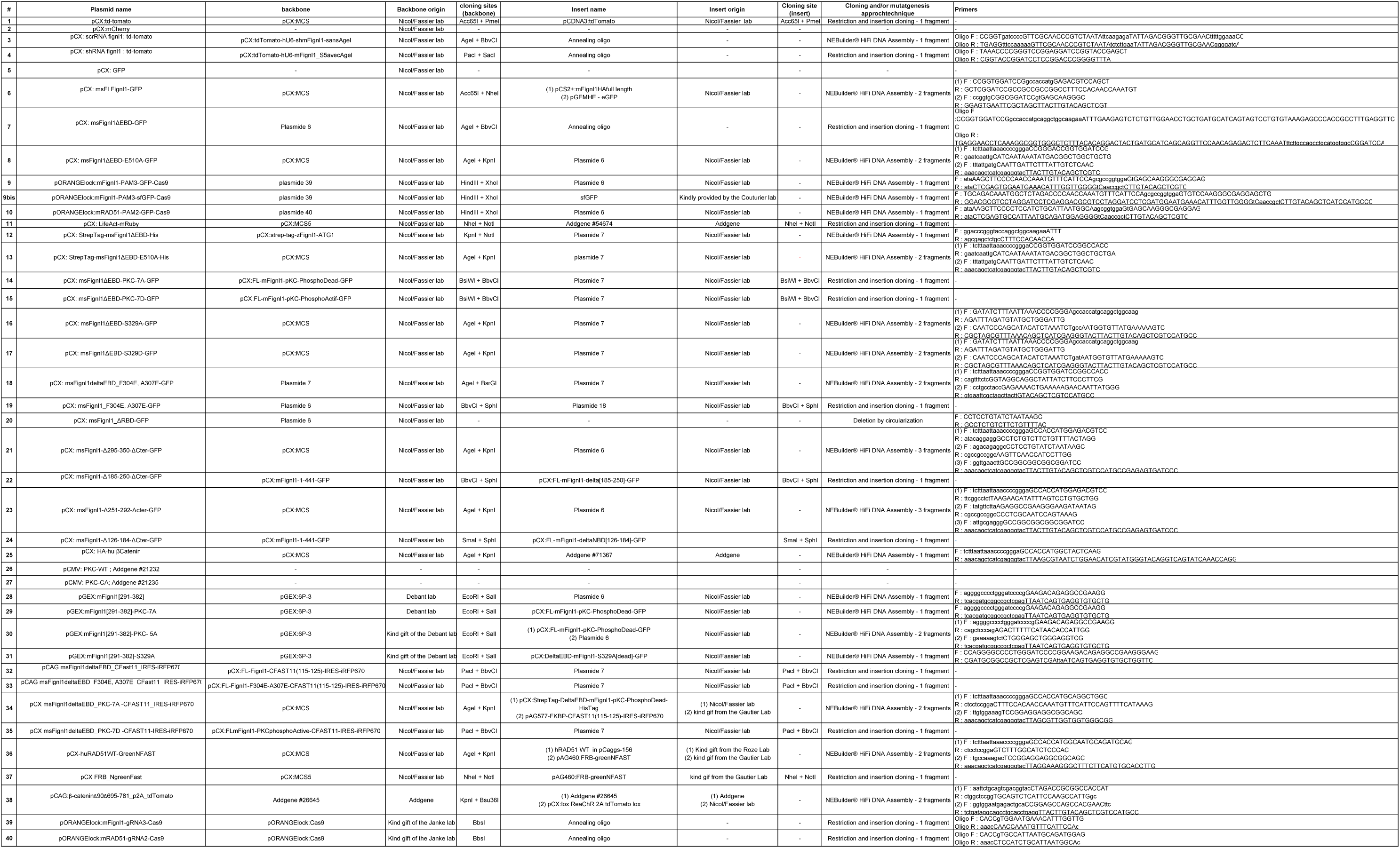

**Table 2.**
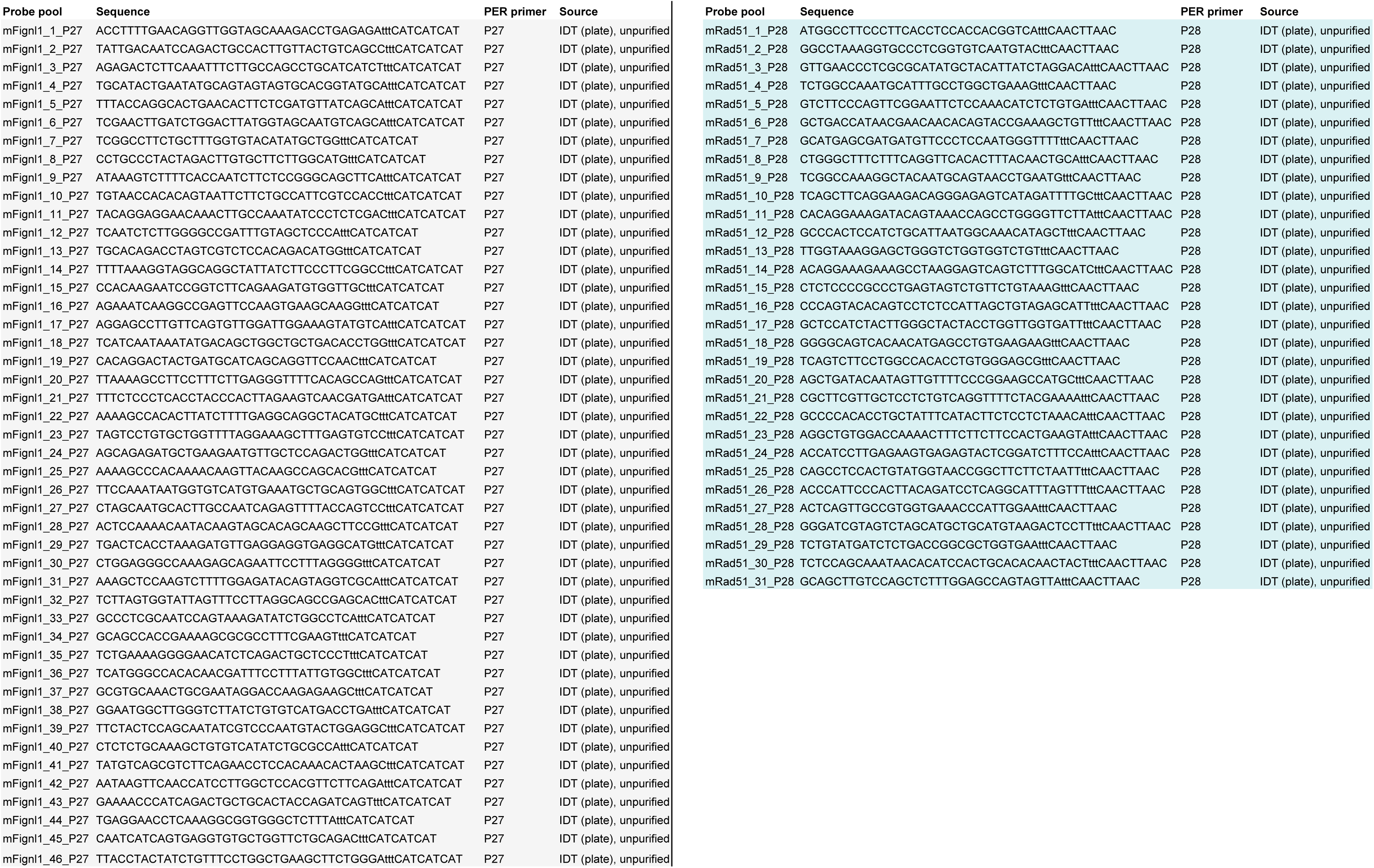

## Supporting information

Video S1: Fignl1 loss of function leads to an excess of ipsilateral projections, related to Figure 1.

Video S2: Fignl1 colocalises with F-actin-based filopodia in RGC growth cones, related to Figure 2.

Video S3: Fignl1 loss of function leads to the premature desensitization of contralateral RGC axons to the Wnt5a repellent cue, related to Figure 4.

Video S4: PKC-mediated phosphorylation of fignl1 S329 is required for RGC axon midline crossing, related to Figure 6.

## Data availability

All data supporting the findings of this study are available from the corresponding author on reasonable request.

## Acknowledgements

This work was supported by research grants awarded to C.F. from the French National Research Agency (ANR) (ANR-20-CE16-0019), UNADEV-AVIESAN funding, and the IHU FOReSIGHT program (IHU Reconnect). X.N. was supported by ANR grant ANR-20-CE13-0011 and the Fondation pour la Recherche Médicale (FRM) grant (MND202003011485). F.B. was supported by ANR grant ANR-17-426 CE12-0015. S.Z., S.B., and J.D. were recipients of PhD fellowships from the Ministère de l’Enseignement supérieur, de la Recherche et de l’Innovation (MESRI). S.Z. also received a one-year PhD fellowship from the Fondation de France. This work was also supported by the DIM C-BRAINS, funded by the Conseil Régional d’Ile-de-France. The NeurImag imaging facility is a member of the national infrastructure France-BioImaging supported by the French National Research Agency (ANR-24-INBS-0005 FBI BIOGEN).

We are grateful to Kim Charvet and Alain Chedotal (IdV/Sorbonne University) for kindly providing the *Dkk3*:Cre line and to Arnaud Gauthier for generously sharing the splitFAST plasmid backbones. We warmly thank Stéphane Fouquet and Nermine Saidi from the Imaging Facility of the Institut de la Vision (IdV/Sorbonne University) for their assistance with light-sheet microscopy and image analysis using Imaris and SyGlass software as well as Lydia Danglot from the IPNP Imaging Facility for granting us access to the STED microscope (LEICA SP8 3X system funded by the Leducq establishment) and for providing support through the allocation of engineering expertise to this project. We are also grateful to Frédéric Blond from the IdV Transcriptomics Facility for the import and coding of RNA-seq data, as well as to Anaïs Potey, Valérie Fradot, and Olivier Goureau from the IdV Cell Culture Facility for managing the cell culture room. We also thank the IdV and CFX mouse facility teams for animal handling, care, controlled breeding, and assistance with surgical procedures. Finally, we would like to thank Sylvia Soares, Christian Poüs, members from the I. Dusart and E. Roze labs as well as all members of the Nicol/Fassier team for their helpful comments throughout the study. We aree also grateful to Amélie Fréal for her careful editing of the manuscript, valuable feedback, and for her key advice on knock-in approaches in primary cell cultures.

## Author Contributions

C.F. conceived and supervised the study. C.F., S.Z., Y.Z., A.A., I.A. and J.B designed the experiments. S.Z., Y.Z., J.D., J.D., J.B., J.V., S.U., J.A., F.R., R.H., C.G.B., S.B., and C.F. performed the experiments. S.Z., Y.Z., J.D., J.B., I.A., A.A., and C.F. analysed the data and interpreted the results with A.A., A.D., and X.N. E.D. developed plugins for image analysis. A.Z. and F.B. provided the *Fignl1^Fl/Fl^* conditional mutant. P.B. acquired and deconvolved the STED images. N.P. performed all modelling analyses. X.G. performed the transcriptomic analysis with C.F. C.F. wrote the manuscript, which was critically reviewed and edited by all authors. All authors reviewed and approved the final manuscript.

---

**Figure S1.**
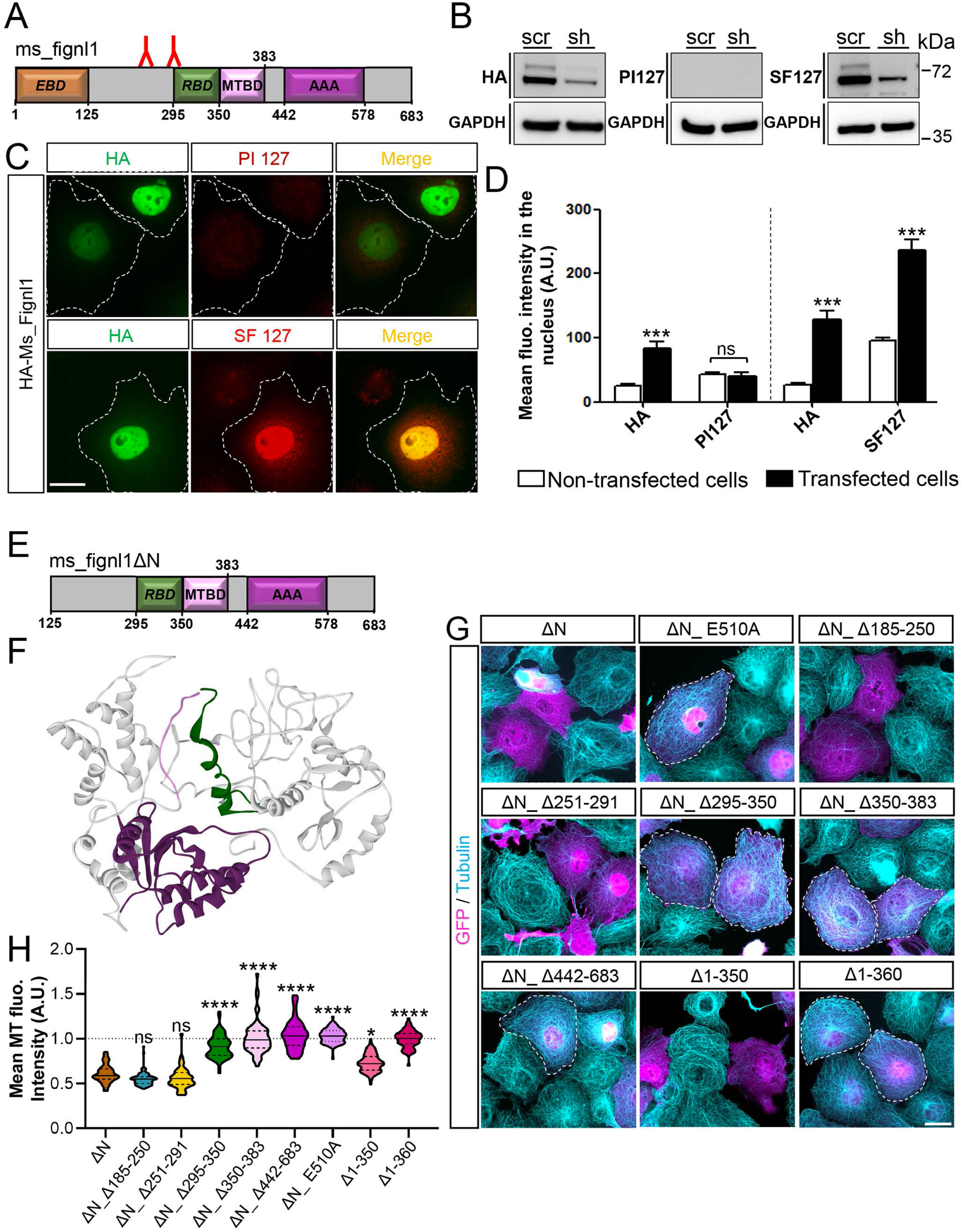
(related to Figure 1): Characterisation of mouse fignl1 MT-depolymerising activity and generation of mouse fignl1 antibody. (A) Schematic showing the locations of epitopes used to generate mouse fignl1 antibodies. EBD, RBD and MTBD: EB, RAD51 and microtubule-binding domains, respectively. AAA: ATPase domain. (B) Western blot analysis of protein extracts from COS-7 cells overexpressing HA-fignl1 and fignl1 or scramble sh-RNA using pre-immune serum (PI 127), anti-fignl1 (SF127), or anti-HA antibody. GAPDH was used as a loading control. (C) Representative epifluorescence images of COS-7 cells transfected with HA-fignl1 and immunolabelled with anti-HA antibody and pre-immune serum (PI 127) or anti-fignl1 antibody (SF127). Dotted lines indicate transfected cells. Scale Bar: 20 μm. (D) Bar graphs showing the mean fluorescence intensity of HA, PI 127 and SF127 staining measured in the nucleus of COS-7 cells overexpressing HA-fignl1 and immunolabelled with HA and PI 127 (n=30) or HA and SF127 (n=30) antibodies pooled from three independent experiments. (E) Schematic representation of mouse fignl1 structural domains. EBD, RBD and MTBD: EB, RAD51 and microtubule-binding domains, respectively. (F) Predicted 3D structure of mouse fignl1. The color code is the same as in panel E. (G) Representative epifluorescence images of COS-7 cells transfected with wild-type or ATPase-dead (E510A) fignl1ΔN constructs or domain-deletion variants of fignl1ΔN, and immuno-labelled with GFP and tubulin antibodies. Dashed lines indicate cells showing defective fignl1-mediated microtubule depolymerization. These data demonstrate that mouse fignl1 exhibits ATP-dependent microtubule-depolymerizing activity requiring both its microtubule-binding domain (residues 350-383) and its RAD51-binding domain (residues 295-350). Scale Bar: 20 μm. (H) Violin plots showing microtubule fluorescence intensity measured in COS-7 overexpressing fignl1ΔN (n=51), ΔN_ Δ185-250 (n=79), ΔN_ Δ251-291 (n=49), ΔN_ Δ295-350 (n=68), ΔN_ Δ350-383 (n=62), ΔN_ Δ442-683 (n=53), ΔN_ E510A (n=73), Δ1-350 (n=98) and Δ1-360 (n=64) and normalized to non-transfected cells from panel (G). Cells were pooled from three independent experiments. (D, H) *p≤0.05; ***p≤0.001, **** p≤0.001. ns: non-significant. Kruskal–Wallis ANOVA with Dunn’s post hoc test. Horizontal solid and dotted bars in violin plots indicate the median ± the 1st and 3rd quartiles, respectively.

**Figure S2.**
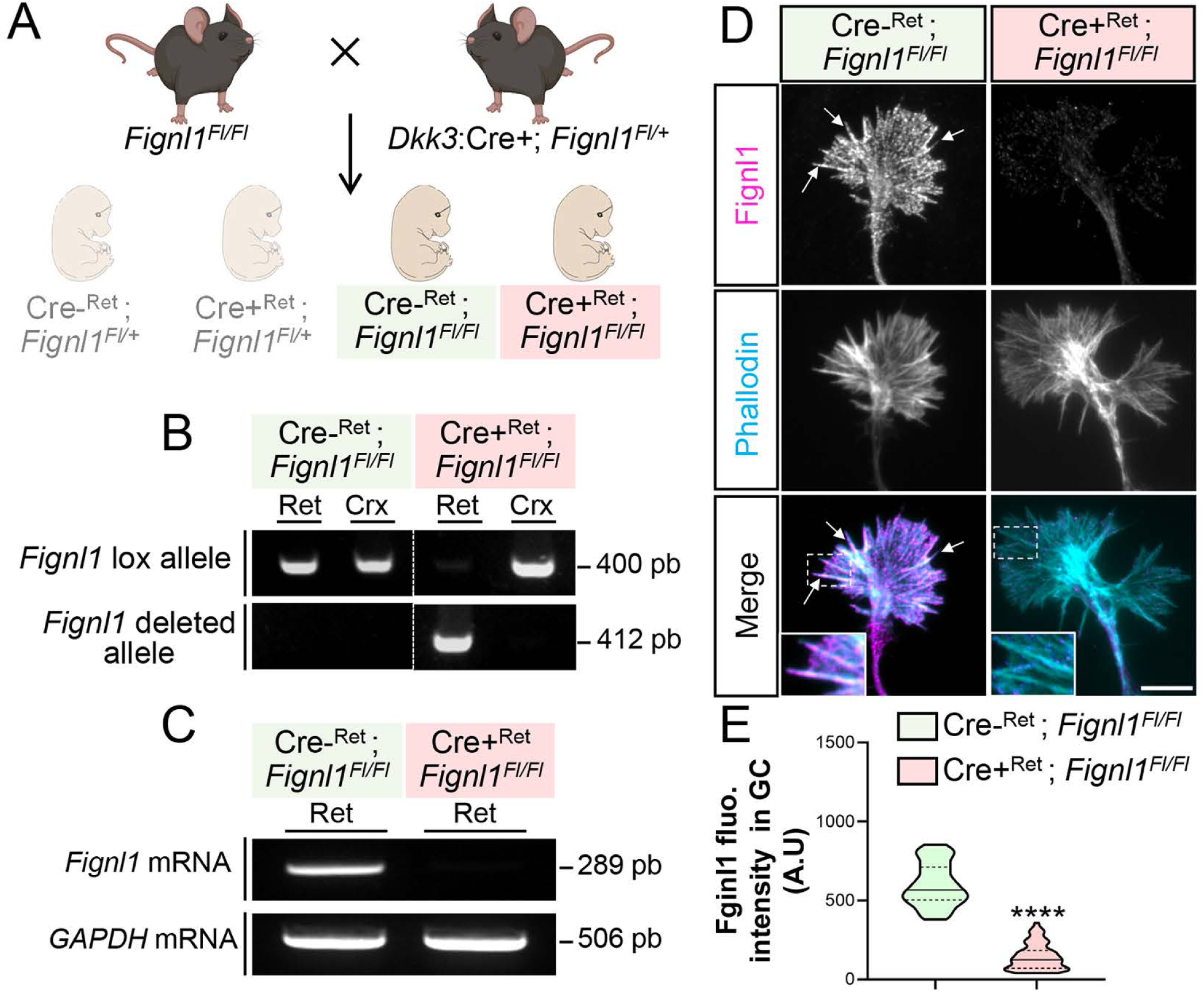
(related to Figures 3-7): Generation of a fignl1 retina-specific conditional mutant. (A) Schematic illustrating the strategy used to generate the retina-specific fignl1 conditional mutant. (B) DNA analysis of fignl1 gene editing efficiency and tissue specificity. (C) RT-PCR analysis of fignl1 knockout efficiency and tissue specificity. (D) Representative TIRF images of growth cones from control (Cre-^Ret^; *Fignl1^Fl/Fl^*) and fignl1-depleted (Cre+^Ret^; *Fignl1^Fl/Fl^*) retinal explants labelled with anti-fignl1 antibody (magenta) and phalloidin (cyan). Arrows indicate fignl1 co-localisation with F-actin-rich filopodia. Insets show higher magnifications of filopodia located in the boxed regions. Scale bar: 5 μm. (E) Violin plots showing the mean fignl1 fluorescence intensity in control (Cre-^Ret^; *Fignl1^Fl/Fl^*; n=51) and fignl1-depleted (Cre+^Ret^; *Fignl1^Fl/Fl^*; n=58) growth cones from (D)s. **** p≤0.001. Mann-Whitney test. Horizontal full and dotted lines indicate the median ± the 1st and 3rd quartiles, respectively.

**Figure S3.**
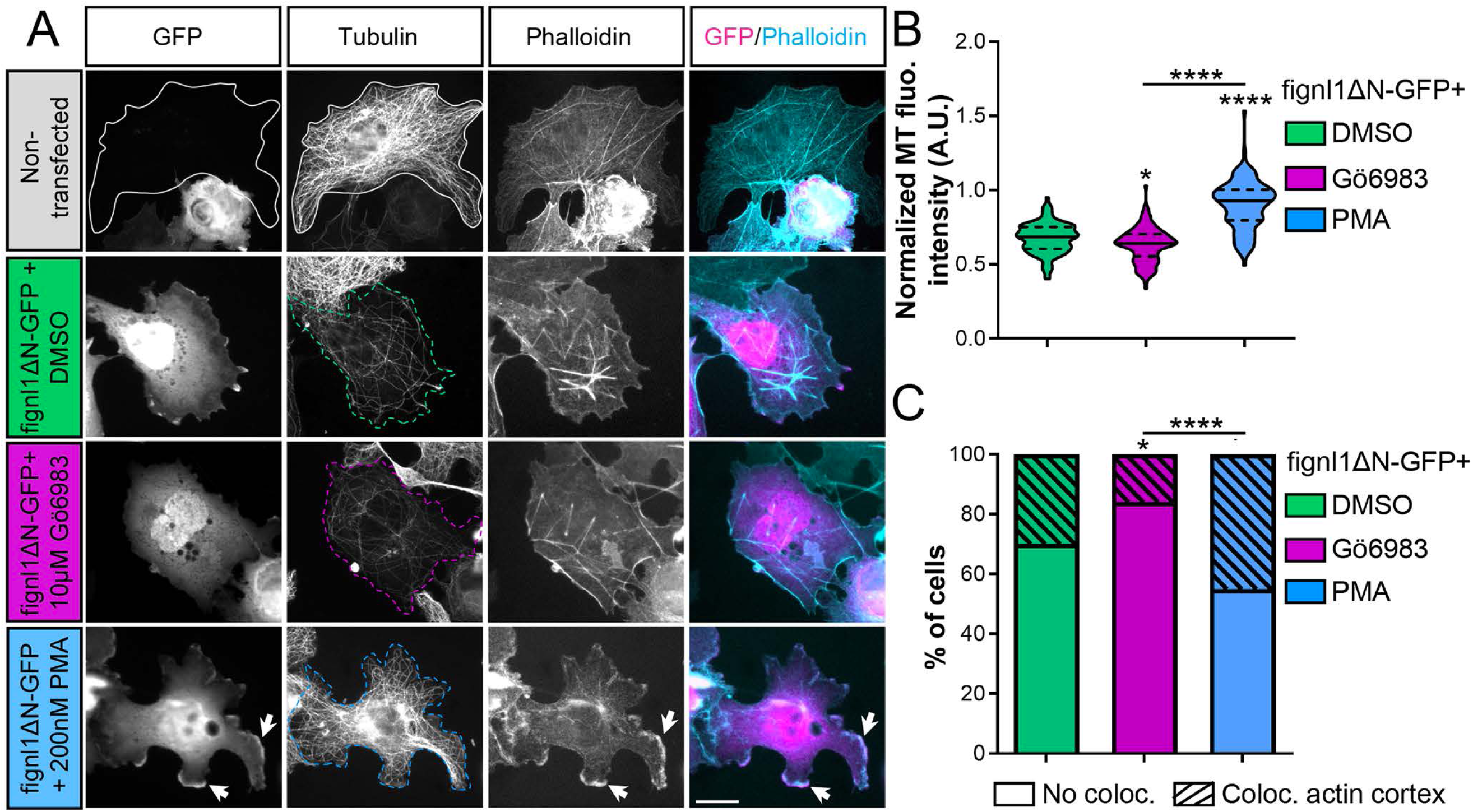
(Related to figure 5): PKC pharmacological modulations influence fignl1 cytoskeleton-related activities. (A) Representative epifluorescence images of COS-7 transfected or not with fignl1ΔN-GFP plasmid, treated for 12h with 10μM of Gö6983, 200nM of PMA, or DMSO (control), and labelled with anti-GFP (magenta), and anti-tubulin antibodies, and phalloidin (cyan). Full and dashed lines outline non-transfected and transfected cells, respectively. Arrows indicate fignl1ΔN-GFP colocalization with the cortical actin network. Scale bar: 20μm. (B) Violin plots showing microtubule fluorescence intensity in transfected cells normalized to non-transfected cells, quantified on Gö6983- (n=119), PMA- (n=112) or DMSO-treated (n=129) cells from (A) pooled from three independent experiments. Horizontal full and dotted lines indicate the median ± the 1st and 3rd quartiles, respectively. (C) Bar graph showing the percentage of fignl1ΔN-GFP-expressing cells treated with Gö6983 (n=119), PMA (n=112), or DMSO (n=129) that exhibit significant colocalization between fignl1 and cortical F-actin. (B-C) *p≤0.05; **** p≤0.001. One-Way ANOVA with Bonferroni post-test (B) and Chi2 test (C).

**Figure S4.**
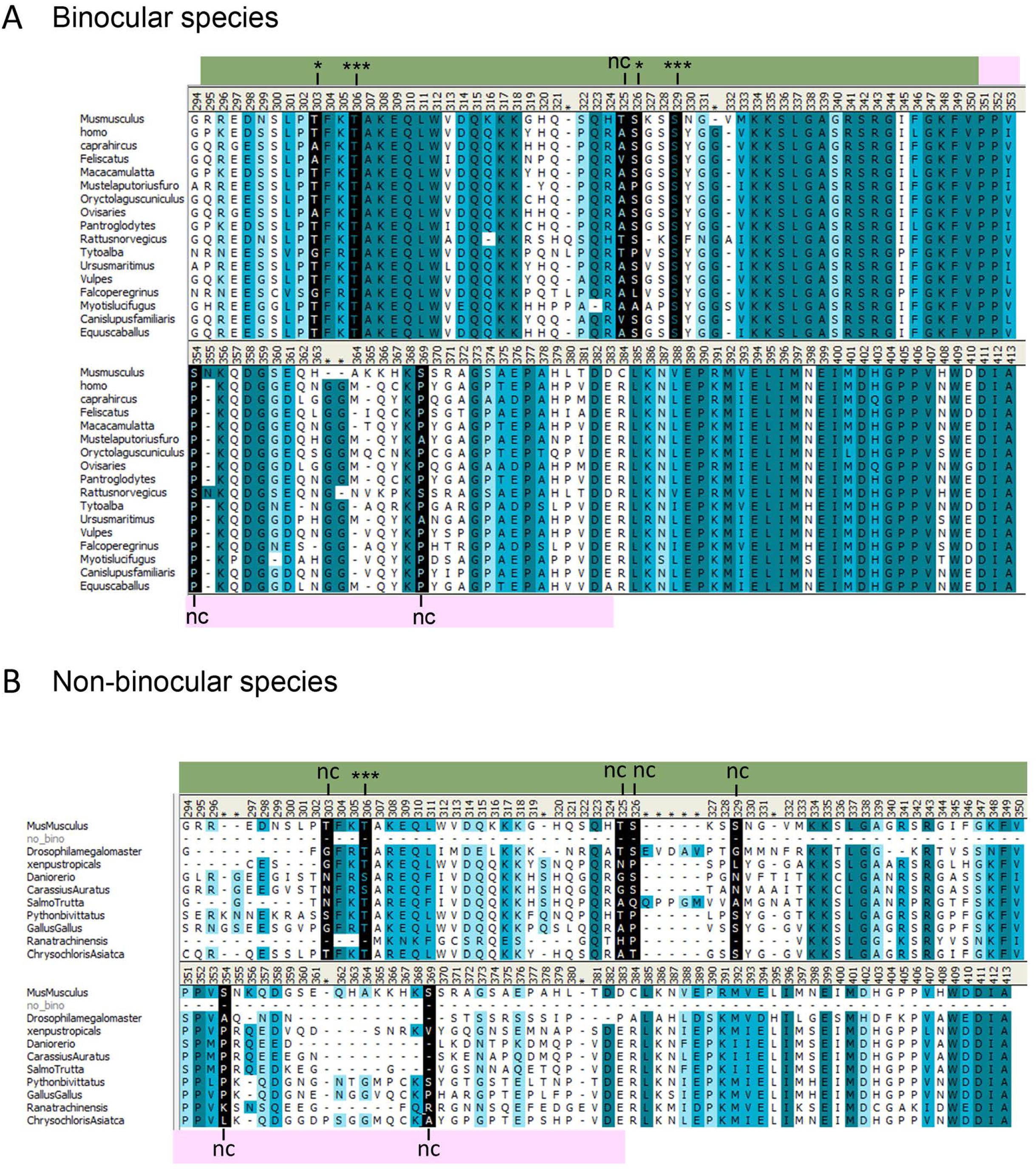
(related to Figure 6 and 7): Alignments of fignl1 sequences from binocular and non-binocular species. (A) Multiple sequence alignment of fignl1 residues 214 to 413 from seventeen binocular species. (B) Multiple sequence alignment of fignl1 residues 214 to 413 from five non-binocular species. (A-B) PKC putative phosphorylation sites are highlighted in black; (-) indicates non-conserved PKC phospho-residues. *, ***, nc, indicate poorly, highly, and non-conserved residues, respectively. PKC phospho-residues conserved across binocular species cluster within the fignl1 RBD domain (in green; residues 295 to 350) and not within the MTBD (in pink; residues 351-383). Gradually darker blue shades indicate additional evolutionarily conserved residues, with darker blue representing higher conservation.

**Figure S5.**
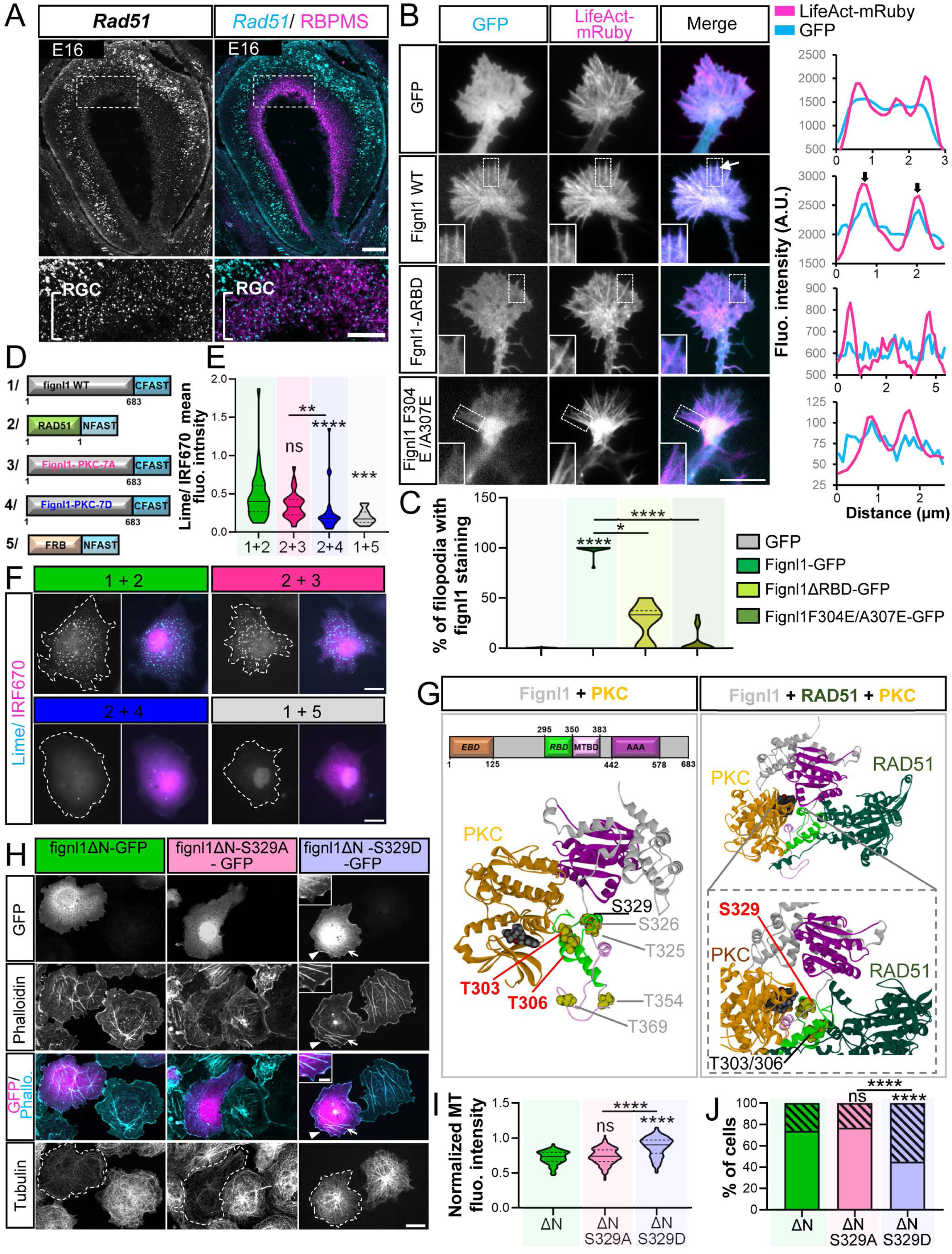
(related to figure 6): RAD51 scaffolds PKC-dependent phosphorylation of fignl1 at S329 to regulate fignl1-related cytoskeletal activity. A) SABER-FISH with *Rad51* probes (in cyan) on E16 mouse embryonic retina cryosections, post-stained with an RBPMS antibody to label RGCs (in magenta). Scale bar: 100μm. Bottom panels show higher magnifications of boxed regions. Scale bar: 50 μm. RGC: Retinal Ganglion Cell. (B) Left: Representative TIRF images of RGC growth cones from central retina explants co-electroporated with LifeAct-mRuby and fignl1-GFP, fignl1-ΔRBD-GFP, or fignl1 F304E/A307E-GFP constructs. Insets shows higher magnifications of filopodia from boxed regions. Scale bar: 10 μm. Right: Representative plot profiles of LifeAct-mRuby and fignl1-GFP, fignl1-ΔRBD-GFP, or fignl1-F304E/A307E-GFP fluorescence intensity in the P-domain of cRGC growth cones. Arrows indicate colocalisation along filopodia. (C) Violin plots showing the percentage of growth cone filopodia positive for fignl1 staining in cRGC growth cones expressing LifeAct-mRuby and GFP (n= 13), fignl1-GFP (n= 16), fignl1-ΔRBD-GFP (n= 6), or fignl1-F304E/A307E-GFP (n=17). (D) Schematic depicting the constructs used for splitFAST assays in COS-7 cells. (E) Violin plots showing splitFAST lime signal normalized to IRF670 signal (expressed from the same plasmid as fignl1-CFAST) in cells co-expressing RAD51-NFAST with fignl1-CFAST (n= 35), or fignl1-PKC-7A-CFAST (n= 32), or fignl1-GFP-PKC-7D-CFAST (n= 29), or fignl1-CFAST with FRB-NFAST (n=13). (F) Representative TIRF images of COS-7 cells transfected with different combinations of the NFAST and CFAST constructs depicted in (D) and incubated with 10 μM lime fluorogen. Scale bar: 20 μm. (G) Three-dimensional model of fignl1 (gray) in complex with PKCα (orange) (left), or with PKCα (orange) and RAD51 (dark green) (right). Candidate PKCα phosphorylation residues are highlighted in red. RAD51-binding domain, microtubule-binding domain (MTBD), and AAA cassette of Fignl1 are shown in light green, pink and purple, respectively. (H) Representative epifluorescence images of COS-7 cells overexpressing fignl1ΔN-GFP, fignl1ΔN-S329A-GFP, or fignl1ΔN-S329D-GFP, immunolabelled for GFP (magenta) and tubulin, and stained with phalloidin (cyan). Dotted lines indicate transfected cells. Arrowheads and arrows indicate fignl1ΔN-GFP colocalization with the actin cortex and actin stress fibres, respectively. Scale bar: 20μm. (I) Violin plots showing microtubule fluorescence intensity in COS-7 cells overexpressing fignl1ΔN-GFP (n=163), fignl1ΔN-S329A-GFP (n=173), or fignl1ΔN-S329D-GFP (n=183) and normalized to non-transfected cells. Cells were pooled from three independent experiments. (J) Bar graph showing the percentage of cells overexpressing fignl1ΔN (n=163), fignl1ΔN-S329A-GFP (n=173), or fignl1ΔN-S329D-GFP (n=183) exhibiting significant colocalization between fignl1 and cortical F-actin. (C, E, I-J) *p≤0.05, **p≤0.01, ***p≤0.01, ****p≤0.001. ns: non-significant. Kruskal–Wallis ANOVA with Dunn’s post hoc test (C, E, I) and Chi2 test (J). Horizontal solid and dotted lines in violin plots indicate the median ± the 1st and 3rd quartiles, respectively.

**Figure S6.**
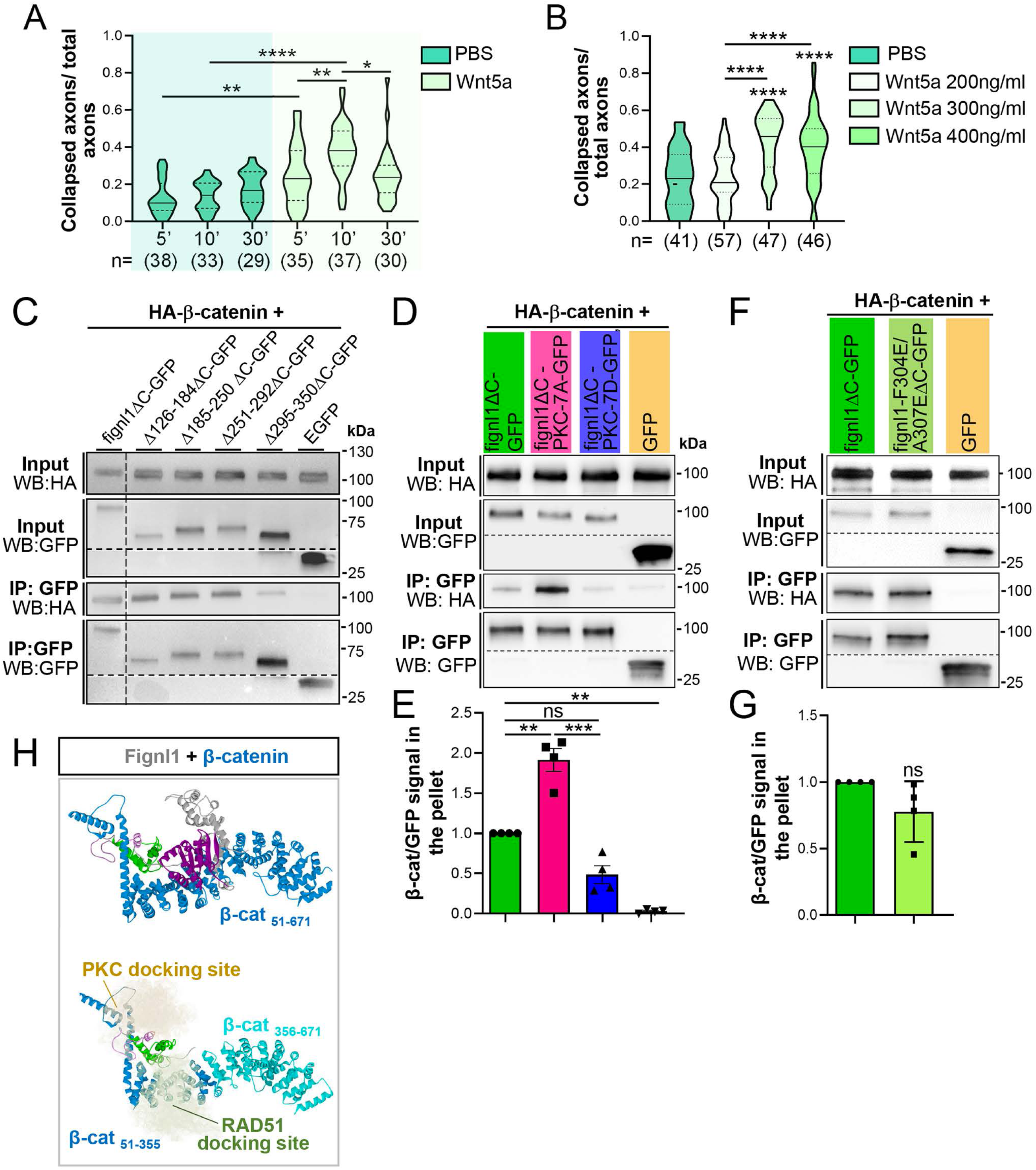
(Related to Figure 7): Competitive interaction between β-catenin and RAD51 for fignl1 binding controls the temporal sensitivity of contralateral axons to Wnt5a. (A-B) Violin plots showing the percentage of collapsed growth cones in cultures of contralateral retinal explants 5, 10, and 30 min after Wnt5a (300ng.mL^−1^) or PBS exposure (A), or after a 10 min treatment with 200, 300 or 400ng/mL Wnt5a or PBS (B). The number of explants analysed per condition, pooled from three independent experiments, is indicated below each violin plot (n=). (C) Western blot analysis of co-immunoprecipitation assays performed with a GFP antibody on protein extracts from COS-7 cells expressing HA-β-catenin together with fignl1ΔC-GFP, fignl1ΔC deletion variants, or GFP alone, and analysed using HA and GFP antibodies. (D) Western blot analysis of co-immunoprecipitation assays performed with a GFP antibody on protein extracts from COS-7 cells expressing HA-β-catenin together with fignl1ΔC-GFP, fignl1ΔC-PKC-7A-GFP, fignl1ΔC-PKC-7D, or GFP alone, and analysed using HA and GFP antibodies. (E) Bar plots showing the ratio of HA signal normalized to GFP signal in the immunoprecipitated fraction. Dots, squares, and triangles indicate individual biological replicates (n=4). For each experiment, the HA/ fignl1ΔC-GFP ratio was set to 1. (F) Western blot analysis of co-immunoprecipitation assays performed with a GFP antibody on protein extracts from COS-7 cells expressing HA-β-catenin together with fignl1ΔC-GFP, fignl1ΔC_F304E/A307E-GFP, or GFP alone, and analysed using HA and GFP antibodies. (G) Bar plots showing the ratio of HA signal normalized to GFP signal in the immunoprecipitated fraction. Dots and squares indicate individual biological replicates (n=4). For each experiment, the HA/ fignl1ΔC-GFP ratio was set to 1. (C-D, F) WB: Western blot; IP: immunoprecipitation. (H) 3-dimensional Model of fignl1 in complex with β-catenin (β-cat, blue). PKCα and RAD51 binding sites are represented by their surface volumes. Fignl1 is shown in grey, with its RBD, MTBD, and AAA domains coloured green, pink, and purple, respectively. (A-B, E-G) **p≤0.01, ***p≤0.01, **** p≤0.001. ns: non-significant. Kruskal–Wallis ANOVA with Dunn’s post hoc test (A), One-Way ANOVA with Bonferroni’s post-test (B, E), and unpaired t-test (G). Horizontal solid and dotted lines in violin plots indicate the median ± the 1st and 3rd quartiles, respectively.

**Figure S7:**
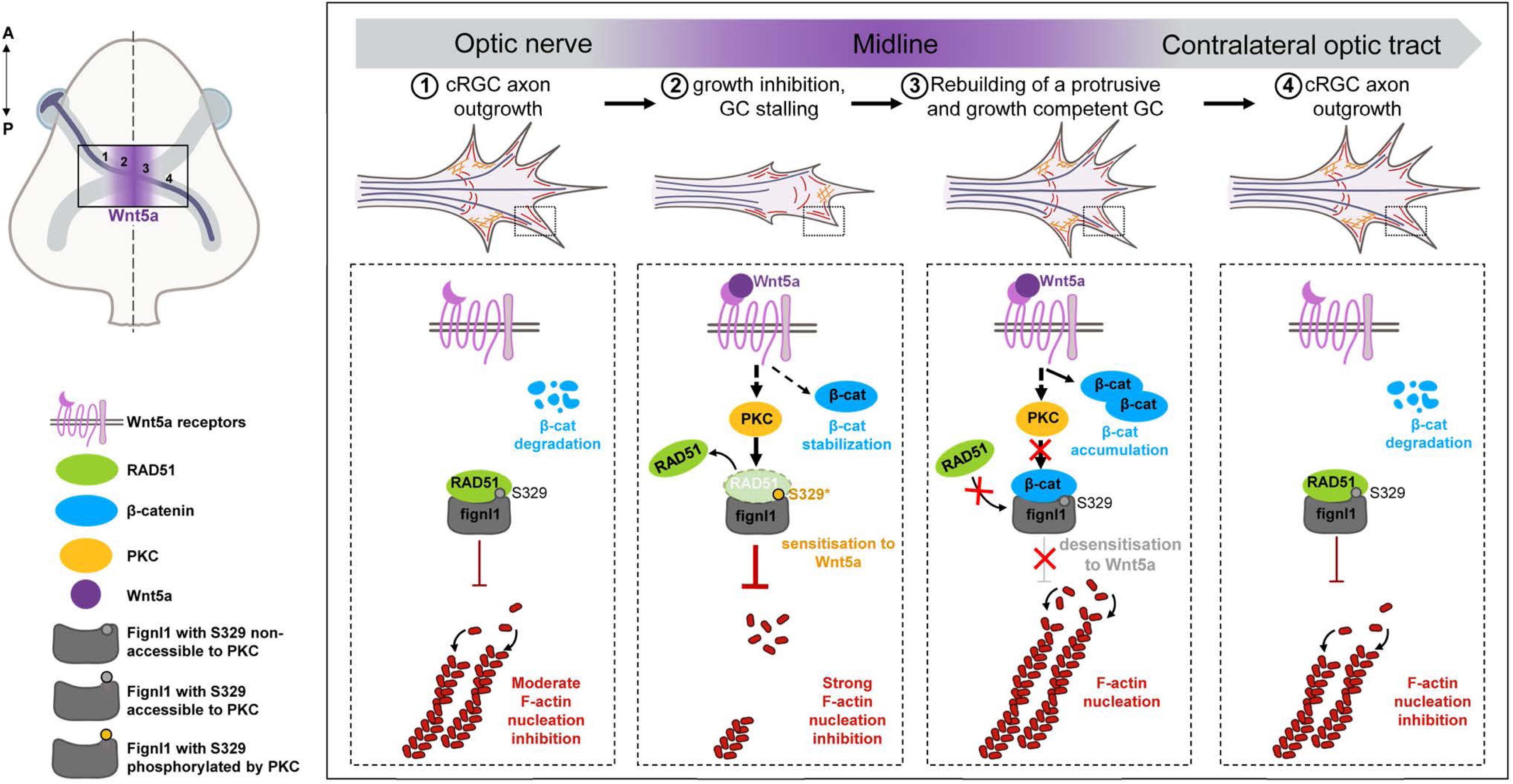
Hypothetical working model. Schematic depicting a novel key cytoskeleton-based intrinsic mechanism that controls axon midline crossing in the central nervous system. This mechanism relies on fignl1-driven inhibition of F-actin nucleation, which is (i) induced by Wnt5a at the midline, (ii) dependent on PKC-mediated phosphorylation of RAD51-scaffolded fignl1, and (iii) temporally tuned by β-catenin-mediated inhibition of RAD51 binding to fignl1, as β-catenin progressively accumulates in cRGC growth cones upon Wnt5a stimulation. This model proposes that β-catenin establishes a negative feedback loop that rapidly desensitizes cRGC axons to Wnt5a – by relieving fignl1-mediated inhibition of F-actin nucleation – allowing growth cones to exit their paused state and resume contralateral growth. GC: growth cone.

**Video S1: Fignl1 loss of function leads to an excess of ipsilateral projections, related to Figure 1**.

3D light-sheet imaging of E18.5 cleared embryo heads electroporated in the left retina with *Fignl1* scramble-(scr) or sh-RNA; tdTomato constructs and immunolabelled with anti-DsRed antibody. Arrow points at aberrant ipsilateral projections. Scale bar: 500 μm. A light-sheet, of a 4μm thickness, was generated by lasers at two different wavelengths (561nm, 639nm). Samples were scanned in LightSpeed mode with a 11% overlap and imaged using a 4X objective (N.A. 0.35) and a 4.2 Megapixel resolution sCMOS camera. TdTomato-labelled axons were segmented manually using SyGlass 2.2 software.

**Video S2: Fignl1 colocalises with F-actin-based filopodia in RGC growth cones, related to Figure 2**. Live TIRF imaging of RGC growth cones co-electroporated with fignl1-GFP, fignl1-GFP knockin (KI fignl1-GFP), or GFP control constructs together with a LifeAct-mRuby plasmid. Images were acquired every 1 sec during a 5 min time period with a 100× oil-immersion objective (N.A. 1.49) and a TIRF module. The z-drift was minimised using the Nikon Perfect Focus correction system. The incidence angles of the lasers were calibrated to obtained a 190 nm deep excitation for each wavelength. Arrows point at fignl1 co-localisation with F-actin along filopodia. Scale bar: 5 μm.

**Video S3: Fignl1 loss of function leads to the premature desensitization of contralateral RGC axons to the Wnt5a repellent cue, related to Figure 4**.

Live DIC imaging of control (Cre-^Ret^; *Fignl1^Fl/Fl^*) and fignl1-depleted (Cre+^Ret^; *Fignl1^Fl/Fl^*) contralateral RGC axons monitored 15 min before and 1 hr after exposure to PBS (Sham) or Wnt5a (300 ng.mL^−1^). Images were acquired every 15 seconds with an inverted Eclipse Ti2 (Nikon) microscope, equipped with a 40× oil-immersion objective, a Prime95B camera (Photometrics). Scale bar: 10 μm.

**Video S4: PKC-mediated phosphorylation of fignl1 S329 is required for RGC axon midline crossing, related to Figure 6**.

3D light-sheet imaging of cleared E18.5 embryo heads electroporated in the left retina with *Fignl1* sh-RNA;tdTomato and fignl1ΔN-GFP or fignl1ΔN-S329A-GFP constructs and immunolabelled with anti-DsRed and GFP antibodies. A light-sheet, of a 4μm thickness, was generated by lasers at two different wavelengths (561nm, 639nm). Samples were scanned in LightSpeed mode with a 11% overlap and imaged using a 4X objective (N.A. 0.35) and a 4.2 Megapixel resolution sCMOS camera. TdTomato- and GFP-labelled axons were segmented manually using SyGlass 2.2 software. Scale bar: 500μm.

**Video S5: Forced β-catenin accumulation in contralateral RGC axons induces their premature desensitization to Wnt5a, related to Figure 7**.

Live DIC imaging of cRGC axons expressing a degradable cytoplasmic β-catenin variant (Δ90-β-cat-ΔCT_T2A_tdTomato) or tdTomato alone and monitored 15 min before and 45 min after exposure to PBS (Sham) or Wnt5a (300 ng.mL^−1^). Images were acquired every 15 seconds with an inverted Eclipse Ti2 (Nikon) microscope, equipped with a 40× oil-immersion objective, a Prime95B camera (Photometrics). Scale bar: 10 μm.

## References

1. Chedotal, A., and Richards, L.J. (2010). Wiring the Brain: The Biology of Neuronal Guidance. Cold Spring Harbor Perspectives in Biology 2, a001917–a001917. 10.1101/cshperspect.a001917.

2. Dickson, B.J., and Zou, Y. (2010). Navigating Intermediate Targets: The Nervous System Midline. Cold Spring Harbor Perspectives in Biology 2, a002055–a002055. 10.1101/cshperspect.a002055.

3. Herrera, E., Chédotal, A., and Mason, C. (2024). Development of the Binocular Circuit. Annual Review of Neuroscience 47, 303–322. 10.1146/annurev-neuro-111020-093230.

4. Jen, J.C., Chan, W.-M., Bosley, T.M., Wan, J., Carr, J.R., Rüb, U., Shattuck, D., Salamon, G., Kudo, L.C., Ou, J., et al. (2004). Mutations in a Human ROBO Gene Disrupt Hindbrain Axon Pathway Crossing and Morphogenesis. Science 304, 1509–1513. 10.1126/science.1096437.

5. Srour, M., Rivière, J.-B., Pham, J.M.T., Dubé, M.-P., Girard, S., Morin, S., Dion, P.A., Asselin, G., Rochefort, D., Hince, P., et al. (2010). Mutations in DCC cause congenital mirror movements. Science 328, 592. 10.1126/science.1186463.

6. Izzi, L., and Charron, F. (2011). Midline axon guidance and human genetic disorders. Clinical Genetics 80, 226–234. 10.1111/j.1399-0004.2011.01735.x.

7. Nugent, A.A., Kolpak, A.L., and Engle, E.C. (2012). Human disorders of axon guidance. Curr Opin Neurobiol 22, 837. 10.1016/j.conb.2012.02.006.

8. Gallea, C., Popa, T., Hubsch, C., Valabregue, R., Brochard, V., Kundu, P., Schmitt, B., Bardinet, E., Bertasi, E., Flamand-Roze, C., et al. (2013). RAD51 deficiency disrupts the corticospinal lateralization of motor control. Brain 136, 3333–3346. 10.1093/brain/awt258.

9. Pernet, V., Joly, S., Dalkara, D., Jordi, N., Schwarz, O., Christ, F., Schaffer, D.V., Flannery, J.G., and Schwab, M.E. (2013). Long-distance axonal regeneration induced by CNTF gene transfer is impaired by axonal misguidance in the injured adult optic nerve. Neurobiology of Disease 51, 202–213. 10.1016/j.nbd.2012.11.011.

10. Luo, X., Salgueiro, Y., Beckerman, S.R., Lemmon, V.P., Tsoulfas, P., and Park, K.K. (2013). Three-dimensional evaluation of retinal ganglion cell axon regeneration and pathfinding in whole mouse tissue after injury. Experimental Neurology 247, 653–662. 10.1016/j.expneurol.2013.03.001.

11. Schaeffer, J., Vilallongue, N., Belin, S., and Nawabi, H. (2023). Axon guidance in regeneration of the mature central. Neural Regen Res 18, 2665. 10.4103/1673-5374.373663.

12. Depienne, C., Bouteiller, D., Méneret, A., Billot, S., Groppa, S., Klebe, S., Charbonnier-Beaupel, F., Corvol, J.-C., Saraiva, J.-P., Brueggemann, N., et al. (2012). RAD51 haploinsufficiency causes congenital mirror movements in humans. Am J Hum Genet 90, 301–307. 10.1016/j.ajhg.2011.12.002.

13. Bonilla, B., Hengel, S.R., Grundy, M.K., and Bernstein, K.A. (2020). RAD51 Gene Family Structure and Function. Annu. Rev. Genet. 54, 25–46. 10.1146/annurev-genet-021920-092410.

14. Shioi, T., Hatazawa, S., Takizawa, Y., and Kurumizaka, H. (2025). Mechanistic insights into RAD51-mediated nucleosome binding and remodeling in homologous recombination. DNA Repair 154, 103891. 10.1016/j.dnarep.2025.103891.

15. Tsaridou, S., and van Vugt, M.A.T.M. (2024). FIRRM and FIGNL1: partners in the regulation of homologous recombination. Trends Genet 40, 467–470. 10.1016/j.tig.2024.02.007.

16. Kumar, R. (2025). FIGNL1 hexamer dissociates RAD51-filament: a new mechanism. Trends in Biochemical Sciences 50, 287–289. 10.1016/j.tibs.2025.01.002.

17. Yuan, J., and Chen, J. (2013). FIGNL1-containing protein complex is required for efficient homologous recombination repair. Proc. Natl. Acad. Sci. U.S.A. 110, 10640–10645. 10.1073/pnas.1220662110.

18. Matsuzaki, K., Shinohara, A., and Shinohara, M. (2024). Human AAA+ ATPase FIGNL1 suppresses RAD51-mediated ultra-fine bridge formation. Nucleic Acids Research 52, 5774–5791. 10.1093/nar/gkae263.

19. Tischler, J.D., Tsuchida, H., Bosire, R., Oda, T.T., Park, A., and Adeyemi, R.O. (2024). FLIP(C1orf112)-FIGNL1 complex regulates RAD51 chromatin association to promote viability after replication stress. Nat Commun 15, 866. 10.1038/s41467-024-45139-9.

20. Kuthethur, R., Vz, S.N., Sengodan, S.K., Fonseca, C., Braunshier, S., Nagar, N., Acharya, A., Wang, X., Theil, A.F., Ibini, O., et al. (2025). FIGNL1 inhibits homologous recombination in BRCA2 deficient cells by dissociating RAD51 filaments. Science 390, eadt1210. 10.1126/science.adt1210.

21. Zainu, A., Dupaigne, P., Bouchouika, S., Cau, J., Clément, J.A.J., Auffret, P., Ropars, V., Charbonnier, J.-B., De Massy, B., Mercier, R., et al. (2024). FIGNL1-FIRRM is essential for meiotic recombination and prevents DNA damage-independent RAD51 and DMC1 loading. Nat Commun 15, 7015. 10.1038/s41467-024-51458-8.

22. Carver, A., Yu, T.-Y., Yates, L.A., White, T., Wang, R., Lister, K., Jasin, M., and Zhang, X. (2025). Molecular basis of FIGNL1 in dissociating RAD51 from DNA and chromatin. Science 387, 426–431. 10.1126/science.adr7920.

23. Cao, H., Qiu, C., Fang, A., Shang, J., Xu, W., He, L., Duan, X., Zhang, Q., and Yu, C. (2025). Extensive homologous recombination safeguards oocyte genome integrity in mammals. Nucleic Acids Research 53, gkae1304. 10.1093/nar/gkae1304.

24. Fassier, C., Fréal, A., Gasmi, L., Delphin, C., Ten Martin, D., De Gois, S., Tambalo, M., Bosc, C., Mailly, P., Revenu, C., et al. (2018). Motor axon navigation relies on Fidgetin-like 1–driven microtubule plus end dynamics. Journal of Cell Biology 217, 1719–1738. 10.1083/jcb.201604108.

25. Atkins, M., Gasmi, L., Bercier, V., Revenu, C., Del Bene, F., Hazan, J., and Fassier, C. (2019). FIGNL1 associates with KIF1Bβ and BICD1 to restrict dynein transport velocity during axon navigation. Journal of Cell Biology 218, 3290–3306. 10.1083/jcb.201805128.

26. Kidd, T., Brose, K., Mitchell, K.J., Fetter, R.D., Tessier-Lavigne, M., Goodman, C.S., and Tear, G. (1998). Roundabout Controls Axon Crossing of the CNS Midline and Defines a Novel Subfamily of Evolutionarily Conserved Guidance Receptors. Cell 92, 205–215. 10.1016/S0092-8674(00)80915-0.

27. Kidd, T., Bland, K.S., and Goodman, C.S. (1999). Slit Is the Midline Repellent for the Robo Receptor in Drosophila. Cell 96, 785–794. 10.1016/S0092-8674(00)80589-9.

28. Charron, F., Stein, E., Jeong, J., McMahon, A.P., and Tessier-Lavigne, M. (2003). The Morphogen Sonic Hedgehog Is an Axonal Chemoattractant that Collaborates with Netrin-1 in Midline Axon Guidance. Cell 113, 11–23. 10.1016/S0092-8674(03)00199-5.

29. Long, H., Sabatier, C., Le Ma, Plump, A., Yuan, W., Ornitz, D.M., Tamada, A., Murakami, F., Goodman, C.S., and Tessier-Lavigne, M. (2004). Conserved Roles for Slit and Robo Proteins in Midline Commissural Axon Guidance. Neuron 42, 213–223. 10.1016/S0896-6273(04)00179-5.

30. Yokoyama, N., Romero, MI., Cowan, C.A., Galvan, P., Helmbacher, F., Charnay, P., Parada, L.F., and Henkemeyer, M. (2001). Forward signaling mediated by ephrin-B3 prevents contralateral corticospinal axons from recrossing the spinal cord midline. Neuron 29, 85–97. 10.1016/s0896-6273(01)00182-9

31. Pourchet, O., Morel, M.-P., Welniarz, Q., Sarrazin, N., Marti, F., Heck, N., Galléa, C., Doulazmi, M., Roig Puiggros, S., Moreno-Bravo, J.A., et al. (2021). Loss of floor plate Netrin-1 impairs midline crossing of corticospinal axons and leads to mirror movements. Cell Reports 34, 108654. 10.1016/j.celrep.2020.108654.

32. Morenilla-Palao, C., López-Cascales, M.T., López-Atalaya, J.P., Baeza, D., Calvo-Díaz, L., Barco, A., and Herrera, E. (2020). A Zic2-regulated switch in a noncanonical Wnt/βcatenin pathway is essential for the formation of bilateral circuits. Sci. Adv. 6, eaaz8797. 10.1126/sciadv.aaz8797.

33. Chen, Z., Gore, B.B., Long, H., Ma, L., and Tessier-Lavigne, M. (2008). Alternative Splicing of the Robo3 Axon Guidance Receptor Governs the Midline Switch from Attraction to Repulsion. Neuron 58, 325–332. 10.1016/j.neuron.2008.02.016.

34. Colak, D., Ji, S.-J., Porse, B.T., and Jaffrey, S.R. (2013). Regulation of Axon Guidance by Compartmentalized Nonsense-Mediated mRNA Decay. Cell 153, 1252–1265. 10.1016/j.cell.2013.04.056.

35. Leggere, J.C., Saito, Y., Darnell, R.B., Tessier-Lavigne, M., Junge, H.J., and Chen, Z. (2016). NOVA regulates Dcc alternative splicing during neuronal migration and axon guidance in the spinal cord. eLife 5, e14264. 10.7554/eLife.14264.

36. Johnson, V., Junge, H.J., and Chen, Z. (2019). Temporal regulation of axonal repulsion by alternative splicing of a conserved microexon in mammalian Robo1 and Robo2. eLife 8, e46042. 10.7554/eLife.46042.

37. Nawabi, H., Briançon-Marjollet, A., Clark, C., Sanyas, I., Takamatsu, H., Okuno, T., Kumanogoh, A., Bozon, M., Takeshima, K., Yoshida, Y., et al. (2010). A midline switch of receptor processing regulates commissural axon guidance in vertebrates. Genes Dev. 24, 396–410. 10.1101/gad.542510.

38. Gorla, M., Santiago, C., Chaudhari, K., Layman, A.A.K., Oliver, P.M., and Bashaw, G.J. (2019). Ndfip Proteins Target Robo Receptors for Degradation and Allow Commissural Axons to Cross the Midline in the Developing Spinal Cord. Cell Reports 26, 3298–3312.e4. 10.1016/j.celrep.2019.02.080.

39. Keleman, K., Rajagopalan, S., Cleppien, D., Teis, D., Paiha, K., Huber, L.A., Technau, G.M., and Dickson, B.J. (2002). Comm Sorts Robo to Control Axon Guidance at the Drosophila Midline. Cell 110, 415–427. 10.1016/S0092-8674(02)00901-7.

40. Alther, T.A., Domanitskaya, E., and Stoeckli, E.T. (2016). Calsyntenin1-mediated trafficking of axon guidance receptors regulates the switch in axonal responsiveness at a choice point. Development, dev.127449. 10.1242/dev.127449.

41. Ferent, J., Giguère, F., Jolicoeur, C., Morin, S., Michaud, J.-F., Makihara, S., Yam, P.T., Cayouette, M., and Charron, F. (2019). Boc Acts via Numb as a Shh-Dependent Endocytic Platform for Ptch1 Internalization and Shh-Mediated Axon Guidance. Neuron 102, 1157–1171.e5. 10.1016/j.neuron.2019.04.003.

42. Lee, S., Nahm, M., Lee, M., Kwon, M., Kim, E., Zadeh, A.D., Cao, H., Kim, H.-J., Lee, Z.H., Oh, S.B., et al. (2007). The F-actin-microtubule crosslinker Shot is a platform for Krasavietz-mediated translational regulation of midline axon repulsion. Development 134, 1767–1777. 10.1242/dev.02842.

43. Zhuang, M., Li, X., Zhu, J., Zhang, J., Niu, F., Liang, F., Chen, M., Li, D., Han, P., and Ji, S.-J. (2019). The m6A reader YTHDF1 regulates axon guidance through translational control of Robo3.1 expression. Nucleic Acids Research 47, 4765–4777. 10.1093/nar/gkz157.

44. Lee, H., Engel, U., Rusch, J., Scherrer, S., Sheard, K., and Van Vactor, D. (2004). The Microtubule Plus End Tracking Protein Orbit/MAST/CLASP Acts Downstream of the Tyrosine Kinase Abl in Mediating Axon Guidance. Neuron 42, 913–926. 10.1016/j.neuron.2004.05.020.

45. Engel, U., Zhan, Y., Long, J.B., Boyle, S.N., Ballif, B.A., Dorey, K., Gygi, S.P., Koleske, A.J., and VanVactor, D. (2014). Abelson phosphorylation of CLASP2 modulates its association with microtubules and actin. Cytoskeleton 71, 195–209. 10.1002/cm.21164.

46. Arbeille, E., and Bashaw, G.J. (2018). Brain Tumor promotes axon growth across the midline through interactions with the microtubule stabilizing protein Apc2. PLoS Genet 14, e1007314. 10.1371/journal.pgen.1007314.

47. Sánchez-Huertas, C., and Herrera, E. (2021). With the Permission of Microtubules: An Updated Overview on Microtubule Function During Axon Pathfinding. Front. Mol. Neurosci. 14, 759404. 10.3389/fnmol.2021.759404.

48. Atkins, M., Nicol, X., and Fassier, C. (2023). Microtubule remodelling as a driving force of axon guidance and pruning. Seminars in Cell & Developmental Biology 140, 35–53. 10.1016/j.semcdb.2022.05.030.

49. Sato, S., Inoue, T., Terada, K., Matsuo, I., Aizawa, S., Tano, Y., Fujikado, T., and Furukawa, T. (2007). Dkk3-Cre BAC transgenic mouse line: a tool for highly efficient gene deletion in retinal progenitor cells. Genesis 45, 502–507. 10.1002/dvg.20318.

50. Antkowiak, A., Guillotin, A., Boiero Sanders, M., Colombo, J., Vincentelli, R., and Michelot, A. (2019). Sizes of actin networks sharing a common environment are determined by the relative rates of assembly. PLoS Biol 17, e3000317. 10.1371/journal.pbio.3000317.

51. Wang, L., Lam, J.S.-Y., Zhao, H., Wang, J., and Chan, S.-O. (2014). Localization of protein kinase C isoforms in the optic pathway of mouse embryos and their role in axon routing at the optic chiasm. Brain Research 1575, 22–32. 10.1016/j.brainres.2014.05.027.

52. Tebo, A.G., and Gautier, A. (2019). A split fluorescent reporter with rapid and reversible complementation. Nat Commun 10, 2822. 10.1038/s41467-019-10855-0.

53. Fernández-Nogales, M., López-Cascales, M.T., Murcia-Belmonte, V., Escalante, A., Fernández-Albert, J., Muñoz-Viana, R., Barco, A., and Herrera, E. (2022). Multiomic Analysis of Neurons with Divergent Projection Patterns Identifies Novel Regulators of Axon Pathfinding. Advanced Science 9, 2200615. 10.1002/advs.202200615.

54. Herrera, E., Erskine, L., and Morenilla-Palao, C. (2019). Guidance of retinal axons in mammals. Seminars in Cell & Developmental Biology 85, 48–59. 10.1016/j.semcdb.2017.11.027.

55. Prieur, D.S., and Rebsam, A. (2017). Retinal axon guidance at the midline: Chiasmatic misrouting and consequences. Developmental Neurobiology 77, 844–860. 10.1002/dneu.22473.

56. Williams, S.E., Mann, F., Erskine, L., Sakurai, T., Wei, S., Rossi, D.J., Gale, N.W., Holt, C.E., Mason, C.A., and Henkemeyer, M. (2003). Ephrin-B2 and EphB1 mediate retinal axon divergence at the optic chiasm. Neuron 39, 919–935. 10.1016/j.neuron.2003.08.017.

57. Peng, J., Fabre, P.J., Dolique, T., Swikert, S.M., Kermasson, L., Shimogori, T., and Charron, F. (2018). Sonic Hedgehog Is a Remotely Produced Cue that Controls Axon Guidance Trans-axonally at a Midline Choice Point. Neuron 97, 326–340.e4. 10.1016/j.neuron.2017.12.028.

58. Li, L., Hutchins, B.I., and Kalil, K. (2009). Wnt5a Induces Simultaneous Cortical Axon Outgrowth and Repulsive Axon Guidance through Distinct Signaling Mechanisms. J. Neurosci. 29, 5873–5883. 10.1523/JNEUROSCI.0183-09.2009.

59. Li, L., Fothergill, T., Hutchins, B.I., Dent, E.W., and Kalil, K. (2014). Wnt5a evokes cortical axon outgrowth and repulsive guidance by tau mediated reorganization of dynamic microtubules. Developmental Neurobiology 74, 797–817. 10.1002/dneu.22102.

60. Biswas, S., and Kalil, K. (2018). The Microtubule-Associated Protein Tau Mediates the Organization of Microtubules and Their Dynamic Exploration of Actin-Rich Lamellipodia and Filopodia of Cortical Growth Cones. J. Neurosci. 38, 291–307. 10.1523/JNEUROSCI.2281-17.2017.

61. McNally, F.J., and Roll-Mecak, A. (2018). Microtubule-severing enzymes: From cellular functions to molecular mechanism. Journal of Cell Biology 217, 4057–4069. 10.1083/jcb.201612104.

62. Roll-Mecak, A., and Vale, R.D. (2008). Structural basis of microtubule severing by the hereditary spastic paraplegia protein spastin. Nature 451, 363–367. 10.1038/nature06482.

63. Zehr, E., Szyk, A., Piszczek, G., Szczesna, E., Zuo, X., and Roll-Mecak, A. (2017). Katanin spiral and ring structures shed light on power stroke for microtubule severing. Nat Struct Mol Biol 24, 717–725. 10.1038/nsmb.3448.

64. Zhang, D., Rogers, G.C., Buster, D.W., and Sharp, D.J. (2007). Three microtubule severing enzymes contribute to the “Pacman-flux” machinery that moves chromosomes. The Journal of Cell Biology 177, 231–242. 10.1083/jcb.200612011.

65. Mukherjee, S., Valencia, J.D.D., Stewman, S., Metz, J., Monnier, S., Rath, U., Asenjo, A.B., Charafeddine, R.A., Sosa, H.J., Ross, J.L., et al. (2012). Human Fidgetin is a microtubule severing the enzyme and minus-end depolymerase that regulates mitosis. Cell Cycle 11, 2359–2366. 10.4161/cc.20849.

66. Kessels, M.M., Schwintzer, L., Schlobinski, D., and Qualmann, B. (2011). Controlling actin cytoskeletal organization and dynamics during neuronal morphogenesis. European Journal of Cell Biology 90, 926–933. 10.1016/j.ejcb.2010.08.011.

67. Gomez, T.M., and Letourneau, P.C. (2014). Actin dynamics in growth cone motility and navigation. Journal of Neurochemistry 129, 221–234. 10.1111/jnc.12506.

68. Schneider, F., Metz, I., and Rust, M.B. (2023). Regulation of actin filament assembly and disassembly in growth cone motility and axon guidance. Brain Research Bulletin 192, 21–35. 10.1016/j.brainresbull.2022.10.019.

69. Paunola, E., Mattila, P.K., and Lappalainen, P. (2002). WH2 domain: a small, versatile adapter for actin monomers. FEBS Letters 513, 92–97. 10.1016/S0014-5793(01)03242-2.

70. Dent, E.W., Kwiatkowski, A.V., Mebane, L.M., Philippar, U., Barzik, M., Rubinson, D.A., Gupton, S., Van Veen, J.E., Furman, C., Zhang, J., et al. (2007). Filopodia are required for cortical neurite initiation. Nat Cell Biol 9, 1347–1359. 10.1038/ncb1654.

71. Edwards, M., Zwolak, A., Schafer, D.A., Sept, D., Dominguez, R., and Cooper, J.A. (2014). Capping protein regulators fine-tune actin assembly dynamics. Nat Rev Mol Cell Biol 15, 677–689. 10.1038/nrm3869.

72. Evans, K.J., Gomes, E.R., Reisenweber, S.M., Gundersen, G.G., and Lauring, B.P. (2005). Linking axonal degeneration to microtubule remodeling by Spastin-mediated microtubule severing. The Journal of Cell Biology 168, 599–606. 10.1083/jcb.200409058.

73. Kuo, Y.-W., Trottier, O., Mahamdeh, M., and Howard, J. (2019). Spastin is a dual-function enzyme that severs microtubules and promotes their regrowth to increase the number and mass of microtubules. Proc Natl Acad Sci U S A 116, 5533–5541. 10.1073/pnas.1818824116.

74. Liu, H., Matsumoto, N., Kaneshiro, T., Kato, T., Shigematsu, H., Kikkawa, S., Nitta, E., Imasaki, T., and Nitta, R. (2025). Human spastin functions as an ATPase-independent microtubule nucleator via ring-stack formation. Preprint at Biophysics, 10.1101/2025.11.28.691112 10.1101/2025.11.28.691112.

75. Caridi, C.P., Plessner, M., Grosse, R., and Chiolo, I. (2019). Nuclear actin filaments in DNA repair dynamics. Nat Cell Biol 21, 1068–1077. 10.1038/s41556-019-0379-1.

76. Schrank, B.R., Aparicio, T., Li, Y., Chang, W., Chait, B.T., Gundersen, G.G., Gottesman, M.E., and Gautier, J. (2018). Nuclear ARP2/3 drives DNA break clustering for homology-directed repair. Nature 559, 61–66. 10.1038/s41586-018-0237-5.

77. Caridi, C.P., D’Agostino, C., Ryu, T., Zapotoczny, G., Delabaere, L., Li, X., Khodaverdian, V.Y., Amaral, N., Lin, E., Rau, A.R., et al. (2018). Nuclear F-actin and myosins drive relocalization of heterochromatic breaks. Nature 559, 54–60. 10.1038/s41586-018-0242-8.

78. Schlienger, S., Yam, P.T., Balekoglu, N., Ducuing, H., Michaud, J.-F., Makihara, S., Kramer, D.K., Chen, B., Fasano, A., Berardelli, A., et al. (2023). Genetics of mirror movements identifies a multifunctional complex required for Netrin-1 guidance and lateralization of motor control. Science Advances 9, eadd5501. 10.1126/sciadv.add5501.

79. Chaudhari, K., Zhang, K., Yam, P.T., Zang, Y., Kramer, D.A., Gagnon, S., Schlienger, S., Calabretta, S., Michaud, J.-F., Collins, M., et al. (2024). A human DCC variant causing mirror movement disorder reveals that the WAVE regulatory complex mediates axon guidance by netrin-1–DCC. Science Signaling 17, eadk2345. 10.1126/scisignal.adk2345.

80. Rogers, S., and Scholpp, S. (2022). Vertebrate Wnt5a – At the crossroads of cellular signalling. Seminars in Cell & Developmental Biology 125, 3–10. 10.1016/j.semcdb.2021.10.002.

81. Weeraratna, A.T., Jiang, Y., Hostetter, G., Rosenblatt, K., Duray, P., Bittner, M., and Trent, J.M. (2002). Wnt5a signaling directly affects cell motility and invasion of metastatic melanoma. Cancer Cell 1, 279–288. 10.1016/S1535-6108(02)00045-4.

82. Kurayoshi, M., Oue, N., Yamamoto, H., Kishida, M., Inoue, A., Asahara, T., Yasui, W., and Kikuchi, A. (2006). Expression of Wnt-5a Is Correlated with Aggressiveness of Gastric Cancer by Stimulating Cell Migration and Invasion. Cancer Research 66, 10439–10448. 10.1158/0008-5472.CAN-06-2359.

83. Asem, M., Buechler, S., Wates, R., Miller, D., and Stack, M. (2016). Wnt5a Signaling in Cancer. Cancers 8, 79. 10.3390/cancers8090079.

84. Hershberg, E.A., Camplisson, C.K., Close, J.L., Attar, S., Chern, R., Liu, Y., Akilesh, S., Nicovich, P.R., and Beliveau, B.J. (2021). PaintSHOP enables the interactive design of transcriptome- and genome-scale oligonucleotide FISH experiments. Nat Methods 18, 937–944. 10.1038/s41592-021-01187-3.

85. Attar, S., Browning, V.E., Krebs, M., Liu, Y., Nichols, E.K., Tsue, A.F., Shechner, D.M., Shendure, J., Lieberman, J.A., Schweppe, D.K., et al. (2025). Efficient and highly amplified imaging of nucleic acid targets in cellular and histopathological samples with pSABER. Nat Methods 22, 156–165. 10.1038/s41592-024-02512-2.

86. Baudet, S., Zagar, Y., Roche, F., Gomez-Bravo, C., Couvet, S., Bécret, J., Belle, M., Vougny, J., Uthayasuthan, S., Ros, O., et al. (2023). Subcellular second messenger networks drive distinct repellent-induced axon behaviors. Nat Commun 14, 3809. 10.1038/s41467-023-39516-z.

87. Garcia-Frigola, C., Carreres, M.I., Vegar, C., and Herrera, E. (2007). Gene delivery into mouse retinal ganglion cells by in utero electroporation. BMC Dev Biol 7, 103. 10.1186/1471-213X-7-103.

88. Petros, T.J., Rebsam, A., and Mason, C.A. (2009). In utero and ex vivo Electroporation for Gene Expression in Mouse Retinal Ganglion Cells. JoVE, 1333. 10.3791/1333.

89. Belle, M., Godefroy, D., Couly, G., Malone, S.A., Collier, F., Giacobini, P., and Chédotal, A. (2017). Tridimensional Visualization and Analysis of Early Human Development. Cell 169, 161–173.e12. 10.1016/j.cell.2017.03.008.

90. Sharma, A., Aher, A., Dynes, N.J., Frey, D., Katrukha, E.A., Jaussi, R., Grigoriev, I., Croisier, M., Kammerer, R.A., Akhmanova, A., et al. (2016). Centriolar CPAP/SAS-4 Imparts Slow Processive Microtubule Growth. Dev Cell 37, 362–376. 10.1016/j.devcel.2016.04.024.

91. Jégou, A., Carlier, M.-F., and Romet-Lemonne, G. (2013). Formin mDia1 senses and generates mechanical forces on actin filaments. Nat Commun 4, 1883. 10.1038/ncomms2888.

92. Elie, A., Prezel, E., Guérin, C., Denarier, E., Ramirez-Rios, S., Serre, L., Andrieux, A., Fourest-Lieuvin, A., Blanchoin, L., and Arnal, I. (2015). Tau co-organizes dynamic microtubule and actin networks. Sci Rep 5, 9964. 10.1038/srep09964.

93. Blom, N., Gammeltoft, S., and Brunak, S. (1999). Sequence- and structure-based prediction of eukaryotic protein phosphorylation sites. J Mol Biol 294, 1351. 10.1006/jmbi.1999.3310.

94. El Hajji, L., Bunel, B., Joliot, O., Li, C., Tebo, A.G., Rampon, C., Volovitch, M., Fischer, E., Pietrancosta, N., Perez, F., et al. (2025). A tunable and versatile chemogenetic near-infrared fluorescent reporter. Nat Commun 16, 2594. 10.1038/s41467-025-58017-9.

95. Šali, A., and Blundell, T.L. (1993). Comparative protein modelling by satisfaction of spatial restraints. J Mol Biol 234, 779. 10.1006/jmbi.1993.1626.

96. Jumper, J., Evans, R., Pritzel, A., Green, T., Figurnov, M., Ronneberger, O., Tunyasuvunakool, K., Bates, R., Žídek, A., Potapenko, A., et al. (2021). Highly accurate protein structure prediction with AlphaFold. Nature 596, 583. 10.1038/s41586-021-03819-2.

97. Phillips, J.C., Braun, R., Wang, W., Gumbart, J., Tajkhorshid, E., Villa, E., Chipot, C., Skeel, R.D., Kalé, L., and Schulten, K. (2005). Scalable molecular dynamics with NAMD. J Comput Chem 26, 1781. 10.1002/jcc.20289.

98. Chen, R., Li, L., and Weng, Z. (2003). ZDOCK: an initial-stage protein docking algorithm. Proteins 52, 80. 10.1002/prot.10389.

